# Studies of a siderophore-producing cyclization domain: A refined proposal of substrate binding

**DOI:** 10.1101/2022.03.18.483834

**Authors:** Andrew D. Gnann, Yuan Xia, Jess Soule, Clara Barthélemy, Jayata Mawani, Sarah Nzikoba Musoke, Brian Castellano, Edward Brignole, Dominique Frueh, Daniel P. Dowling

## Abstract

Nonribosomal peptide synthetase (NRPS) heterocyclization (Cy) domains generate biologically important ox-/thiazoline modifications in natural products, including in production of compounds targeting disease or siderophores that are important for bacterial pathogenicity. Cy domains share the NRPS condensation domain fold but catalyze consecutive condensation and cyclodehydration reactions via an unknown mechanism. To further understanding of Cy domain catalysis, we report the crystal structure of the second Cy domain (Cy2) of yersiniabactin synthetase from the causative agent of the plague, *Yersinia pestis*. We find the high-resolution structure of Cy2 adopts a conformation enabling exploration of binding the extended, thiazoline-containing cyclodehydration intermediate for catalysis and the acceptor carrier protein to which it is tethered. We also report complementary electrostatic interfaces between Cy2 and its donor carrier protein that mediate donor binding. Lastly, we explore domain flexibility through the normal mode approximation and identify small-molecule fragment binding sites to inform antibiotic design targeting Cy function. Our results suggest how carrier protein binding may influence global conformation, with consequences for active site catalytic states and inhibitor development.

## Introduction

A vast and growing variety of nonribosomal peptide (NRP) and polyketide (PK) natural products is produced by modular, multidomain polyketide synthase (PKS) or nonribosomal peptide synthetase (NRPS) systems^1–4^. NRP, PK, and hybrid NRP/PK natural products have widely ranging bioactivities with clear importance to humans^5–8^, and the modularity of NRPS and PKS systems makes them prime candidates for bioengineering^9^. Additionally, NRP and PK secondary metabolites are primarily produced in bacteria and, to a lesser extent, fungi, and some are virulence factors of pathogenic bacteria^10–18^; therefore, a thorough understanding and control of NRPS and PKS catalytic machinery may afford new options for antibiotic targets^19^.

NRPS and PKS systems follow an assembly line logic in which each module is responsible for activating and incorporating a specific building block. Canonical NRPS elongation modules consist of adenylation (A), condensation (C), and peptidyl carrier protein (PCP) domains. A domains activate and load specific amino acids onto PCP phosphopantetheine (Ppant) cofactors as thioesters. PCP domains from different modules (herein called upstream/ donor and downstream/acceptor PCPs) shuttle their thioester-tethered substrates/intermediates to the C domain of one of the modules, which catalyzes their condensation to yield the elongated, peptide product attached to the acceptor PCP. The product may then act as a donor intermediate in a subsequent condensation within the next module or be released hydrolytically within the final module. Substrates and intermediates are often also modified by tailoring domains, giving rise to various oxidation states^20–23^, methylation patterns^24–29^ and other alterations^30–33^. Of particular interest for the present work, many NRPSs contain a variant of C domains, the heterocyclization (Cy) domain, that converts hydroxyl- or thiol-containing peptides into heterocycles during chain elongation. For more information on the C domain family, see Bloudoff and Schmeing’s 2017 review^34^.

Thiazoline and (methyl)oxazoline rings formed by Cy domains are prevalent and important for the bioactivity of many natural products (Fig. S1), a prime example being thiazol(in)e-containing siderophores such as yersiniabactin (Ybt)^35^. Ybt is a virulence factor produced by *Yersinia* bacteria, the causative agents of the plague. Ybt is produced under iron-limiting conditions to scavenge necessary trace metal ions from the extracellular environment, including Fe^3+^ from the host^36,37^, and the Fe^3+^-Ybt complex^37^ is delivered to the cell via a TonB-dependent outer membrane receptor and an inner membrane ABC transporter^38^. Since yersiniabactin is just one of a number of shikimate-derived, thiazole/oxazole siderophores that are required for growth of some bacteria in iron-restricted environments^3,12,15,39,40^, understanding the crucial Cy domain activities in its biosynthesis may support development of a new class of antibiotics against these bacteria^41–43^. It is interesting to note that Cy domain sequences diverge considerably, with Cy domains in the same biosynthetic gene cluster sharing only ~20% sequence identity in some instances (such as anguibactin synthetase’s AngN-Cy1 and -Cy2 or vibriobactin synthetase’s VibF-Cy1 and -Cy2)^44^, and the three Cy domains of the high-molecular weight proteins in *Yersinia pestis* (HMWP1 and HMWP2) share only ~35% sequence identity. This sequence diversity might permit development of selective inhibitors against Cy domains of pathogenic species, avoiding endogenous Cy domains of the human microbiome^45^.

The biosynthesis of Ybt provides an example of a noncanonical NRPS/PKS system that entails the activity of seven proteins encoded within a pathogenicity island^25,33,46–48^: YbtS, YbtD, YbtE, HMWP1, HMWP2, YbtU, and YbtT (Fig. 1). HMWP2 contains the starter module of the pathway and has the domain order ArCP-Cy1-A(E)-PCP1-Cy2-PCP2, where ArCP is an aryl carrier protein and A(E) is an interrupted adenylation domain embedding an unconventional 37 kDa epimerase domain^28,33^. HMWP1 contains Cy3 and PCP3 in a hybrid PKS/NRPS context. The CPs are phosphopantetheinylated by YbtD, the ArCP is loaded with a salicyl thioester by the standalone A domain YbtE, and the A(E) domain of HMWP2 is responsible for loading l-cysteine onto PCP1 and PCP2 (*in cis*) and PCP3 (*in trans).* The embedded E domain generates the d stereoisomer of the first thiazoline installed by HMWP2-Cy1. Since Ybt only contains one chirality at that position, despite incomplete conversion by the E domain, the second cyclization domain, HMWP2-Cy2, is expected to provide stereospecificity. YbtU is an NADPH-dependent reductase responsible for conversion of the second heterocycle of Ybt to thiazolidine. Lastly, it is thought that the type II thioesterase YbtT may hydrolytically cleave off aberrantly loaded substrates^49^. The product of HMWP2-Cy2, phosphopantetheine-tethered 2-hydroxyphenylthiazolinylthiazolidine, is passed at an interprotein NRPS/PKS junction between HMWP2 and HMWP1. Both NRPS and PKS systems require movements of CPs to transfer elongating molecules (Fig. 1), however our understanding of NRPS/PKS interfaces outside of docking domain studies is limited^50^. Therefore, the rich, noncanonical architecture of Ybt synthetase is of great interest for exploring structure-function relationships in hybrid NRPS/PKS systems. As a first step toward modeling the NRPS to PKS interface between HMWP2 and HMWP1, we explore the HMWP2-Cy2 domain, which with its acceptor PCP (HMWP2-PCP2) forms the NRPS half of the NRPS/ PKS interface.

**Figure 1.**
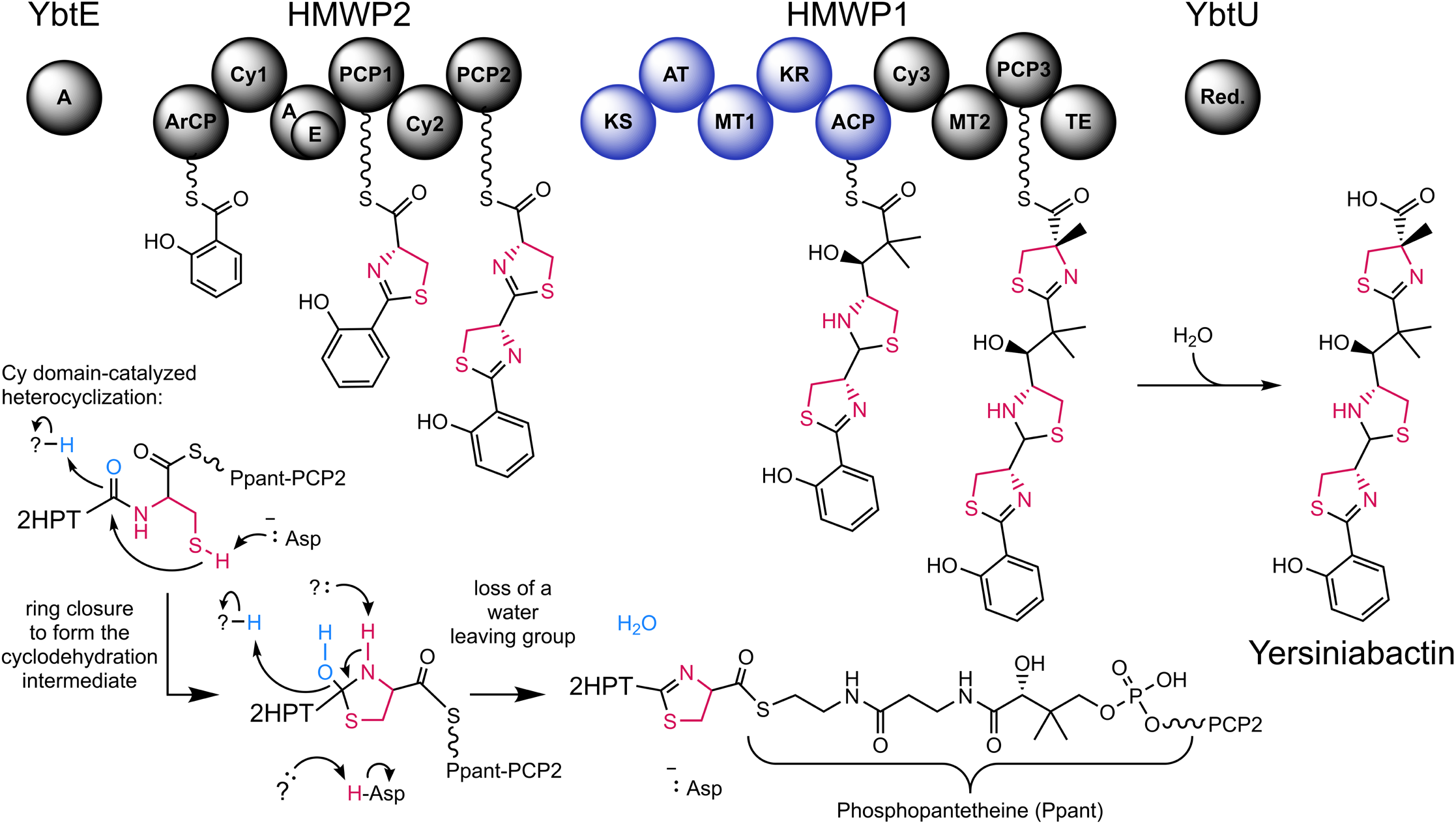
The four protein, hybrid NRPS-PKS, modular assembly line for biosynthesis of yersiniabactin and a mechanistic proposal for HMWP2-Cy2-catalyzed heterocyclization. Each circle represents either a NRPS domain (black) or a PKS domain (blue). Bond-line structures are substrates/products/intermediates carried by their respective carrier proteins. Domain labeling (left to right), and descriptions for PKS domains not detailed in the main text, are: A – Adenylation; ArCP – the Aryl Carrier Protein; Cy – Heterocyclization; E – Epimerization; PCP – Peptidyl Carrier Proteins; KS – Ketosynthase, catalyzes Claisen condensation of acyl donor and malonyl acceptor groups; AT – Acyl Transferase, catalyzes tethering of malonyl units onto the acyl carrier protein (ACP); MT – Methyltransferase, catalyzes methyl transfer from S-adenosyl methionine to cysteinyl or malonyl C2 positions; KR – Ketoreductase, catalyzes NAD(P)H-dependent reduction of the keto group from the acyl donor substrate of the KS; ACP – Acyl Carrier Protein; TE –Thioesterase, catalyzes hydrolytic cleavage of the product thioester; Red. – Reductase. (Not shown: YbtS – synthesizes input salicylate from chorismate; YbtD – transfers phosphopantetheine to CPs; YbtT – type II thioesterase putatively hydrolyzing improperly loaded substrates). The phosphopantetheine (Ppant) cofactor is represented as a wavy line-S and is drawn out in the bottom right of the figure. Atoms in light blue in the partial mechanism in the bottom left constitute the water leaving group. Thiazoline heterocycle atoms derived from cysteine are colored magenta throughout the figure.

NRPS Cy domains are typically responsible for both condensation and cyclodehydration reactions, affording (methyl)ox-/thiazoline modifications (Figs. 1 and S2), and their catalytic mechanisms are only partially understood. Five structures of Cy domains have been reported, involved in biosynthesis of epothilones^51^, bacillamide^52^, yersiniabactin^53^, JBIR-34/35^54^ and pyochelin^55^. The first two Cy domain structures identified a structural role for aspartates of the characteristic DXXXXD motif in the crystallized state, which was in keeping with observed roles for residues at these positions in crystallized C domains^34,51,52^. The Cy domain’s active site, formed in a tunnel between its N- and C-terminal subdomains as in C domains, was found to contain a putative catalytic aspartate-threonine dyad^51,52^ as well as a nearby glutamine^51^, mutations of which severely affected cyclodehydration. Interestingly, the glutamine is not present in BmdB-Cy2, suggesting complementation of its yet unknown role by some other means in that system. Several additional conserved residues have been shown to be important for the efficiency of catalysis but also serve unknown roles. Notably, HMWP2’s Cy1 domain has been shown to display global dynamics, wherein a mutation leads to a global molecular response^53^, highlighting the limitations of traditional structural studies relying on a single, static structure and the need to identify conformations critical for function. Finally, there is limited structural information for binding of (amino)acyl/peptidyl-Ppant in a Cy domain or for binding carrier proteins (CPs) in catalytically competent states, with the recent cryoEM structures of FmoA3^54^ and PchE^55^ providing the first snapshots of CP binding to Cy domains within modular contexts. Overall, more structural snapshots of Cy domains are needed to resolve important gaps in understanding how they function.

Here we report the 1.94 Å-resolution crystal structure of the second Cy domain of Ybt synthetase (HMWP2-Cy2), which with its acceptor PCP (HMWP2-PCP2) forms the NRPS half of the NRPS/PKS interface. In addition to Ybt’s role in pathogenesis for *Y. pestis*, this cyclization domain is of further interest because it accepts a donor that already features a heterocycle and because Cy2 is expected to govern the stereochemistry of the end product, Ybt^33^. We construct protein-protein docking models suggesting interactions in the tridomain complex HMWP2-PCP1-Cy2-PCP2, identifying likely protein contacts contributing to HMWP2’s synthetic directionality at the second cyclization domain. We then take advantage of the less constricted active site tunnel of the HMWP2-Cy2 crystal structure to report covalent ligand docking experiments tethering potential cyclodehydration intermediates to the conserved PCP serine from the PCP_acceptor_ protein-protein docking result. Alongside comparisons to previous C and Cy domain structures, the results motivate consideration of a limited set of possibilities for the positioning of an extended cyclodehydration intermediate. By considering knowledge of conformational flexibility in the C domain family and approximating HMWP2-Cy2’s flexibility in the normal mode approximation, consequences for positioning of the condensation product for cyclodehydration are also discussed. The Cy conformation we present here provides important clues on communication between Cy domains and their substrates and partner CP domains, suggesting relay mechanisms relying on structural fluctuations that may couple domain engagements with active site remodeling and global remodeling of the entire Cy domain.

## Results

### Structure of a Cy domain from siderophore biosynthesis

The crystal structure of HMWP2-Cy2 was solved to 1.94 Å resolution by molecular replacement using the structure of the Cy domain from epothilone biosynthesis^51^ (EpoB-Cy, 35% sequence identity) as a search model, and the subsequent 2.35 Å-resolution structure of HMWP2-Cy2 was solved using the 1.94 Å-resolution HMWP2-Cy2 model (Table S1). These structures demonstrate the pseudodimeric fold characteristic of the condensation domain superfamily, which is a two-lobed arrangement of two chloramphenicol acetyltransferase (CAT)-like folds (Fig. 2A and Fig. S3). The CAT-like folds, called the N- and C-terminal subdomains here, each contain a mixed β sheet with intervening α helices. The C-terminal β sheet is formed by strands 7, 13, 12, 10, 8 and 9 (in order), and the N-terminal β sheet is formed by strands 4, 5, 6, 1 and 11, with strand 11 donated from the C-terminal subdomain. Strand 11 and the preceding loop have been referred to as a latch and are thought to mediate partial dissociation of the subdomains (Fig. 2A)^56^. The two points of crossover between the subdomains are the predominantly loop region between β-strands 8 and 9 in the C-terminal subdomain, often referred to as the floor loop, and the latch.

**Figure 2.**
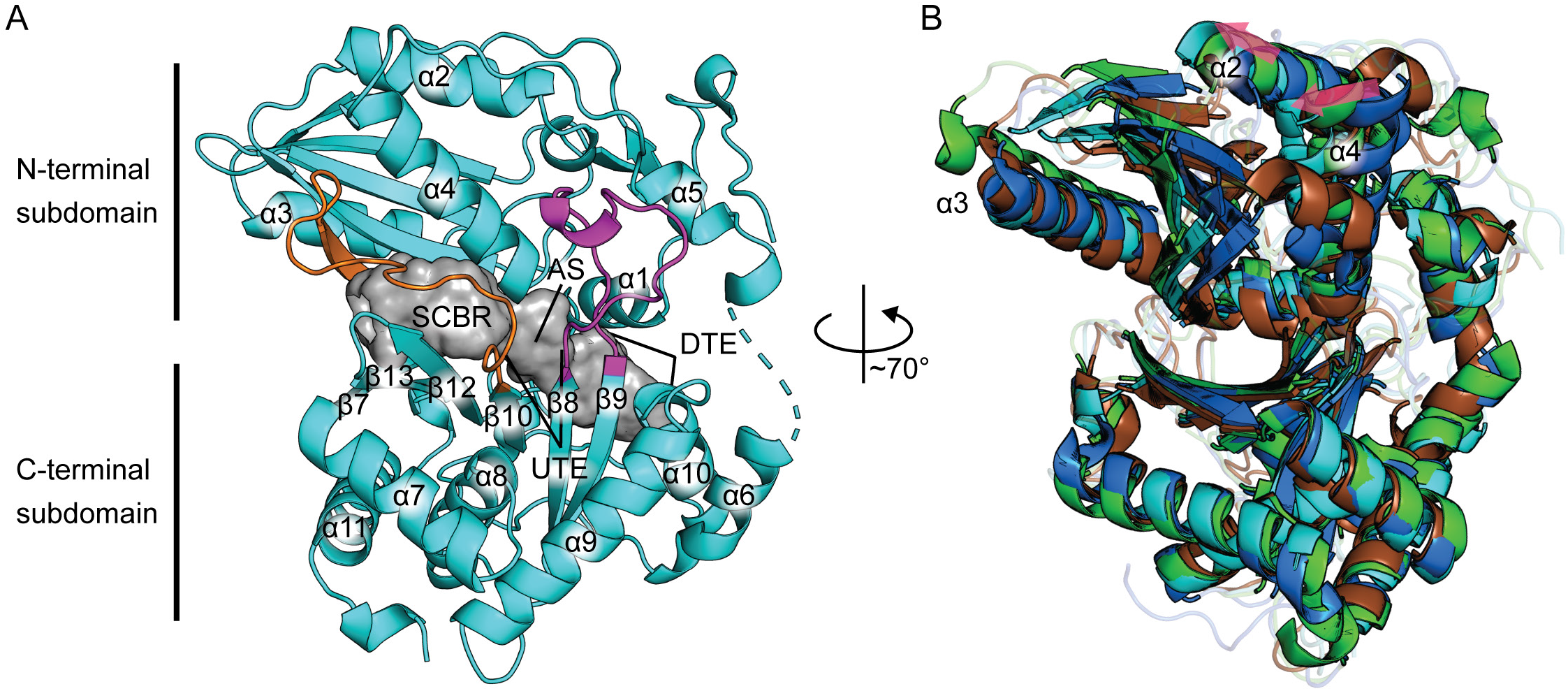
The HMWP2-Cy2 crystal structure exhibits a more open active site tunnel. **A**, an overview of the 1.94 Å-resolution HMWP2-Cy2 crystal structure with the tunnel as a gray surface, the floor loop and latch in light magenta and orange cartoon, respectively, and other features discussed in the text labeled (SCBR – side chain-binding region, AS – active site, UTE – upstream tunnel entrance, DTE - downstream tunnel entrance). **B**, Cy domain C-terminal subdomain Cα alignment demonstrates intersubdomain conformational variation among Cy domains. HMWP2-Cy2 is cyan, EpoB-Cy is green, BmdB-Cy2 is blue, and PchE-Cy is brown. Loop regions are transparent. Pink arrows highlight correlated variation between the models.

HMWP2-Cy2 crystallizes in the space group *P*4_1_2_1_2 with one molecule in the asymmetric unit. The largest interface area between Cy2 molecules within the crystal buries ~680 Å^2^ (3.6%) of the solvent accessible surface area, based on analysis using the PISA server^57^; therefore, HMWP2-Cy2 is likely a monomer in solution, consistent with the domain’s behavior by size exclusion chromatography (Fig. S4). Electron density maps obtained using the 1.94 Å-resolution data permitted building residues Q1483-L1663 and N1669-Q1910 (HMWP2 numbering). The structure solved at 2.35 Å resolution includes weak electron density for the missing residues except P1666, which remains disordered, allowing for generation of a composite model using coordinates from the higher-resolution data set for all atoms other than residues L1663-P1668 and residues 1663-1665, 1667 and 1668 from the 2.35 Å-resolution structure. P1666 was built and refined to reduce potential energy in the final composite model and achieve acceptable connectivity.

Of the prior Cy domain structures, the Cy domain from EpoB^51^ most closely resembles HMWP2-Cy2 (root-mean-square deviation, RMSD, of 1.4 Å between 338 out of 431 Cα atoms, aligning against PDB ID 5T7Z). Interestingly, protein alignments improve when considering either the N-terminal subdomain (residues 1483 to 1665) or the C-terminal subdomain (residues 1666 to 1910) independently, consistent with movement of these subdomains relative to each other within published NRPS Cy domain structures (Fig. 2B)^51–55^. Specifically, in a Cα alignment of the C-terminal subdomains of HMWP2-Cy2 and EpoB-Cy, helices α2 and α4 of HMWP2-Cy2’s N-terminal subdomain are shifted in the direction of the N-terminal subdomain β sheet, and helix α3 of HMWP2-Cy2 is shifted toward the latch strand β11. This pattern of motion also appears to involve twisting of the N-terminal subdomain β sheet, resulting in strands β4 and β5 shifting in the direction of helix α3. In a similar alignment of HMWP2-Cy2 with BmdB-Cy2^52^, helices α2 and α4 of BmdB-Cy2 are found on the opposite side (relative to HMWP2-Cy2) of their position in the EpoB-Cy-to-HMWP2-Cy2 alignment—further in the direction of the floor loop. Accompanying this shift, helix α3 and the N-terminal subdomain β sheet of BmdB-Cy2 are also shifted toward the floor loop so that α3 nearly coincides with HMWP2-Cy2’s α3. With respect to these shifts in secondary structure features, the recent cryoEM structures of PchE-Cy1^55^ are overall highly similar to the HMWP2-Cy2 conformation.

The conformational differences between HMWP2-Cy2 and the prior Cy domain crystal structures contribute toward a more open, continuous tunnel in HMWP2-Cy2, with a solvent accessible surface volume of ~543 Å^3^ (Fig. 2 and Fig. S5). This relatively straight tunnel has been observed previously in EpoB-Cy and BmdB-Cy2 as well as in members of the condensation domain family^51,52,58,59^, however the tunnels of EpoB-Cy and BmdB-Cy2 are more constricted (Fig. S5) with the BmdB-Cy2 tunnel returning a volume of ~382 Å^3^ and EpoB-Cy’s appearing as three chambers, separated by bottlenecks narrower than the diameter of water, collectively measuring ~199 Å^3^. In particular, the EpoB-Cy and BmdB-Cy2 structures are constricted in the vicinity of the donor substrate side chain-binding region between helix α4, the latch strand, and β-strands 7, 13, 12, and 10 of the C-terminal subdomain sheet (Fig. 2A). Interestingly, the upstream tunnel entrance is closed to donor substrate binding in the Cy crystal structures, and the downstream tunnel entrance is open. The conformations seen in these crystallographic studies resemble those captured by a solution-NMR structure bundle recently determined for HMWP2’s Cy1^53^, suggesting that the domains adopt these conformations in solution. The entrance to the upstream tunnel is typically more open in the NMR bundle than in the HMWP2-Cy2 structure, and the downstream entrance more closed in some Cy1 states, and both regions display relaxation dispersion and hence malleability. CryoEM structures of PchE similarly report both closed and open conformations for the upstream tunnel entrance^55^.

The active site residues D1862, T1830, and Q1864 are found in conformations resembling EpoB-Cy^51^ and BmdB-Cy2^52^, excepting the replacement of the equivalent position of HMWP2 Q1864 with valine in BmdB-Cy2. Intriguingly, the HMWP2-Cy2 active site contains an unexpected metal-binding site against β1 and the latch strand β11. This metal site involves three backbone carbonyl groups (S1520, H1521 and Q1855), a carboxamide (N1621 from position two of the DXXXXD motif), and two crystallographic waters, giving rise to a distorted octahedral coordination geometry (Fig. S6). This site refines as a sodium with a B factor of 28.1 Å^2^ compared to an average of 24.8 Å^2^ for protein atoms with which it interacts. The position after the first aspartate of the DXXXXD motif is most commonly leucine or methionine, with asparagine (as in HMWP2-Cy2) being the fourth most common (Fig. S7). There does not appear to be any consistent preference for a polar amino acid at the second position of the DXXXXD motifs within a subset of Cy domain sequences from siderophore biosynthesis. Even an enzyme catalyzing cyclodehydration of an identical peptide as HMWP2-Cy2, the second Cy domain of pyochelin synthetase (PchF-Cy2), instead contains a phenylalanine at this position (Fig. S8).

### Carrier protein interaction surfaces

The likely HMWP2-PCP1 binding site on Cy2 lies at the N-terminus of α8, where the donor Ppant would extend between the C-termini of β8 and β10 in the C-terminal subdomain, near the N-termini of helices α4 and α8 (Fig. S9A)^55,58^. In that region, the structure of Cy2 differs from crystal structures of existing Cy domains principally at the loop extending from β10, which adopts a conformation in between those seen for Epo^51^ and BmdB^52^ Cy domains (Fig. S9B, Fig. S9C contains a comparison of this region in HMWP2-Cy2 and more open C domain models). Despite this difference, all three upstream tunnel entrances remain unavailable for donor substrate binding. This loop adopts a similar conformation to BmdB-Cy2 in PchE-Cy1 cryoEM structures^55^ despite the PchE-Cy1 entrance being open in some cases, revealing that entrance accessibility in that structure is determined predominantly by the conformation of a conserved phenylalanine at the opening (PchE F372, HMWP2 F1755) rather than the conformation of the loop^55^. The NMR solution structure of HMWP2-PCP1^60^ was used to investigate potential PCP1-Cy2 interaction surfaces with the conformation displayed by Cy2. To better mimic a tethered Ppant, a HMWP2-PCP1 model bearing a phosphoserine at S1439 (SEP1439) was prepared and docked against HMWP2-Cy2 using HADDOCK2.4^61–63^, which has parameterization for phosphoserines and can take as input information about residues believed or known to be involved in interactions. Guiding the phosphoserine toward the expected binding site, near what would be the upstream tunnel entrance in an open Cy conformation, returned clusters of poses that resemble the C-bound, upstream CP structure from linear gramicidin synthetase, LgrA-PCP1-C2^58^ (Fig. 3A and S9A) and the recently reported cryoEM PchE structure^55^ (Fig. S9A). For the binding site to become accessible to a donor Ppant in HMWP2-Cy2, β10 would likely need to splay away from β8 by 2-3 Å relative to its position in the crystal structure (Fig. S9D). An analysis of surface electrostatic potential at this interface clearly shows the footprint of charge complementarity that favors this binding mode (Fig. 3B). In this position, it is apparent that the omitted 6-residue linker can be constructed to tether HMWP2-PCP1 to HMWP2-Cy2 without requiring rearrangement of either model. The model complex reported here includes the interacting residue pairs (in HMWP2 numbering) E1836_Cy_-R1444_PCP’_ K1717_Cy_-D1438_PCP_ (4.4 Å apart in the presented model but forming a close pair in a lower-ranked pose from the same cluster), and N1801_Cy_-S1459_PCP_ and involves hydrophobic packing around L1800_Cy_ and F1462_PCP_ (Fig. 3C,D).

**Figure 3.**
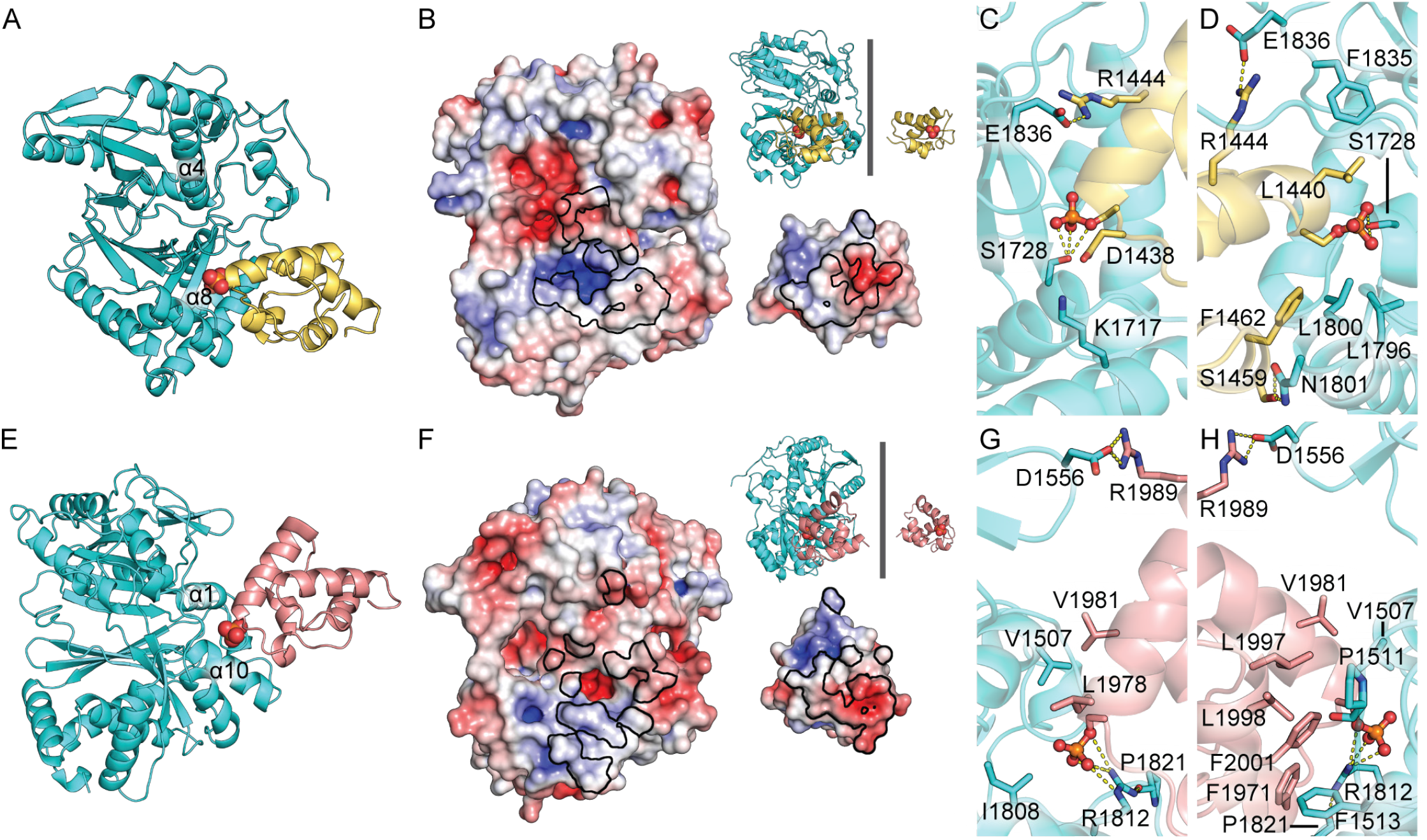
Protein-protein docking poses of HMWP2-PCP1 and HMWP2-PCP2 with HMWP2-Cy2. **A**, An NMR-based model of HMWP2-PCP1 (yellow cartoon with the phosphate of phosphoserine in spheres with elemental coloring) is shown bound at the expected upstream PCP-binding site of HMWP2-Cy2 (cyan). **B**, Electrostatic potential maps of the upstream PCP-binding site of HMWP2-Cy2 (left) and the Cy-binding site of HMWP2-PCP1 (right) are related here by a rotation of 180° so that the interaction surfaces of both components are displayed. The orientations of the components can be observed in the inset cartoon representations. The interaction footprint shared by the proteins is outlined in black. **C**,**D**, Detail of interacting residues (sticks) at the PCP1-Cy2 interface (phosphoserine is displayed in ball and sticks here). **E-H**, Same as A-D but for the downstream HMWP2-PCP2 interaction (PCP2 is in salmon).

The PCP-binding site at the downstream tunnel entrance, which interacts with the acceptor PCP (HMWP2-PCP2) for condensation and cyclodehydration, provides a point of contrast relative to prior Cy domain crystal structures. Specifically, loop 1 (after α1) extends further toward α10 than in EpoB-Cy^51^ or BmdB-Cy2^52^, in which it is blocked by the loop following α10 (Fig. S10A-C). Docking experiments similar to those conducted for HMWP2-PCP1 were conducted with a homology model of HMWP2-PCP2 built on the PCP from *Streptomyces* sp. Acta 2897 (~34% sequence ID, PDB ID 4PWV)^20^. Restrained docking in HADDOCK2.4 using a model of HMWP2-PCP2 prepared with a phosphoserine at S1977 (SEP1977) returned clusters (Fig. 3E) that resemble the Cy-PCP complexes of PchE^55^ and the C-PCP complex of AB3403 (PDB ID 4ZXI)^59^ or, to increasingly lesser degrees, FscG (PDB ID 7KW0)^64^ or ObiF1 (PDB ID 6N8E)^65^ and had favorable metrics (for the cluster containing the top result presented here: HADDOCK score −183.2 +/− 6.5, cluster size 34, 1404.9 +/− 88.2 Å^2^ buried surface area, and negative van der Waals, electrostatic and desolvation terms). As with the upstream site, analysis of surface electrostatic potential at the downstream interface clearly shows the footprint of charge complementarity that favors this binding mode (Fig. 3F). This interface involves hydrophobic packing of V1507_Cy_ against L1978_PCP_ and V1981_PCP_ and of P1511Cy against L1997_PCP_, as well as stacking of the phenyl ring of the weakly conserved F1513_Cy_ with both F1971_PCP_ and F2001_PCP_ (Fig. 3G,H). The other main contribution to HMWP2-PCP2 binding according to our model is from electrostatic interaction between a conserved arginine in α10 (R1812) and SEP1977. As input for the HMWP2-PCP2 docking runs, combinations of rotameric states for F1513 and R1812 other than those used for the models reported here were also tested but yielded lower scoring models. In the lesser poses from the full collection of results, there are some PCP_acceptor_ orientations that more closely resemble the ObiF1-C-PCP complex (PDB ID 6N8E)^65^, but these poses do not have favorable metrics. Notably, the downstream entrance of PchE-Cy1 harbors several differences relative to the Cy domain crystal structures including a different conformation of loop 1 lacking a phenylalanine (having a glutamate instead) and having an additional arginine at the C-terminus of helix α10.

Near the downstream interaction surface, two interesting regions of difference between Cy domain crystal structures exist. First, a conserved arginine in α1 (R1509), which forms a salt bridge with the first aspartate of the DXXXXD motif in EpoB-Cy^51^ and PchE-Cy1^55^, is oriented completely away from the center of the Cy domain in HMWP2-Cy2 (Fig. S10). In BmdB-Cy2^52^, the corresponding arginine adopts an intermediate position between the EpoB-Cy and HMWP2-Cy2 positions. Similarly, a conserved glutamine in the loop after the latch (Q1858), which forms hydrogen bonds to loop 1 in HMWP2-Cy2, is found projecting its side chain into the tunnel in EpoB-Cy (and PchE-Cy1), and a corresponding intermediate position is observed in BmdB-Cy2 (Figs. S10 and S11). In Figs. S11-15, the environment around Q1858 and the Ppant-binding region is compared with EpoB-Cy^51^, BmdB-Cy2^52^, PchE-Cy1^55^, AB3403-C^59^, FscG-C3^64^ and ObiF1-C^65^ (including Ppant molecules from the C(y)-PCP complexes), and Fig. S16 shows how reference C-PCP_acceptor_ models compare in alignment of their C domains. It appears that positioning of R1509 and Q1858 may provide a link between binding of PCP2, the state of the downstream interface—especially loop 1 containing F1513—and the placement of residues within the active site for catalysis, such as Y1505 and residues preceding Q1858. Interestingly, this region appears flexible in Cy1 as determined by relaxation dispersion experiments, in line with conformational malleability required for our model.

### Exploration of cyclodehydration intermediate binding in HMWP2-Cy2 highlights potential roles for active site tunnel residues

We sought to determine whether the active site tunnel of the HMWP2-Cy2 crystal structure could accommodate its cognate cyclodehydration intermediate. Because nucleophilic attack during cyclization can occur on either face of the condensation product amide, we employed rigid receptor covalent docking to sample potential binding modes of both 2*R* and 2*S* diastereomers of the intermediate mimic (2*R/S*,4*R*)-2,2-hydroxy-(2′-(2″-hydroxyphenyl)-thiazolinyl)-thiazolidinyl phosphopantetheine (Ppant-2HPTT-OH) (Fig. S17). Covalent, induced fit docking was performed using Glide^66–68^ and Prime^69–71^ in BioLuminate^72^ with the putative catalytic dyad in both acidic (neutral, presumed after cyclization) and basic (anionic, presumed before cyclization) states. The covalent docking approach permits identification of docked poses that are compatible with covalent linkage to the conserved serine of our HMWP2-Cy2-PCP2 complex (SEP1977). Both tested diastereomers of the hydroxythiazolidine moiety docked with favorable Glide scores in the HMWP2-Cy2 tunnel, returning in total 120 poses which were then refined to search for induced fit changes favoring various substrate orientations. Models of top-ranked poses obtained by this approach as well as their docking scores, energies from refinement, MM-GBSA binding free energies (detailed in the Methods section) and dyad ionization states are displayed in Figs. S18-23. These poses can be organized into three categories based on the direction in which the leaving group oxygen of the hydroxythiazolidine moiety is directed: 1) hydroxyl roughly toward the N-terminal end of helix α4, 2) hydroxyl toward the N-terminal subdomain β sheet, or 3) hydroxyl roughly toward the putative catalytic dyad. Depending on the chirality, the direction of the relevant C-O bond vector corresponds to different potential interaction configurations between Cy domain active site features and the other reactive groups of the intermediate, namely the thiazolidine ring N-H and S. Overwhelmingly, poses in the receptor with the neutral dyad were more favorable for both diastereomers. Fig. 4 shows top poses for each hydroxythiazolidine diastereomer, which both direct the leaving group oxygen (previously of the carbonyl electrophile) toward the N-terminus of helix α1.

**Figure 4.**
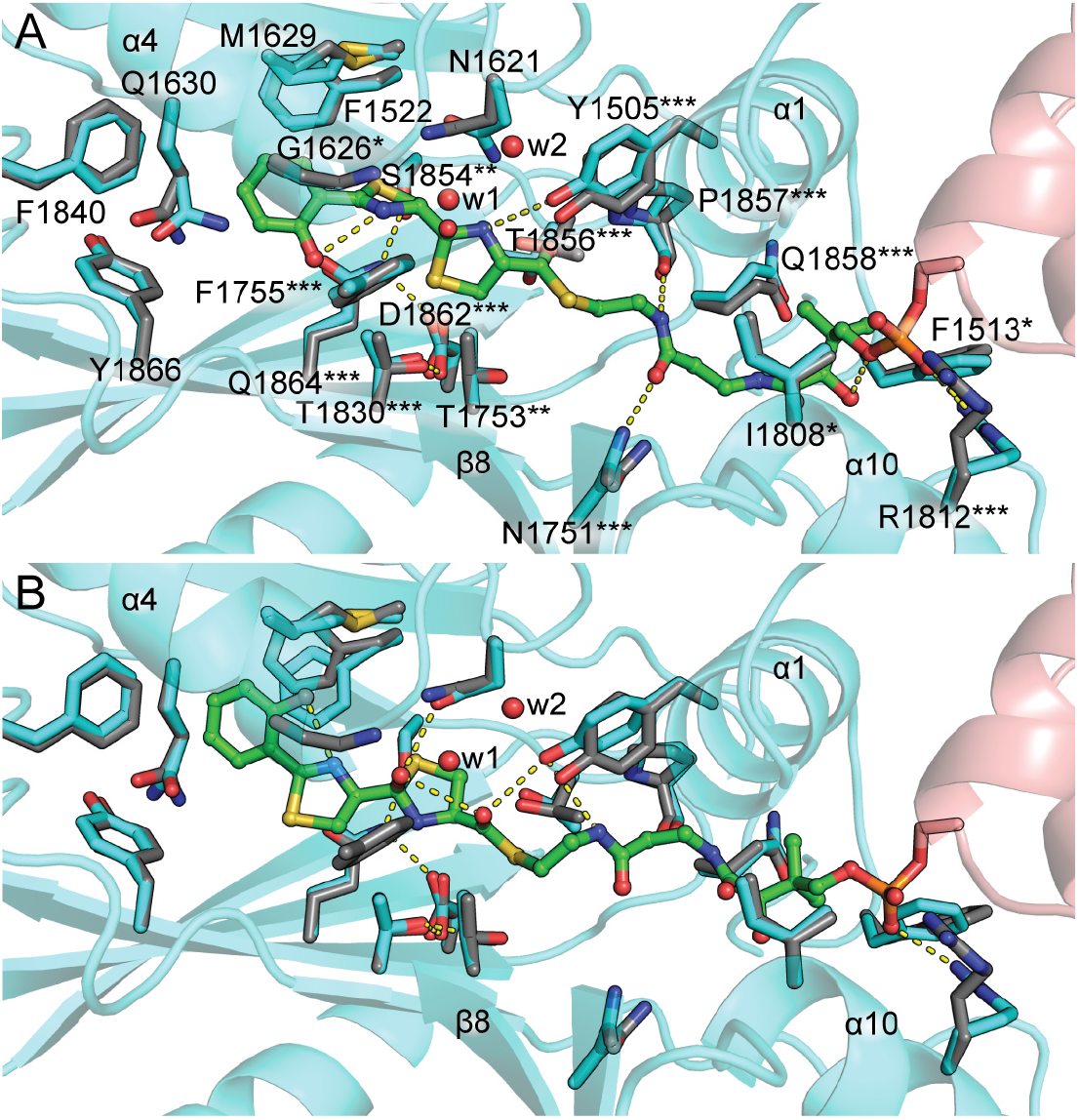
The HMWP2-Cy2 crystal structure acceptor tunnel and active site accommodate a cyclodehydration intermediate. **A**, Covalent docking of Ppant-2HPTT-OH in an HMWP2-Cy2-PCP2 complex based on the Cy2 crystal structure and PCP placement from protein-protein docking supports a model in which catalysis proceeds through an *R* hydroxythiazolidine intermediate (green) in HMWP2-Cy2 (refined covalent docking Cy model in cyan and PCP in salmon with transparent cartoon, and crystal structure coordinates for displayed residues in gray). This pose (#10 in Fig. S21) is selected as a top candidate based on its Glide docking score from covalent docking, MM-GBSA-approximated binding energy for the corresponding noncovalent phosphonate-S1977A complex and ligand and receptor strain energies. Two water sites (red spheres) are shown in the vicinity of the active site (w1 and w2). HMWP2 residues marked with asterisks agree with conservation trends at positions > ca. 2 bits in the sequence logo (Fig. S7) that are somewhat conserved (*, property conserved or HMWP2-Cy2 residue in at least ~25% of aligned sequences), moderately conserved (**, primarily two options and HMWP2-Cy2 residue at least ~50%), or highly conserved (***, overwhelmingly one option). **B**, The top *S* hydroxythiazolidine intermediate (#1 in Fig. S18).

In *S* intermediate poses, the leaving group oxygen is preferentially directed into a cavity along the side of α4 (52/60 poses, Fig. S18). In most of this category of *S* intermediate poses (49/60 poses), the ring sulfur and nitrogen are not positioned well to interact with the putative catalytic dyad. Notably, the Ppant arm appears to prefer a conformation in which the dimethyl moiety points in the direction opposite its orientation in reference models from AB3403 and FscG (PDB IDs 4ZXI and 7KW0, respectively)^59,64^, somewhat resembling the position observed in the ObiF1 model PDB ID 6N8E^65^. In this position, the Ppant arm can hydrogen bond and/or form polar contacts with the side chain of the conserved residue N1751 as well as the backbone carbonyls of the conserved residues P1857 and Q1858 and possibly the backbone N-H of M1825. Furthermore, some of these poses suggest that Y1505 can form an interaction with the Ppant amide nearest the thioester linkage. The other subset of the *S* intermediate hydroxyl-toward-α4 category contains only 3 representatives but is intriguing because it places the leaving group oxygen nearer the supposed donor carbonyl position (from which it is derived) at the N-terminus of α4 expected based on the trapped condensation donation state of LgrA-C2 (PDB ID 6MFX)^58^. The second highest-rated pose of this subcategory demonstrates possibilities for Ppant-protein interactions like those in the main subset of hydroxyl-toward-α4 poses just described, involving N1751 and the backbone N-H of M1825, but it also demonstrates hydrophobic packing of the dimethyl moiety against surfaces of α1 and α10 like the AB3403 and FscG references^59,64^. The pose also permits hydrogen bonding between the hydroxythiazolidine ring nitrogen and the dyad aspartate.

The *R* intermediate poses overall score similarly to the *S* poses, but a few notable differences are observed between these sets (Figs. S21-23). First, in one category of poses the leaving group oxygen is directed roughly toward the partial positive charge of the N-terminus of α4. This position is more compatible with expectations based on the trapped condensation donation state of LgrA-C2 (PDB ID 6MFX)^58^ and more closely resembles the cyclodehydration intermediate binding model proposed by Bloudoff et al.^52^, although the orientation of the hydroxythiazolidine in our model places the leaving group oxygen directed away from T1830, rather than toward it. This set of poses places the ring sulfur in the middle of the putative catalytic dyad and the hydroxyls of Y1505 and T1856 in positions in which it appears they could act as hydrogen bonding partners with the ring nitrogen. As is observed in the *S* poses, the *R* poses permit interactions between the Ppant arm and N1751, P1857 and Q1858, and the Ppant dimethyl is preferentially rotated away from the position expected based on AB3403 and FscG C-PCP_acceptor_. didomains mentioned previously^59,64^. Variation within this category alternatively suggests that T1856 could be involved in an interaction with the Ppant-2HPTT-OH thioester. Additional observations within the full set of docking results, including descriptions of possible donor side chain interactions, are presented in the Supplemental Results section of the Electronic Supporting Information document.

The top cyclodehydration intermediate poses acquired by these means were overlaid on the Cy2 structure, including two active site water molecules consistently observed in Cy domain crystal structures (Fig. 4). These water molecules between helix α4, the DXXXXD motif loop and Y1505 were found to be roughly compatible with Ppant-2HPTT-OH positioning, although the water molecule nearest helix α4 might instead resemble positioning of the water leaving group. Taken together, these observations of potential cyclodehydration intermediate binding modes promote further consideration of the *R* binding mode. Additional support for this stance comes from more negative free energy of binding of the top *R* hydroxyl-toward-α4 pose as approximated by Prime^69–71^ MM-GBSA. Fig. S24 provides additional comparisons between downstream tunnels of C-PCP models and Cy domain models and the top *R* hydroxyl-toward-α4 intermediate pose. The Cy domains all share features of the downstream tunnel that could contribute to binding Ppant in the reported manner, and these interactions are distinct from those in C-PCP models. It should also be noted that the acceptor Ppant does not fill the entire volume of the downstream tunnel, implying that ordered water molecules are likely involved in Ppant binding.

By comparing C domain donor and acceptor substrate mimic models, a picture of the condensation preceding cyclodehydration can be obtained (Fig. S25). Here, we consider a mechanism in which a peptide bond is first formed as in C-domains—an intermediate observed in biochemical studies of Cy domains. We do not further consider the alternative mechanism involving a possible thioester intermediate. The PCP-C model from LgrA^58^ provides the donor electrophile positioning at the N-terminus of α4 as mentioned previously, and the C-PCP model from FscG (PDB ID 7KW0)^64^ or the covalently tethered acceptor mimic of CDA-C1 (PDB ID 5DUA)^73^ provides nucleophile positioning. Whereas the CDA-C1 mimic was demonstrated to be catalytically active, the FscG mimic may not be. Excitingly, the hydroxythiazolidine of the top *R* intermediate pose fits between the CDA-C1 nucleophile and the LgrA-C2 electrophile. Furthermore, comparison of the reactive region of the cyclodehydration intermediate in the pose reported here and the product reported for PchE-Cy1^55^ shows good agreement with respect to placement of the newly formed (hydroxy)thiazol(id)ine (Fig. S26).

### Low-frequency vibrational normal mode approximations for HMWP2-Cy2

Since Cy domain crystal structures have been solved in closed conformations that preclude donor substrate binding, we sought to explore flexibility of the HMWP2-Cy2 model around the putative cyclodehydration-like state using the normal mode approximation. Although our top cyclodehydration intermediate candidate pose is promising, it is also interesting to consider how flexibility of the Cy domain alters relationships between groups within the active site.

The 10 lowest-frequency normal modes of the crystallized HMWP2-Cy2 conformation were calculated using a model to which additional water molecules were added to fill voids (primarily in the active site tunnel). Overall, the observed modes are characterized by relative motion of the N- and C-terminal subdomains (see movies in Electronic Supporting Information). The lowest-frequency mode (number 1) involves counter rotation of the N- and C-terminal subdomains about an axis running approximately through the centers of mass of the two subdomains. The second mode also involves rotation, but in this mode it is about an axis running between the subdomains, roughly from upstream to downstream entrances, and is characterized by modulation of the distance between α3 and α11 (please refer to Fig. S3 for complete secondary structure assignments). These motions are reminiscent of variation among crystal structures. Mode 3 resembles both modes 1 and 2 in some respects, but it also includes displacement of α10 away from α1 at the downstream tunnel entrance, which suggests the potential for coupling between the state of the downstream entrance and the global conformation of the Cy domain. Mode 4 is very similar to mode 2. Notably, these lowest-frequency modes alter the shape of the PCP_donor_ binding site and appear to affect the accessibility of the N-terminus of α8 where the Ppant phosphate is expected to bind. Mode 5 is a highly localized motion of R1817 at the C-terminus of α10 at the downstream opening. R1817 forms a salt bridge with D1679 in α6, following the hinge region that is disordered in the HMWP2-Cy2 crystal structures. Although mode 5 does not directly involve this hinge, the hinge undergoes motion in all of the other modes (this motion is, however, small in mode 6). The motions depicted by these normal modes are consistent with recent NMR studies depicting a large global motion of the N-terminal subdomain with respect to the C-terminal subdomain.

Accompanying intersubdomain motion, there is more intrasubdomain displacement in the higher-frequency modes 7-10, especially between two lobes of the C-terminal subdomain: 1) the floor loop, the strands preceding and following it (β8/9) and helices α6, 9 and 10; and 2) strands β7,10,12 and 13 and helices α7, 8 and 11. One mode (number 7) had considerably greater relative amplitude than the other modes (Fig. S27), and it is particularly interesting because it results in opening of the upstream tunnel entrance by over 3 Å, albeit involving some unphysical deformations of the model geometry owing to the approximate method. We note that it resembles an amplified combination of the lower-frequency modes in that rotations of N- and C-terminal subdomains result in modulation of the α3-α11 distance and displacement of α10 relative to α1 is observed. It also bears similarity to higher-frequency modes affecting the internal state of the C-terminal subdomain by opening the upstream entrance.

Carried by these global motions, relative motion of residues of interest in the first and second shells of the active site was considered further. These residues include Y1505, the putative catalytic dyad (T1830-D1862), conserved floor loop/α1 linkage involving T1499, D1770 and T1772, the upstream opening between L1754 and S1831, and the D1620 (<u>D</u>XXXXD)/R1509 (α1) and D1625 (DXXXX<u>D</u>)/R1757 salt bridges (Fig. S28). Several modes demonstrate modulation of the active site dimensions in ways that would influence substrate binding. Modes 1, 6, 7 and 8 modulate width along a vector between the floor loop and the latch strand β11 (T1772-T1856). Modes 7 and 9 modulate the distance between the conserved aspartate of the floor loop (D1770) and the active site aspartate (D1862). Modes 6-10 modulate the distance between Y1505 and the active site aspartate, and motions associated with the open-closed transition bringing the catalytic dyad toward and away from Y1505, when considered alongside our top *R* cyclodehydration intermediate pose, suggest global motion may couple to substrate rearrangements after condensation to obtain a cyclization reaction complex. Additionally, the environment of D1770 has been identified as a region of fluctuating electrostatic potential by our coauthor, and alternating between states in which D1770 is more surface exposed (contributing to the transiently observed surface negative electrostatic potential) or closer to the active site may play a role in managing charge and hydrogen-bonding networks during turnover.

The proposed role of Y1505 that could stem from its interaction with the acceptor Ppant thioester is stabilization of the acceptor in a geometry conducive for condensation. Compressing the distance between the dyad—envisioned to be interacting with the acceptor cysteine thiol(ate)—and Y1505 may facilitate a shift from Y1505 interacting with the Ppant thioester to its interacting with the newly formed amide’s nitrogen. In this way, tyrosine would be preferred at this site for its geometric compatibility with stabilizing the condensation product and cyclodehydration intermediate states as well as facilitating the transition between them. This consideration and its position in α1, combined with its ability to act in proton relays and adopt a phenolate form make it an interesting candidate residue to underly a mechanism by which the Cy domain could sense the state of the active site and respond with global conformational changes.

### Potential inhibitory hot spots for HMWP2-Cy2

To identify possible locations within the HMWP2-Cy2 structure for future inhibitor development, the structure was analyzed using the FTMap server (Fig. S29)^74^. Several ligand hot spots were identified within the protein tunnel (Fig. S29A-C), consistent with the idea of mimicking Ppant-tethered intermediates as possible inhibitors. Of these hot spots, the cluster in the side chain-binding pocket is of interest for its enrichment in ring structures and inclusion of phenol, which roughly mimics the 2HPT moiety’s orientation in the top *R* intermediate pose. The active site clusters are compared to the top *R* intermediate pose in Fig. S30. Phenol is deeper into the side chain binding region than the 2HPT moiety of the docked pose, suggesting 2HPT may not be fully accessing the basin of attraction occupied by phenol in this volume. Also, the clusters near the N-terminus of helix α4 or next to the active site dyad place polar groups in orientations consistent with orientations of polar groups in the intermediate docking results. On the other hand, the Ppant-binding region does not return clusters exhibiting specific polar contacts, and instead it appears that binding of the small molecule probes in this region is largely driven by burying hydrophobic surface area. Excitingly, three additional hot spots were identified outside of the protein tunnel: a minor site was located at the expected donor binding site (Fig. S29D), and two more populated sites were found nestled between the floor loop and α1 (Fig. S29E,F). The donor site cluster and floor loop/α1 clusters also seem to be driven largely by hydrophobic packing, but the donor site also demonstrates polar contacts with S1728 at the predicted Ppant phosphate binding site. Therefore, in addition to competitive inhibitor development based on intermediate mimics, sites at the PCP binding sites or around the floor loop may enable development of an alternative class of inhibitors that target C domain family members. In particular, the floor loop region has been shown to undergo fluctuations in surface electrostatic potential across the conformational variation of the C domain family^53^ despite divergent sequence-level features. Conserved motions’ effects on the electrostatic potential and divergent sequences may allow blocking a key conformational transition in a particular Cy domain with some degree of specificity.

## Discussion

The crystallographic structures of HMWP2-Cy2 presented here add to the limited array of high-resolution structural information about Cy domains acquired to date and include characterization of a siderophore system linked to pathogenicity of *Y. pestis.* The observed more-open structure of HMWP2-Cy2 and its different organization around the downstream CP binding site were opportune for exploring PCP_acceptor_ docking to a Cy domain. Here we demonstrate that molecular docking with phosphoserine-modified PCP models identifies binding modes stabilized by complementary Cy-PCP interfaces that resemble observed binding orientations within prior Cy-PCP and C-PCP complex models^55,58,59,64^. A potential similarity between the downstream entrances of C and Cy domains is the conserved arginine in Cy domains (R1812 in HMWP2-Cy2) that can adopt a conformation satisfying the same electrostatic contribution as conserved arginine residues at the downstream entrances of C domains. Based on comparison with the recent PchE cryoEM structures^55^, it may also be that Cy-PCP_acceptor_ complexes use this arginine to bind the Ppant phosphate in a manner similar to what is observed in ObiF1-C^65^; however, PchE differs markedly from the other Cy structures at the downstream opening in its positioning of loop 1 (bearing a glutamate where Cy crystal structures have a phenylalanine) and in the number of arginine residues in α10. These differences allow for interactions with Ppant and the acceptor PCP that would not be available in EpoB-Cy^51^, BmdB-Cy2^52^ and HMWP2-Cy2. We note that the three conformations of the loop following helix α1 observed in Cy crystal structures may be selected by crystal packing^51,52^. Intriguingly, the crystal contacts formed by this region of the HMWP2-Cy2 structure are with helix α7 in a symmetry-related molecule, and the orientation of that helix is reminiscent of the predicted orientation of PCP2 helix α3′ in the Cy-PCP_acceptor_ complex, which harbors the hydrophobic residues around F1513 that we predict to form a substantial part of the hydrophobic interfacial surface. It is possible that this similarity between crystal contacts and the putative PCP2 orientation accounts for the different loop conformation following helix α1 in the HMWP2-Cy2 structure.

Prior Cy crystal structures show a closed state in which the upstream active site tunnel entrance, predicted by analogy to condensation domains, is not accessible. This is also the case for the HMWP2-Cy2 structures reported here. Protein-protein docking of HMWP2-PCP1-Cy2 suggests that PCP1 would interact with the floor loop and helices α7 and α9, even in the case of a closed Cy conformation. What seems clear is that condensation-competent Cy domains prefer the closed state in the absence of PCP interactions, so introduction of a PCP may be required to shift the conformational equilibrium toward the open, condensation state, which was similarly observed in cryoEM studies of PchE^55^. Owing to the proximity of T1753 and T1830 (of the putative catalytic dyad), which flank the upstream opening, it is interesting to consider how breaking or forming interactions with these residues would influence the global conformational change between open and closed states. In some of our docking models, T1753 is nearly able to interact with the sulfur of the acceptor Ppant thioester linkage.

In comparison to C domains, there are additional catalytic demands for the cyclization reaction involving constriction of the condensation intermediate to overcome ring strain upon forming the thiazoline and efficient proton transfer to quench the cyclization transition state to yield the hydroxythiazolidine intermediate for dehydration. The closed Cy domain state has also been proposed to facilitate the catalytic requirement of cyclodehydration at least in part through exclusion of bulk water^52^. It has remained an assumption, however, that the Cy domain conformations crystallized to date resemble a catalytic state of the enzyme. Importantly, docking performed here suggests how substrate placement can utilize several features of the fold. First, the cysteine thiol nucleophile can be positioned at the putative catalytic dyad. In this position, internal motion of the C-terminal subdomain, perhaps additionally involving global intersubdomain motion, could compress the reaction complex along the S-C bond axis. Furthermore, binding in this manner could support deprotonation of the ring nitrogen by a mostly conserved residue (Y1505) and sequestration of the water leaving group in a position that already binds water in the crystal structures (near the N-terminus of α4). A phenylalanine mutant at the tyrosine equivalent to HMWP2 Y1505 in the EpoB Cy domain gave a moderate 6-fold decrease in reaction rate^51^, suggesting that another feature, such as an ordered water, may be able to compensate for its mutation. This could explain why this tyrosine is not strictly conserved. Structural alignments with extant structures of C-PCP_acceptor_ complexes harboring Ppant molecules demonstrate that Ppant conformations in those models are inconsistent with placement of a cyclodehydration intermediate in the Cy active site. This difference between C and Cy domains is due to the Cy domain’s more-closed nature, which reduces the distance between the downstream tunnel entrance and the N-terminus of helix α4. If this closed state resembles a catalytic state, then it is implied that the Ppant conformation in that state differs from the Ppant conformations observed in C-PCP_acceptor_ complexes to date.

Following the principle of parsimony, we show that an amine-first condensation in Cy domains (analogous to C domain activity) could, with minimal rearrangement, result in an intermediate with the acceptor cysteine thiol positioned near the conserved active site aspartate and the electrophilic carbonyl of the newly condensed peptide positioned at the N-terminus of helix α4, similar to the putative position of the donor thioester in the LgrA condensation donation complex (cf., PDB ID 6MFX)^58^ (Fig. S25). In either amine- or thiol-first condensation, the electrophilic carbonyl could be found at the N-terminus of α4. Considering this orientation of the donor electrophile for condensation, zwitterionic transition state stabilization would facilitate nucleophilic attack, as has been proposed in C domains^75^. With respect to this concept, we also note that the crystallographic sodium site resides on the wall of the active site opposite the upstream opening and N-terminus of α4, where stabilization of a positive charge could contribute to stabilizing a zwitterionic intermediate or even deprotonating it using an ordered water molecule.

Envisioning extending the C–S bond formed in the cyclization reaction from its length in our docked cyclodehydration intermediate model to a pre-cyclization state reveals the potential for change in this bond-forming coordinate to be accompanied by a large-scale conformational change between open and closed states approximated in our low-frequency normal mode analysis. This is in agreement with the recent solution NMR studies from our coauthor’s laboratory^53^ that demonstrate substantial flexibility of the first Cy domain from Ybt synthetase. Katsuyama et al.^54^ also report disorder in the DXXXXD motif of their Cy structure, therefore flexibility of the two subdomains is likely to be a feature of the entire family. The conformational change we describe brings the putative catalytic dyad closer to the center of the active site, potentially positioning it to act as a base while compressing the C–S axis for bond formation (Figs. 5 and S31). The NMR study further indicates that the conserved, putative catalytic aspartate plays a key role in global dynamics that mediate allosteric communication of CP binding^53^, which works with our model to suggest how PCP binding, global conformation, and the stage of catalysis may be related.

**Figure 5.**
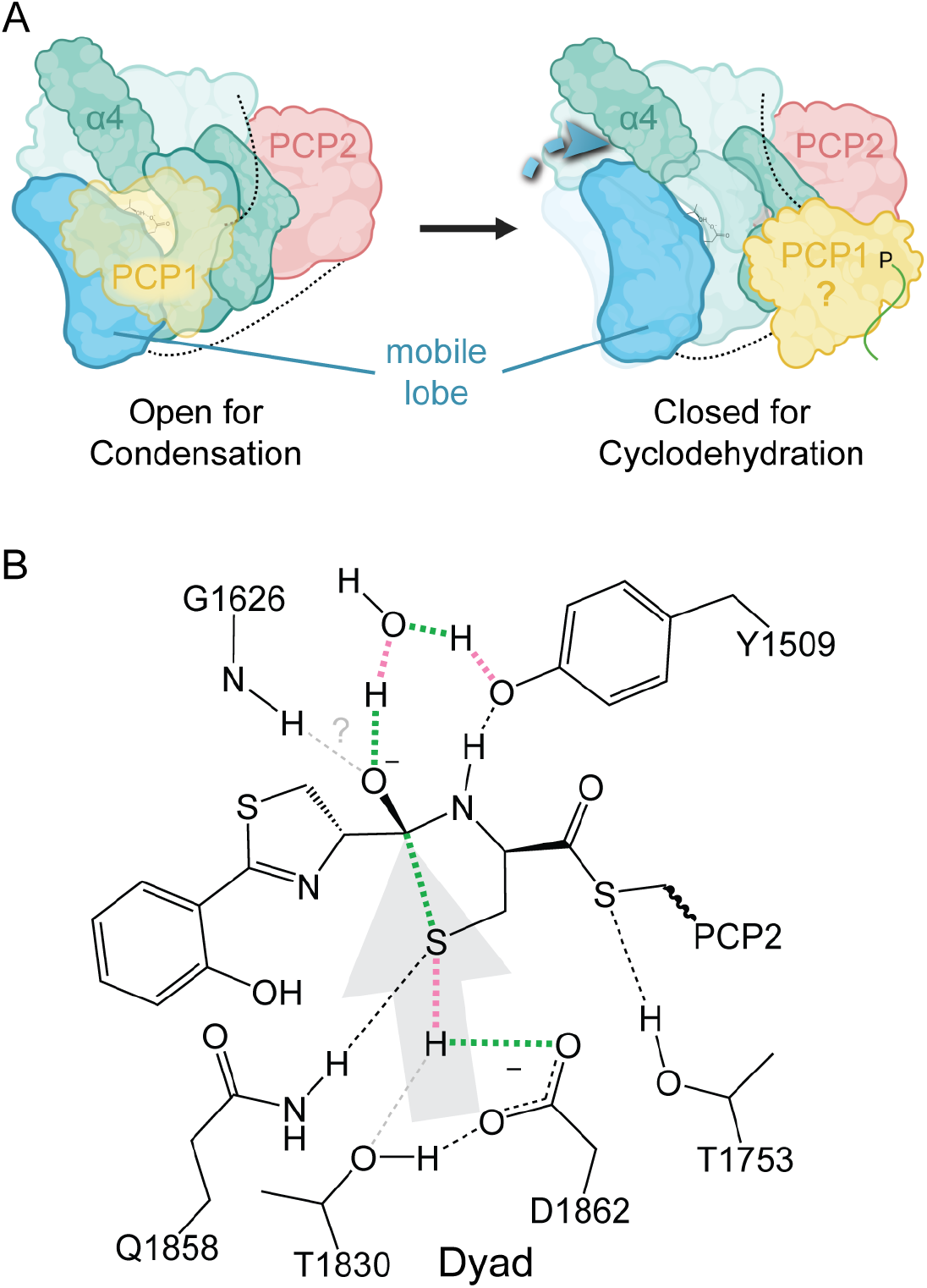
A proposal linking a global Cy domain conformational change to ring closure in the cyclodehydration reaction. **A**, After the condensation reaction, the PCP_donor_ (yellow) dissociates from its position in the open-for-condensation state (Cy domain in green) at least enough to remove the pantetheine moiety from the upstream tunnel entrance. Helix α4 is labeled for reference, and other darker green regions in the open-for-condensation state are the floor loop and hinge regions. To obtain the closed-for-condensation state, one lobe of the C-terminal subdomain (blue, mobile lobe), containing T1830 and D1862, rotates through 5-10° around its point of contact with the floor loop strands. It is unknown what position the PCP_donor_ occupies in the closed-for-cyclodehydration state. The PCP_acceptor_ position is probably similar in both states. **B**, During the transition from the open-for-condensation state to the closed-for-cyclodehydration state, the condensation product collapses into a cyclization reaction complex, with motion of the C-terminal β strands harboring the dyad (light gray arrow) contributing to compression of the nucleophile-electrophile distance. A representation of the envisioned transition state is depicted. Green and pink dashed lines indicate bonds formed or broken during this reaction step, respectively. Light gray dashed lines indicate where alternate/additional interactions could form and our approximate model remains ambiguous. Panel A was prepared using BioRender.

In addition to the putative catalytic dyad, Cy domains contain a conserved phenylalanine at the C-terminal end of β8 leading into the floor loop (HMWP2 F1755). In crystallographic Cy domain structures solved to date, this residue is consistently found to contribute to a blockade of the upstream tunnel entrance and is the basis for the suggestion that Cy domains exclude bulk solvent from the cyclodehydration active site^52^, thereby facilitating the dehydration step. For condensation-competent Cy domains, this phenylalanine residue rotates away from the end of helix α4^55^, and it might be that early recognition events between the Cy domain and the PCPs alter the conformational preferences of the phenylalanine and nearby residues. This phenylalanine is in the region of varying electrostatic potential identified recently by Mishra et al.^53^, where varying the electrostatic potential may contribute to flipping toward a condensation-competent state. Mutation of this residue was found to decrease cyclodehydration activity to 31.6% of wild type in PchE^55^ and to decrease cyclodehydration activity to ~20% of wild type in BmdB (with a concomitant increase in release of the linear peptide intermediate)^52^, clearly demonstrating its importance in Cy domains. Notably, although this residue is largely conserved in Cy domains, it is absent in some tandem Cy domain contexts where one of the two Cy domains is expected to catalyze condensation only (further discussed in the Supplemental Discussion, alongside additional thoughts on catalysis, in the Electronic Supporting Information document).

Collectively, the Cy domain structures to date, structural comparison with prior PCP_donor_ -C and C-PCP_donor_ complexes, and molecular docking work presented here lead us to propose that conserved residues of the downstream tunnels in Cy domains, thought to be involved in Ppant binding, may dynamically contribute to the energy landscape of an open-to-closed transition upon post-condensation PCP_donor_ release. We envision this to involve a 2-3 Å translation of atoms in C-terminal β sheet strands 7,10,12 and 13 relative to C-terminal β sheet strands 8 and 9. In the open state, the putative catalytic dyad would be shifted away from the opposing side of the active site formed in part by Y1505 in helix α1. This would open additional tunnel volume for binding the cysteine thiol near the dyad in the event of initial condensation by the amine nucleophile analogous to the activity of C domains.

Upon transitioning toward the closed-for-cyclodehydration state (Fig. 5), the distance between the cysteine thiol and the electrophilic carbonyl carbon of the condensation product could be decreased as F1755 closes over the upstream entrance and the C-terminal β sheet reanneals. Translation along that path could be facilitated by rearrangement of the Q1858 and N1751 carboxamides and by the ability of Y1505 to act as either hydrogen bond donor or acceptor to the Ppant-linking thioester or the newly formed hydroxythiazolidine’s nitrogen. Adding some credence to the possibility of two spatially separated states for the condensation and cyclodehydration intermediates is the observation of better docking scores and lower potential energies for Ppant conformations in which the dimethyl moiety is directed toward the hydrophobic patches in the loops after α1 and α10. These poses differ from the reference models for downstream Ppant positioning in C domains^58,59^, which place the dimethyl moiety so that the methyl groups open toward α10 and the pantetheine thiol is further into the active site. In short, differences in the downstream tunnels of Cy domains relative to C domains may define a special basin of attraction that facilitates transition from the open-for-condensation state to the closed-for-cyclodehydration state by allowing rearrangement of the Ppant in a way not observed in C-PCP_acceptor_ structures to date. Perhaps this feature could be targeted in inhibitor design, especially considering the identification of ligand binding hot spots in the downstream tunnel.

Currently, work is underway to build a more comprehensive picture of dynamics in and around crystallized Cy domain states to test ideas about flexibility as it relates to PCP and substrate binding and catalysis. What seems clear is that condensation-competent Cy domains prefer the closed state in the absence of PCP interactions, so introduction of a PCP or tethered ligand may be required to shift the conformational equilibrium toward the open, condensation state. It appears likely that the closed-for-cyclodehydration state is favored in Cy domains by the organization of loop 1 and the interactions between α1, the first aspartate of the DXXXXD motif and the floor loop, which all differ from C domains. Additionally, the docking results here suggest how systems for trapping PCP-bound states or covalently tethered substrate mimics for crystallography may be devised using the ideas of inhibitory crypto ACPs^76^ or alkyl halide/cysteine-based covalent probes^73^, respectively. Further hypotheses for how mutations may affect dynamics and catalysis are also being tested. The results presented here contribute toward deciphering the complex behavior of Cy domains and present possible sites for inhibitor development targeting Cy domain activity in a crucial enzyme of a problem pathogen.

## Methods

### Cloning and Expression of HMWP2-Cy2

The encoding sequence for HMWP2-Cy2 was amplified from an expression vector containing the entire HMWP2 gene, kindly provided by the C.T. Walsh laboratory^77^. The sequence encoding P1480-Q1910 was amplified using Pfu Ultra II (Agilent) with primers purchased from IDT (forward primer: 5′-GGA AAA CC<u>C A/TA TG</u>C CGG TCG AAC AAC-3’; reverse primer: 5′-CG<u>G /GAT CC</u>T ATT GCC AGG CGC TTT CAT CC-3′; NdeI and BamHI binding sites are underlined, and / marks the cut site). Reactions were purified by electrophoresis and extracted using a Qiagen Gel Extraction kit. Subsequently, both pET28a (Novagen) and the recovered HMWP2-Cy2 DNA insert were double digested with NdeI and BamHI (NEB). The digested pET28a was further reacted with shrimp alkaline phosphatase (NEB) and allowed to react overnight at 16 °C. Digestion reactions were purified using a Qiagen Spin Clear and Concentration kit, and resultant DNA was used for ligation reactions with T4 DNA Ligase (NEB). Ligated product was transformed directly into chemically competent NEB5α cells (NEB) and plated on Miller lysogeny broth (LB-Miller)/agar solid medium supplemented with 50 μg/mL kanamycin. Transformant DNA was isolated and sequenced (Genewiz) to confirm the presence of the intact HMWP2-Cy2 DNA sequence.

The sequenced HMWP2-Cy2 construct was transformed into BL21(DE3) cells (Stratagene) for protein expression and purification. Eight 1 L LB-Miller cultures supplemented with 50 μg/mL kanamycin were inoculated at 16 °C with 5 mL of an overnight culture and allowed to grow for approximately 72 h. Cells were harvested at 5,000 rpm for 15 min in a JLA 9.1000 Beckman rotor at 4 °C. Cell pellets were immediately resuspended in approximately 40 mL of 50 mM Tris (pH 8.0), 100 mM NaCl, 5 mM β-mercaptoethanol, and 5 mM imidazole. The cell suspension was then sonicated using a Branson Sonifier at 50% amplitude, 1 s pulse/2 s off for a total of 2 min for 6 cycles. The cell lysate was then centrifuged for 1 h at 15,000 rpm in a JA 25.50 rotor at 4 °C. The resulting supernatant, including the soluble HMWP2-Cy2 protein, was further purified at 4 °C.

The HMWP2-Cy2 supernatant was loaded at 1 ml/ min onto a 5 mL HisTrap column (GE Healthcare) pre-equilibrated with lysis buffer. Protein was eluted using a 100 mL linear gradient from 0 – 300 mM imidazole. The purest protein-containing fractions as analyzed by sodium dodecyl sulfate – polyacrylamide gel electrophoresis (SDS-PAGE) were collected and dialyzed into 50 mM Tris (pH 8.0) with 5 mM dithiothreitol (DTT). Dialyzed sample was then loaded onto a 10 mL Mono Q column (GE healthcare) pre-equilibrated in the same buffer, and protein was eluted over a linear gradient from 0 – 500 mM NaCl. Protein fractions were pooled, concentrated, and run on a 16/600 S200 column (GE healthcare) pre-equilibrated in 50 mM Tris (pH 8.0), 100 mM NaCl, and 5 mM DTT. Protein-containing fractions were collected and concentrated to 10 mg/mL as determined by the Bradford method with standard bovine serum albumin. Protein was aliquoted, flash frozen in liquid nitrogen and stored at −80 °C. Analytical gel filtration was performed with an ENrich SEC 650 10 × 300 column (Bio-Rad) equilibrated and eluted with 50 mM Tris pH 8.0, 100 mM NaCl and 1 mM DTT at a flow rate of 1 mL/min. Protein standards were from Sigma-Aldrich (ovalbumin, bovine serum albumin and lysozyme) or MP Biomedicals (bovine γ-globulins) and run in 50 mM Tris pH 6.8 and 150 mM NaCl at 1 mL/min.

### Structure Determination

#### Crystallization of HMWP2-Cy2

Conditions for crystallizing HMWP2-Cy2 were identified from sparse matrix screening using a Phoenix micropipetting robot (Art Robbins). Prepared trays were stored and monitored at 4 °C within a Rock Imager 1000 (Formulatrix). Drops of 450 nL of 10 mg/mL protein in SEC buffer and 150 nL of MCSG-1 H10 (Anatrace; 0.1 M 4-(2-hydroxyethyl)-1-piperazineethanesulfonic acid (HEPES) (pH 7.5) and 25% (w/v) PEG 3350) yielded singular crystals of diffraction quality. Crystals were cryoprotected with crystallization solution containing 15% (v/v) glycerol, mounted on a nylon loop, and flash cooled in liquid nitrogen.

#### Data collection, phasing, and refinement

Diffraction data for the 1.94 Å-resolution HMWP2-Cy2 structure were collected at the MIT crystallization facility on a rotating copper anode X-ray generator (Micromax 007) using 1.0° oscillations. Data for the 2.35 Å-resolution HMWP2-Cy2 structure were collected at a wavelength of 0.9791 Å at 100 K at the Advanced Photon Source, Argonne IL, on beamline 24-ID-E, using 0.5° oscillations. Data were indexed to space group P4_1_2_1_2 using HKL2000^78^, and data statistics are presented in Table S1.

Phases for the 1.94 Å-resolution dataset were solved by molecular replacement in PHASER^79–81^ using the structure of EpoB-Cy (PDB ID 5T7Z)^51^, minus residues M1-S42 of the NRPS docking domain, as a search probe. The model was converted to the expected sequence of HMWP2-Cy2 with truncated side chains using CHAINSAW^82^, which yielded a PHASER solution with overall LLG and TFZ scores of 128 and 13.2, respectively. Initial electron density maps were interpretable and enabled direct building of the structure. Iterative rounds of model building were carried out in COOT^83,84^, followed by refinement in PHENIX^80,81,85^. Initial rounds of refinement included simulated annealing, energy minimization, and B-factor refinements. As the model refinement converged, only energy minimization and B-factor refinements were performed. After building the protein chain, a continuous region of electron density was observed near R1703 of the V-shaped opening between β7 and β11, just beyond the donor substrate side chain-binding region. Attempts to refine a chain of water molecules in this density returned positive difference electron density that suggested the presence of a polymer. Therefore, a fragment of a polyethylene glycol (PEG) polymer from the crystallization solution and cryoprotectant was modeled and refined with reasonable B factor values and electron density. In addition to this truncated PEG molecule, waters, and a sodium ion were placed and refined into electron density. Final rounds of refinement included occupancy refinement for alternate side chain confirmations, in addition to translation/libration/screw (TLS) refinement using optimal group definitions determined within PHENIX. The final model was verified using composite omit maps and Ramachandran angle analysis using MolProbity^86^, and no residues were found in disallowed regions. The final model contained all except 24 residues from the N-terminus, which included the hexahistidine tag, and residues 1663-1668 of the loop connecting the N- and C-terminal halves of the Cy domain structure. Refinement statistics are presented in Table S1.

The 2.35 Å-resolution HMWP2-Cy2 structure was solved by molecular replacement using the 1.94 Å-resolution structure as a search model in PHASER through PHENIX^79–81^. Rounds of model building and refinement were performed identically as above until the structure refinement converged. The final model was verified using composite omit maps and Ramachandran angle analysis using MolProbity^86^ and contained all except 24 residues from the N-terminus, which included the hexahistidine tag, and P1666.

### Model Preparation for Molecular Docking

#### Cyclization Domain

A complete HMWP2-Cy2 model was generated by combining residues 1483-1662 and 1669-1910 from the 1.94 Å-resolution structure with residues 1663-1668 of the 2.35 Å-resolution structure (keeping a nonredundant set of ordered water molecules from both structures). The model was prepared with neutral -CHO and -NH_2_ termini using standard methods in the BioLuminate Protein Preparation Wizard^72,87,88^: missing atoms were built in, charges were predicted using Epik^89,90^, expected bond orders were verified, and hydrogen bonding networks were optimized. Hydrogen coordinates were refined using the OPLS3e forcefield in the program Impact within the BioLuminate Protein Preparation Wizard^72,87,88,91–94^. All non-water ligands were removed from Cy domain models before preparation and simulations. Crystallographic water molecules were removed from the HMWP2-Cy2 models generated for molecular docking after preparation steps (see the *Cyclodehydration Intermediates* subsection for further information on HMWP2-Cy2-PCP2 models for covalent docking). The HMWP2-Cy2 model used for protein-protein docking was modified by altering the rotameric state of R1812 so that its guanidinium group was toward a location similar to the guanidinium group found interacting with the pantetheine phosphate group in most C-PCP complexes^58,59,64^. The modified R1812 rotamer was installed in a model with crystallographic waters, which was then subjected to hydrogen bond network optimization and hydrogen coordinate refinement before water molecules were removed and R1812 only was subjected to Prime^69–71^ energy minimization using the automatic method (up to 2 iterations of 65 minimization steps using conjugate gradient descent for large gradients and truncated Newton otherwise). HMWP2-Cy2 models for protein-protein docking ultimately included a set containing possible combinations of crystallographic rotameric states of F1513 and R1812 and the C-PCP-mimicking R1812 rotamer.

*Carrier Proteins*

The first model in the deposited NMR bundle for apo HMWP2-PCP1, PDB ID 5U3H^60^, was used for docking experiments. HMWP2 residues 1404-1476—comprising the folded core of the apo/holo conformation of PCP1—were used. This model excludes much of the flanking, flexible linker regions. Additionally, the conserved serine was phosphorylated to approximate electrostatic contributions from the phosphopantetheine phosphate of the holo PCP. Since no structure has been reported for HMWP2-PCP2, a homology model of HMWP2 residues 1943-2015 was generated in BioLuminate^72,95^ using as template a cytochrome p450-bound holo PCP crystal structure, PDB ID 4PWV chain B^20^, which shares 34% sequence identity with HMWP2-PCP2. HMWP2-PCP2 was also phosphorylated at its conserved serine residue. PCP models were prepared with neutral -CHO and -NH2 termini in BioLuminate^72,88–90,92^ similarly to the preparation of the Cy domain models. An orthorhombic TIP3P water environment with 10 Å buffer regions was built around the HMWP2-PCP2 model using the Desmond System Builder tool in BioLuminate^72,96,97^. The resulting system also contained 150 mM NaCl plus nine neutralizing Na^+^ ions. This system was relaxed using the default Desmond molecular dynamics relaxation protocol for the constant pressure (NPT) ensemble (see Electronic Supporting Information), then simulated at 300 K and 1.01325 bar for 10 ns recording coordinates at 5 ps intervals^96,97^. A representative frame was selected with lowest Cα RMSD to the average model of the last 1,000 frames of the trajectory (i.e., a set of frames with roughly constant Cα RMSD to the input model). This model was then minimized for 100 ps using the Desmond molecular dynamics energy minimization protocol. The protein model was extracted from the energy-minimized system and reprepared in the same fashion as the HMWP2-PCP1 model.

#### Cyclodehydration Intermediates

BioLuminate was used to model potential phosphopantetheinylated HMWP2-Cy2 cyclodehydration intermediates of *R* and *S* chiralities at the hydroxyl-bearing carbon center of the hydroxythiazolidine ((2*R/S*,4*R*)-2,2-hydroxy-(2’-(2”-hydroxyphenyl)-thiazolinyl)-thiazolidinyl phosphopantetheine)^72^. Note that the thiazolines of the 2-hydroxyphenyl-thiazolinyl (2HPT) side chains of these intermediates contain the chiral carbon corresponding to the d product of the l/d epimerization carried out by the upstream, embedded E domain of HMWP2^33^. The cyclodehydration intermediate models were prepared as singly charged anions (at the phosphate) using LigPrep in BioLuminate^72,98^. These molecules will be referred to as Ppant-2HPTT(*R/S*)-OH or *R* or *S* (cyclodehydration) intermediates.

### Molecular Docking

#### Protein-Protein Docking

The prepared HMWP2-PCP1 model was docked at the predicted opening of the upstream HMWP2-Cy2 active site tunnel (between strands β8 and β10 of the prepared model with all water molecules removed and the modified R1812 rotamer) using default settings in the HADDOCK2.4 web server^61–63,99^ with an additional, brief molecular dynamics simulation in explicit solvent. Active residues providing restraints were defined as the PCP phosphoserine (HMWP2 residue SEP1439) and Cy residue S1728, near the N terminus of helix α8 where the phosphopantetheine phosphate is predicted to be in PCP_donor_-Cy complexes (based on the PCP1_donor_ -C2 complex of LgrA, PDB ID 6MFX)^58^. The HMWP2-PCP2 homology model was docked similarly at the opening of the downstream active site tunnel of HMWP2-Cy2 (between helices α1 and α10), using Cy models with each possible combination of F1513 and R1812 crystallographic rotamers and the C-PCP-mimicking R1812 rotamer. For the results presented here, active residues were Cy F1513 and R1812 and PCP F1971, SEP1977, F2001, and the Cy model contained the F1513 rotamer B, directing the phenyl toward the PCP_acceptor_ binding site, and the C-PCP-mimicking R1812 rotamer. The additional active PCP2 residues F1971 and F2001 were included to promote favorable π-stacking interactions and/or hydrophobic packing with F1513 of the Cy domain, which adopts two rotameric states in the HMWP2-Cy2 crystal structure (one flipped out toward the PCP2 binding site and one away) and is present in over 40% of selected Cy domain sequences aligned in Fig. S7. Runs were also conducted with only R1812 and SEP1977 defined as active, but these and the other rotameric combinations of F1513 and R1812 did not perform as well, in some cases returning poses that resemble PDB ID 4ZXI or PDB ID 6N8E but that have considerably less favorable HADDOCK scores, lower buried surface areas and/or positive Z scores. These results are therefore not presented. In selecting docked poses, scores, energy terms and cluster sizes output by HADDOCK were considered, and the models were visually inspected for poses that were consistent with the HMWP2 modular architecture and reasonably placed the conserved PCP serine residues (SEP1439 or SEP1977) near the upstream and downstream active site tunnel entrances.

#### Cyclodehydration Intermediates

Because reasonable HMWP2-Cy2-PCP2 didomain complex models were obtained by protein-protein docking in HADDOCK2.4, the best-ranked of them was chosen as the basis for an approximate complex of the HMWP2-Cy2 crystal structure and the docked PCP2, to provide a Ppant phosphate site. The complex was formed by superimposing the docked HMWP2-Cy2-PCP2 complex on the HMWP2-Cy2 crystal structure using Cα alignment of Cy helices α1 and α10 (the secondary structure features forming most of the downstream tunnel entrance) followed by removal of the Cy domain from docking as well as redundant and conflicting interfacial water molecules and crystallographic water molecules in the tunnel that would block substrate access. The rotameric state of R1812 of the HMWP2-Cy2 structure was altered so its guanidinium moiety is positioned similarly to guanidinium moieties of conserved arginine residues observed in C-PCP_acceptor_ complexes. The resulting complex was then preprocessed and subjected to hydrogen bond network optimization and refinement of PCP2 and the interface waters and residues within 5 Å using the automatic method in Prime^69–71^. From this model a second was generated that contained an aspartic acid rather than aspartate at the putative catalytic dyad. Both models (i.e., anionic and neutral dyads) were then subjected to hydrogen coordinate refinement followed by all-atom energy minimization in Impact with a 0.3 Å heavy atom displacement cutoff preventing the models from deviating too much from their input coordinates.

Phosphoryl groups were then removed from the conserved PCP serine residues in preparation for covalent docking using Schrödinger’s Phosphonate_Addition.cdock reaction definition file available at https://www.schrodinger.com/products/covdock (also see Electronic Supporting Information). This reaction definition file, although designed for phosphonate reactions, is suitable for displacement of a phosphate oxygen by a serine side chain oxygen. The covalent docking tool in BioLuminate^72^ uses Glide docking^66–68^ in a rigid receptor approximation and returns only poses that are compatible with the covalent linkage the user defines^100^. Default covalent docking settings were used (maximum of 200 initial poses, keeping only those with Glide scores lower than +2.5, outputting the top 1,000 with 30 poses per ligand reaction site, which is 1 site in this case). The docking grid’s inner box, within which the ligand’s center must be positioned, was 10×10×10 Å^3^ and the outer box had dimensions 34×34×34 Å^3^. The center of the grid was defined as the centroid of tunnel-lining residues so that it was in the center of the tunnel’s diameter and positioned roughly halfway between R1812 and F1522, which lie at the ends of the docking region. Docking results were manually inspected to identify categories of hydroxythiazolidine orientation. Each category of pose for the *R* and *S* intermediates was subjected to optimization/refinement in the following scheme inspired by the refinement steps of Glide/Prime Induced Fit Docking protocols in BioLuminate^66–72,100–102^: 1) refinement of protein residues within 5 Å of the docked ligand using 10 iterations of 100 steps each with Prime’s automatic conjugate gradient/truncated Newton method, 2) refinement of the ligand and residues within 5 Å of it using 10 iterations of 100 steps each with Prime’s automatic method, 3) Monte Carlo protein side chain optimization for residues within 5 Å of the ligand (Predict Side Chains tool running Prime in BioLuminate), and 4) refinement of the ligand and residues within 5 Å of it using 10 iterations of 100 steps each with Prime’s automatic method. These refined models were sorted by Prime energy and evaluated using a combination of that value and the docking score. As a final point of comparison between poses, the phosphate-serine linkages of top covalent docking models were cleaved by deletion of the bridging oxygen (S1977 Oγ), replacing the broken bonds with bonds to hydrogens on the separated groups to leave a phosphonate and alanine, and MM-GBSA minimization calculations were performed on the resulting noncovalent complexes using Prime with 1 kcal/mol·Å^2^ restraints on flexible atoms, which were defined as those of the ligand and residues within 10 Å of it^69–71^.

### Low-Frequency Vibrational Normal Mode Analysis

The 1.94 Å-resolution structure of HMWP2-Cy2 (PDB ID 7JTJ) was retrieved from the RCSB PDB, and the missing loop and side chains were filled using Prime in BioLuminate^69–72^. Using the BioLuminate Protein Preparation Wizard^72,87–94^, hydrogen bond networks were optimized, hydrogen coordinates were refined to minimize potential energy, and all atom coordinates were refined to decrease potential energy with a cutoff before heavy atom RMSD reached 0.3 Å. The polyethylene glycol fragment and loosely bound surface water molecules were removed. The Desmond^96,97^ solvate pocket tool was used to add water molecules to the downstream portion of the active site tunnel where the polyethylene glycol fragment was removed (details available in Electronic Supporting Information). This model was placed in a solvent box with at least 10 Å separating the protein and the sides of the box, neutralizing with sodium ions and including 150 mM NaCl. The crystallographic sodium ion was replaced with a water molecule in this process. The default Desmond molecular dynamics^96,97^ (MD) relaxation protocol was run, and the output from the last stage with solute restraints (stage 5) was used as input to a series of 100 ps MD runs in the NPT ensemble at 277 K and 1.01325 bar, gradually releasing solute restraints as follows: 10 kcal/mol·Å^2^, 5 kcal/mol·Å^2^, then 1 kcal/mol·A^2^. These were followed with a series of 100 ps MD warming steps in 5 K increments from 280 K to 300 K, each in the NPT ensemble at 1.01325 bar. An additional 5.5 ns of NPT simulation at 300 K were performed recording frames every 1 ps. A representative frame was selected from the final 2,200 frames as the frame with the lowest RMSD to the average structure over this range. Surface water and ions were removed by selecting molecules more than 5 Å from the protein, then extending that selection to water or ions within 5 Å, leaving only the water most closely associated with the protein and the active site tunnel water undisturbed. An initial energy minimization in MacroModel^103^ using the Polak-Ribier Conjugate Gradient (PRCG) method with a cutoff of 0.0418 kJ/mol was performed and followed by normal mode calculation in BioLuminate^72^, returning 80 frames for each of the top 10 modes. This allowed for identification of loosely bound water molecules on the surface of the input model that contributed modes with negative eigenvalues. These water molecules were removed from the model used as input to the initial minimization, and the resulting model was subjected to PRCG energy minimization as before. Normal modes reported here were calculated for the minimized model without loosely bound surface water molecules.

### FTMap Ligand Fragment Binding Site Identification

The composite model built using the 1.94 Å- and 2.35 Å-resolution HMWP2-Cy2 structures was used as input to the FTMap server^74^. The entire protein was defined as the search space and default settings were used for binding site identification.

### Figure Preparation

Chemical figures were generated in ChemDraw (PerkinElmer), protein figures were generated using PyMOL (Schrödinger), and electrostatics figures were produced using the APBS PyMOL plugin with charges from the OPLS3e force field available in BioLuminate and van der Waals radii^72,92,104^. Calculations of approximate tunnel volumes were performed using the CASTp 3.0 web server available at the time of publication at http://sts.bioe.uic.edu/castp/calculation.html^105^. Sequence alignments were generated using Clustal Omega^106^. Representative Cy domain sequences were retrieved from UniProtKB by SANSparallel^107,108^, searching with seed Cy domain sequences from anguibactin, bacillamide, bacitracin, bleomycin, epothilone, JBIR-34/45, mycobactin, myxothiazole, pyochelin, vibriobactin, and yersiniabactin biosynthesis. Sequence redundancy was cut at 80% in JalView^109^ before alignment. WebLogo3^110^ was used to generate the sequence logo.

## Supporting information

Electronic Supporting Information Text, Tables and Figures

Low-Frequency Normal Mode Analysis Videos

## PDB Accession Numbers

The 1.94 Å- and 2.35 Å-resolution crystal structures for HMWP2-Cy2 have been deposited to the Protein Data Bank (http://www.rcsb.org/) and assigned the accession numbers 7JTJ and 7JUA, respectively.

## Acknowledgements

The authors kindly acknowledge Catherine L. Drennan for support in starting this project, the MIT X-ray facility and Staff Scientist Robert Grant for continued usage of the facility, and Subrata Mishra, Kenny Marincin, and Aswani Kancherla for enlightening discussions. Support for this research was provided by National Institute of General Medicinal Sciences (NIGMS) at the National Institutes of Health (NIH) (1R15GM123425-01) to D.P.D. A.D.G. was supported by a Sanofi Genzyme Doctoral Research Fellowship. B.C. was supported by an HHMI EXROP and EXROP Capstone Scholarship. This work is based upon research conducted at the Northeastern Collaborative Access Team beamlines, which are funded by the NIGMS (P30 GM124165). The Eiger 16M detector on 24-ID-E beam line is funded by a NIH-ORIP HEI grant (S10OD021527). D.F. acknowledges support from the National Institute of General Medicinal Sciences, NIGMS R01 GM104257. This research used resources of the Advanced Photon Source, a U.S. Department of Energy (DOE) Office of Science User Facility operated for the DOE Office of Science by Argonne National Laboratory under Contract No. DE-AC02-06CH11357. The content is solely the responsibility of the authors and does not necessarily represent the official views of the NIH.

