## Supplementary material for "Studies of a siderophore-producing cyclization domain: A refined proposal of substrate binding": Electronic Supporting Information Text, Tables and Figures

**Electronic Supporting Information for:**

**Files for computational work are available at https://github.com/adgnann/HMWP2-Cy2**

#### Contents

| | |
|---|---|
| **Supplemental Results** | **Page 1** |
| **Supplemental Discussion** | **Page 2** |
| **Supporting Information Table S1** | **Page 4** |
| **Supporting Information Figures** | **Page 5** |

### Supplemental Results

#### *Additional Cyclodehydration Intermediate Docking Observations*

Here we will address results for the docking of the *S* and *R* intermediates to HMWP2-Cy2 that are deemed less probable based on our scoring algorithm, which considers Glide docking scores, Prime energies and MM-GBSA energies, as well as sensibility in light of conservation trends and mutagenesis results in other systems. We will additionally discuss possible interactions at the donor side-chain binding region.

The two additional distinct conformational categories of *S* cyclodehydration intermediate poses (hydroxyl-toward-N-terminal-sheet and hydroxyl-toward-dyad) are initially intriguing because they also show favorable interactions with the Ppant arm and place reactive groups in the vicinity of conserved active site features (Figs. S18-19). However, it should be considered that the poses placing the hydroxyl toward the N-terminal β sheet would imply a substantial conformational change rotating the leaving group oxygen from its envisioned position during condensation (at the N-terminus of α4) through a large angle with concomitant rearrangement of the bulky 2HPT side chain and presumably of the Ppant arm as well. Similarly, the poses placing the leaving group oxygen toward the dyad also imply a large conformational change, and it is unclear where the cysteine thiol nucleophile would be sequestered during condensation if the cysteine amine is the initial nucleophile. With respect to the hydroxyl-toward-dyad category, one may envision how an initial thiol condensation with the amine nitrogen sequestered near T1856 could occur. One unique *S* intermediate pose was notable for its reference-like dimethyl positioning, rearrangement of Q1858 to form an interaction with the Ppant hydroxyl, and favorable interactions between N1751 and T1856 and the Ppant amide nearest the thioester and the thioester carbonyl, respectively. The 2HPT side chain is also deeper into the hydrophobic pocket in the side chain-binding region formed largely by F1522, M1629, F1840 and Y1866. This pose, however, also necessitates a sizable rotation of the hydroxythiazolidine and 2HPT side chain from the envisioned post-condensation state to the state it represents.

The other *R* intermediate categories may face the same complications as those of the *S* intermediates with leaving group oxygen atoms near the dyad or pointing toward the N-terminal subdomain β sheet. It appears that the *R* intermediate category of poses with the oxygen leaving group directed toward the dyad would, however, be obtainable from the envisioned condensation state with the cysteine amine as the initial nucleophile, suggesting the catalytic dyad could act as an acid protonating the leaving group in the cyclodehydration reaction. Poses of this category also direct the Ppant dimethyl moiety toward I1808, reminiscent of the orientation AB3403 (PDB ID 4ZXI) and FscG (PDB ID 7KW0) (Drake et al., 2016; Izore et al., 2021).

Poses of both intermediate diastereomers place the 2HPT side chain in the vicinity of the nonconserved residues F1522 and M1629 in the side chain-binding region. For the *R* intermediate, this arrangement of its side chain and F1522 could permit M1629 to pack against their phenyl rings, which could offer a specific aromatic-methionine-aromatic interaction that would help select the cognate substrate. The S poses are positioned so that it is nearly possible for staggered parallel stacking between the 2HPT phenyl and F1522, and this positioning is compatible with interactions between the 2HPT hydroxyl and N1621, although it should be noted that this residue is a phenylalanine in the second Cy domain of pyochelin biosynthesis. In the *R* poses, the N1621 carboxamide is often stacked on the thiazoline ring of the 2HPT side chain, sometimes forming a polar contact with S1854. The positioning of N1621 in many of the *S* poses places it in direct contact with the leaving group oxygen, a situation that seems unlikely since this residue is not conserved and its position in the sequence is often occupied by a hydrophobic residue.

#### Supplemental Discussion

##### *Tandem Cy Domains*

Studies of nonribosomal peptide synthetases with tandem Cy domains have shown that condensation can in some cases be followed by a product release step and rebinding in a cyclodehydration-competent state (i.e., in another domain) rather than immediate cyclodehydration (Katsuyama et al., 2021; Marshall et al., 2002). In the case of the tandem Cy domains in FmoA2 and FmoA3 (Katsuyama et al., 2021), for instance, the Cy domain of FmoA2 does not actually produce cyclized product and instead acts only as a condensation domain. This system then requires FmoA3-Cy to catalyze cyclodehydration of the FmoA2-Cy condensation product. Interestingly, FmoA3-Cy has also lost its ability to catalyze peptide formation. Katsuyama et al. (Katsuyama et al., 2021) point out that this loss of peptide formation in FmoA3-Cy and other Cy domains of tandem Cy systems could stem from increased catalytic demands in the ring closure step that are at odds with requirements for condensation (e.g., blocking bulk solvent with a phenylalanine). In VibF, the absence of the capping phenylalanine at the upstream tunnel entrance is a feature of the Cy domain that is only condensation competent (Marshall et al., 2002), whereas in AngN the Cy domain missing this residue is apparently still cyclodehydration competent, as evidenced by low-level production of anguibactin in a mutant in which the other Cy domain should be catalytically inactivated by mutation of its DXXXXD motif (Di Lorenzo et al., 2008).

By dividing the condensation and cyclodehydration reactions into separate domains, evolution would be able to optimize the domains separately. This would be especially salient when the nucleophile is a hydroxyl, as it is in Fmo and Vib Cy reactions, rather than a thiol. By contrast, Cy domains using cysteine to form thiazoline in non-tandem contexts would benefit from cyclodehydration without condensation intermediate release, and evolution would therefore balance the apparent tradeoff between competence for condensation and competence for cyclodehydration. The Cy domain structures from HMWP2, EpoB and BmdB are examples that natively condense and cyclize cysteine, and since HMWP2-Cy2 accepts the cyclodehydration intermediate in docking, it is now possible to envision what conformational changes around the active site would permit elongation of the cyclized S-C bond toward a pre-cyclodehydration state consistent with expectations for donor substrate binding based on LgrA-C2 (PDB ID 6MFX) (Reimer et al., 2019).

##### *Catalysis*

Considering the structural rearrangements likely to accompany the transition between open-for-condensation and closed-for-cyclodehydration states, it seems plausible that the Ppant positioning relative to features of the downstream tunnel entrance in observed C-$PCP_{acceptor}$ complexes may be physiologically relevant, whereas the proximity of the Ppant thiol to the N-terminus of helix α4—too near to accommodate acceptor and donor substrates in between—may be an artifact due to the lack of substrates and represent a state primed for catalysis upon association of a loaded donor PCP.

It remains unclear in the C domain literature to what extent the histidine at the second position of the HHXXXD motif is required for catalysis, with its mutation perhaps complemented by other basic groups in some cases (Keating et al., 2002; Roche and Walsh, 2003). It may be that a proton of a nucleophilic amine can be shuttled away through a hydrogen bond network to the mostly conserved floor loop aspartate (HMWP2-Cy2 D1770) or even that this aspartate reaches further into the active site to act as base during catalysis of that step. The barrier to reaction may also just be low enough to not require a carefully coordinated deprotonation. Alternatively, the cysteine thiol, being an intrinsically better nucleophile, may act in the condensation step. This is an interesting possibility because it might explain differences in polar contacts to the Ppant in the downstream tunnel that could result in sequestration of the amine near the aspartate-threonine dyad and presentation of the thiol(ate) as the nucleophile for condensation. One line of reasoning may work against this idea though. In the region of negative charge surrounding the active site, namely from D1862 and D1770, the thiolate state of the cysteine side chain would be less favorable than in solution, perhaps even necessitating a coordinated deprotonation event as would also be envisioned for deprotonation of the neutral amine nucleophile after attack.

Given the high degree of flexibility observed in the C domain family, exemplified by our coauthor's recent NMR results (Mishra et al., 2021), the correct question may instead be *to what extent are the amine-first and thiol-first pathways traversed?* With such diversity among Cy domains, the field should remain open to the possibility that different systems rely on reduction of substrate conformational heterogeneity, electrostatic arrangement and proton shuttling to different extents, reminiscent of the variable effects that mutagenesis experiments have had on conserved residue positions in C and Cy domains (Bloudoff and Schmeing, 2017).

#### Supporting Information Table

**Table S1 – Data collection and refinement statistics.**

| | | |
|---|---|---|
| ***Data Collection*** | | |
| | **HMWP2-Cy2 at 1.94 Å resolution** | **HMWP2-Cy2 at 2.35 Å resolution** |
| **Detector** | Saturn 944+ | ADSC Quantum 315.1 |
| **Space group** | $P4_12_12$ | $P4_12_12$ |
| **a,b,c** | 89.29, 89.29, 140.07 | 89.32, 89.32, 140.04 |
| **Resolution range (Å)*** | 30.8 - 1.94 (1.99 – 1.94) | 50.0 - 2.35 (2.43 – 2.35) |
| **Completeness** | 98.9 (96.1) | 99.3 (99.3) |
| **Redundancy** | 16.449 (9.418) | 11.8 (12.1) |
| **Unique Reflections** | 41892 (2957) | 24196 (2349) |
| **I/σI** | 28.25 (3.41) | 19.2 (2.94) |
| $R_{sym}$ **#** | 0.077 (0.735) | 0.142 (0.853) |
| $R_{pim}$ **$** | 0.019 (0.236) | 0.043 (0.252) |
| **CC1/2 &** | 1.00 (0.848) | 0.994 (0.851) |
| ***Refinement*** | | |
| **Resolution range (Å)** | 30.8 – 1.94 | 23.33 – 2.35 |
| $R_{free}$ **^** | 0.2167 | 0.2117 |
| $R_{work}$ **^** | 0.1799 | 0.1717 |
| **Reflections/test set** | 41875/2074 | 24158/1200 |
| **Number of atoms** | | |
| **Protein** | 3836 | 3477 |
| **Water** | 355 | 227 |
| **Sodium and PEG** | 9 | 9 |
| **B-factors (Å$^2$)** | 32.80 | 38.33 |
| **Protein** | 32.17 | 38.23 |
| **Water** | 38.29 | 39.31 |
| **Sodium** | 27.30 | 35.80 |
| **PEG** | 60.02 | 57.09 |
| **RMS Deviations** | | |
| **Bonds (Å)** | 0.008 | 0.003 |
| **Angles (°)** | 0.89 | 0.56 |
| **Dihedrals (°)** | 17.75 | 17.20 |
| **Ramachandran Analysis†** | | |
| **Favored (%)** | 98.81 | 99.05 |
| **Allowed (%)** | 1.19 | 0.47 |
| **Outliers (%)** | 0.0 | 0.47 |

† Ramachandran analysis calculated using MolProbity (Williams et al., 2018).

* Highest resolution shell is shown in parentheses

\# $R_{sym/merge} = \Sigma_{hkl}\Sigma_i|I_i(hkl) - \langle I(hkl)\rangle|/\Sigma_{hkl}\Sigma_i I_i(hkl)$.

$ $R_{pim} = \Sigma_{hkl}[1/(N-1)]^{1/2}\Sigma_i|I_i(hkl) - \langle I(hkl)\rangle|/\Sigma_{hkl}\Sigma_i I_i(hkl)$ where $I_i(hkl)$, $\langle I(hkl)\rangle$ and N represent the intensity measurement, the mean intensity, and the redundancy for reflection hkl, respectively.

& $CC^* = [2CC_{1/2}/(1+CC_{1/2})]^{1/2}$ where $CC_{1/2}$ is the correlation between two random halves of the datasets, each containing half of the measured intensities for each unique reflection and CC* is an approximation of the correlation coefficient for a noise-free dataset.

^ $R_{work} = \Sigma|F_{obs}(hkl) - F_{calc}(hkl)|/\Sigma|F_{obs}(hkl)|$, where $F_{obs}(hkl)$ and $F_{calc}(hkl)$ are the observed and calculated structure factor amplitudes of ~95% of the reflections used for refinement. $R_{free}$ was calculated from the ~5% of total reflections that were omitted from the refinement.

#### Supporting Information Figures


| Figure | Page |
|---|---|
| Figure S1 - (methyl)ox-/thiazol((id)ine) natural products. | 6 |
| Figure S2 - A mechanistic proposal for reactions in the Cy domain. | 7 |
| Figure S3 – Secondary structure assignment in HMWP2-Cy2. | 8 |
| Figure S4 – Size exclusion chromatography verifying the monomeric state of HMWP2-Cy2. | 9 |
| Figure S5 – Tunnel representations and solvent-accessible measurements from the CASTp 3.0 server for three Cy domain crystal structures. | 10 |
| Figure S6 – Crystallographic sodium site in the active site tunnel at the interface of strands β1 (N-terminal subdomain) and β11 (C-terminal subdomain). | 11 |
| Figure S7 – Annotated sequence logo representing 1040 Cy domain sequences. | 12 |
| Figure S8 – Alignment of the DXXXXD(XXS) motif of sequences used in SANSparallel to search Uniprot KB. | 13 |
| Figure S9 – Required displacements at the upstream tunnel entrance of HMWP2-Cy2 (cyan) to achieve an open state. | 14 |
| Figure S10 – Variation in loops of the downstream tunnels of Cy domains and representative C-PCP$_{acceptor}$ models. | 15 |
| Figure S11 – Comparison of downstream tunnel entrances in Cy domains. | 16 |
| Figure S12 – Comparison of the top HMWP2-PCP2 docking model to the Cy-PCP complex from PchE (PDB ID 7EN1). | 17 |
| Figure S13 – Comparison of the top HMWP2-PCP2 docking model to the C-PCP complex from AB3403 (PDB ID 4ZXI). | 18 |
| Figure S14 – Comparison of the top HMWP2-PCP2 docking model to the C-PCP complex from ObiF1 (PDB ID 6N8E). | 19 |
| Figure S15 – Comparison of the top HMWP2-PCP2 docking model to the C-PCP complex from FscG (PDB ID 7KW0). | 20 |
| Figure S16 – Superimposition of C domain strand and helix Cα atoms in C-PCP$_{acceptor}$ models used in evaluating protein-protein docking results. | 21 |
| Figure S17 – Cyclodehydration intermediate models used in covalent docking experiments. | 22 |
| Figure S18 – Hydroxyl-toward-α4 Ppant-2HPTT(*S*)-OH poses from covalent docking. | 23 |
| Figure S19 – Hydroxyl-toward-N-terminal-sheet Ppant-2HPTT(*S*)-OH poses from covalent docking. | 24 |
| Figure S20 – Ppant-2HPTT(*S*)-OH covalent docking poses in hydroxyl-toward-dyad or deeply placed hydroxyl-toward-N-terminal-sheet orientations. | 25 |
| Figure S21 – Hydroxyl-toward-α4 Ppant-2HPTT(*R*)-OH poses from covalent docking. | 26 |
| Figure S22 – Hydroxyl-toward-N-terminal-sheet Ppant-2HPTT(*R*)-OH poses from covalent docking. | 27 |
| Figure S23 – Hydroxyl-toward-dyad Ppant-2HPTT(*R*)-OH poses from covalent docking. | 28 |
| Figure S24 – Comparison of possible pantetheine interactions in the downstream tunnels of Cy domains against observed C domain pantetheine interactions. | 29 |
| Figure S25 – Geometric rationale for a proposed pre-condensation state shared by C and Cy domains. | 30 |
| Figure S26 – Comparison of the top cyclodehydration intermediate pose from HMWP2-Cy2 and the product-bound state reported for PchE-Cy. | 31 |
| Figure S27 – The high-amplitude HMWP2-Cy2 low-frequency normal mode number 7. | 32 |
| Figure S28 – Change in inter-residue distances with the top 10 low-frequency normal vibrational modes of HMWP2-Cy2. | 33 |
| Figure S29 – FTmap server results for HMWP2-Cy2. | 34 |
| Figure S30 – The HMWP2-Cy2 active site binds small organic molecules in FTmap docking. | 35 |
| Figure S31 – Proposals for a global Cy domain conformational change associated with catalysis and three main reaction steps. | 36 |

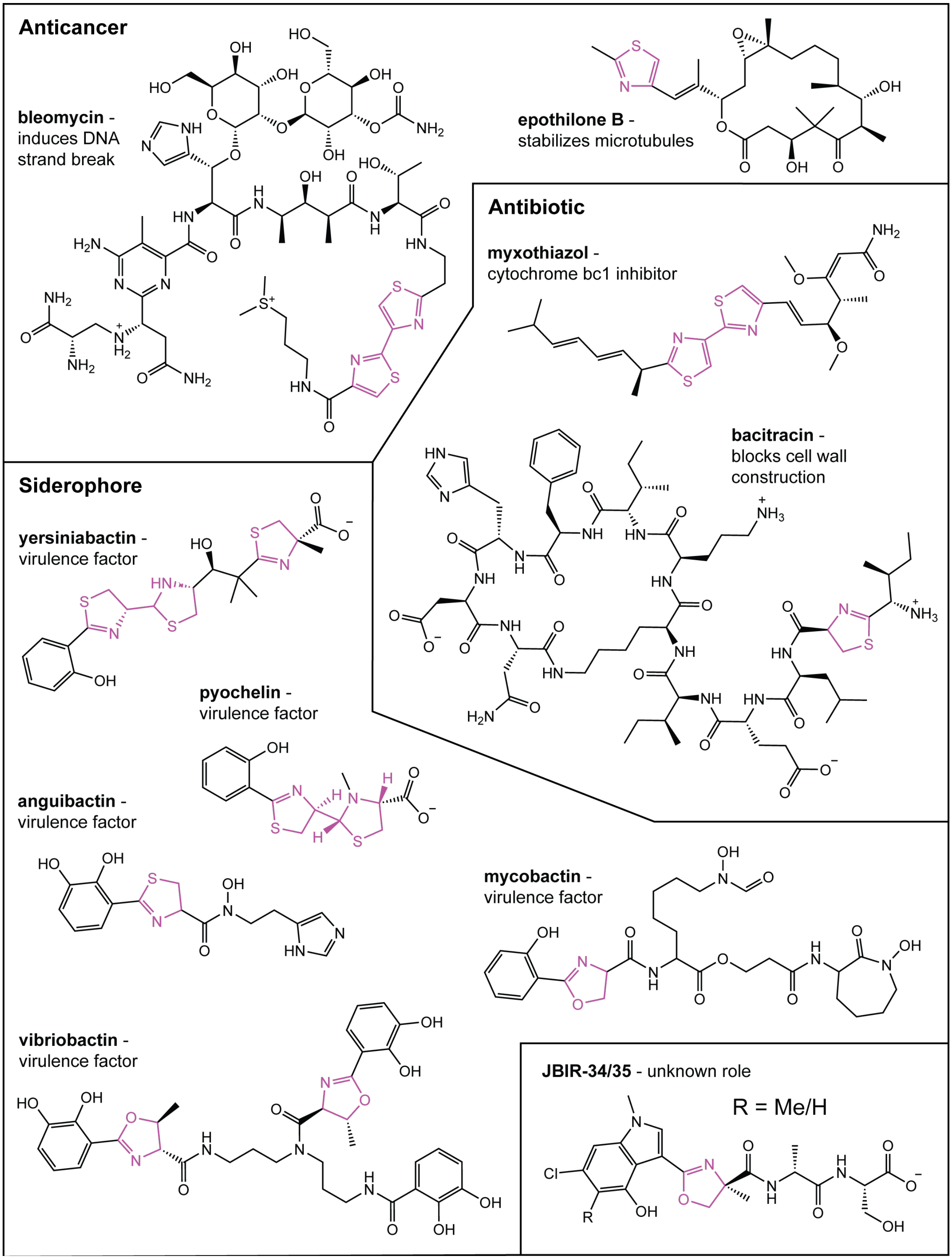


**Figure S1 - (methyl)ox-/thiazol((id)ine) natural products.**

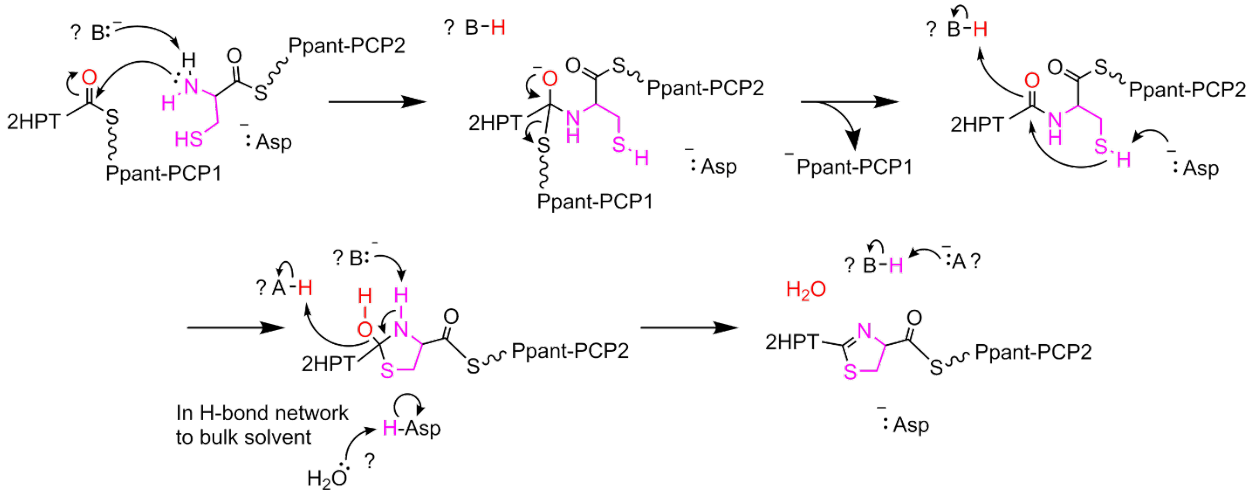


**Figure S2 - A mechanistic proposal for reactions in the Cy domain.** The oxygen incorporated in the water leaving group is colored red. The acceptor cysteine side chain and amine N-H incorporated into the condensation product are colored magenta. Some outstanding questions regarding catalysis by Cy domains involve the nature of proton transfer steps. Unknown species invoked to participate in these steps are indicated by question marks. The possibility of the first step involving a cysteine thiolate nucleophile instead of the amine nucleophile cannot be ruled out at this time, but it is not shown here.

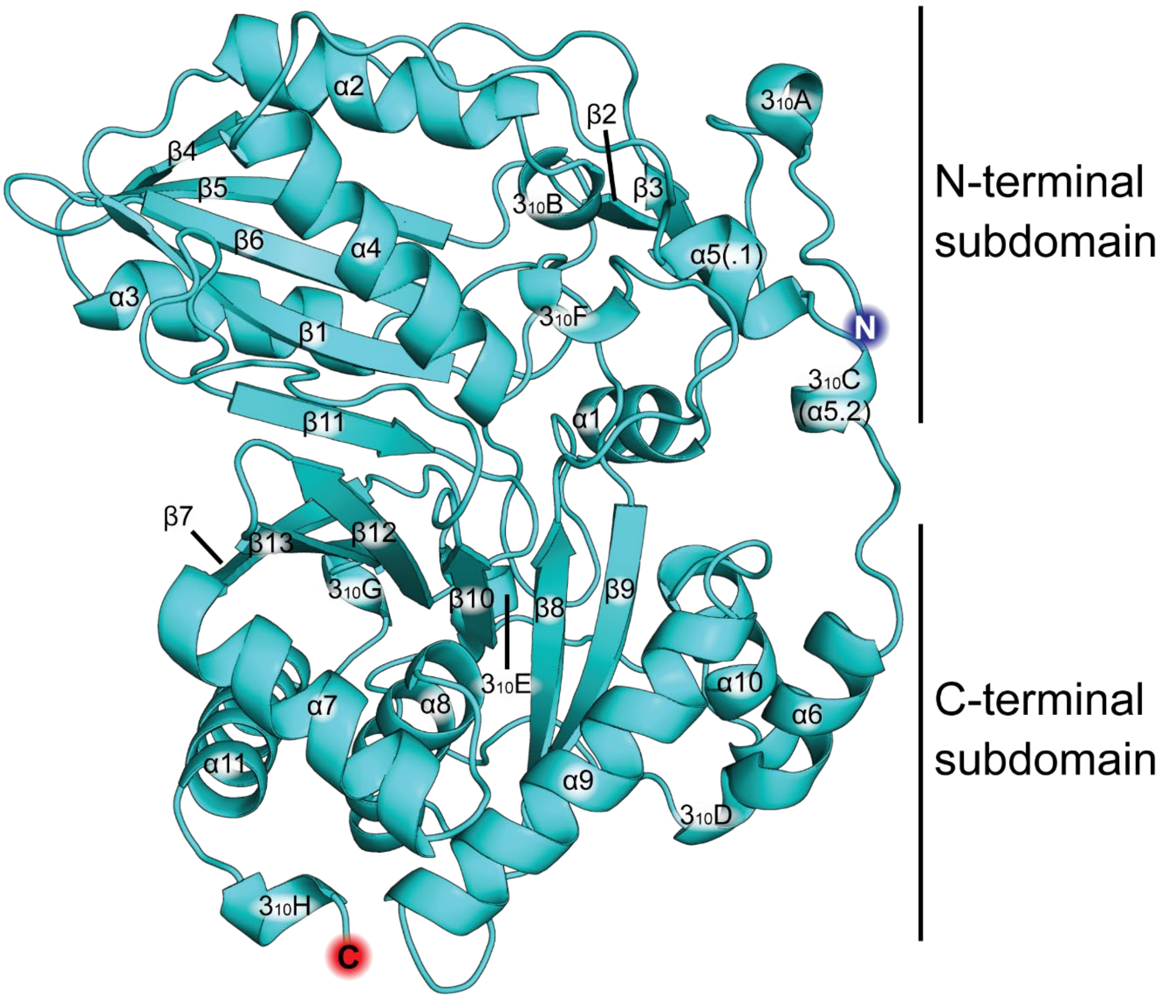


**Figure S3 – Secondary structure assignment in HMWP2-Cy2.** The model depicted here primarily contains coordinates from the higher resolution structure with the exception that residues missing from that structure in the region between $3_{10}$C and α6 (HMWP2 1664-1665 and 1667-1668) are modeled using coordinates from the lower resolution structure. Prime in BioLuminate was used to model P1666, the final missing residue in that region. The N- and C-terminal chloramphenicol acetyltransferase-like subdomains are labeled, and the N- and C termini are marked with blue and red spheres, respectively. $3_{10}$ helices identified by STRIDE (Frishman and Argos, 1995) are displayed in helix cartoon representations, and they are defined to include all residues bearing a hydrogen bond donor/acceptor interacting within the $3_{10}$ helix as identified by manual inspection, which extends the STRIDE prediction by 1 residue in some cases (these definitions are reflected in Fig. S7).

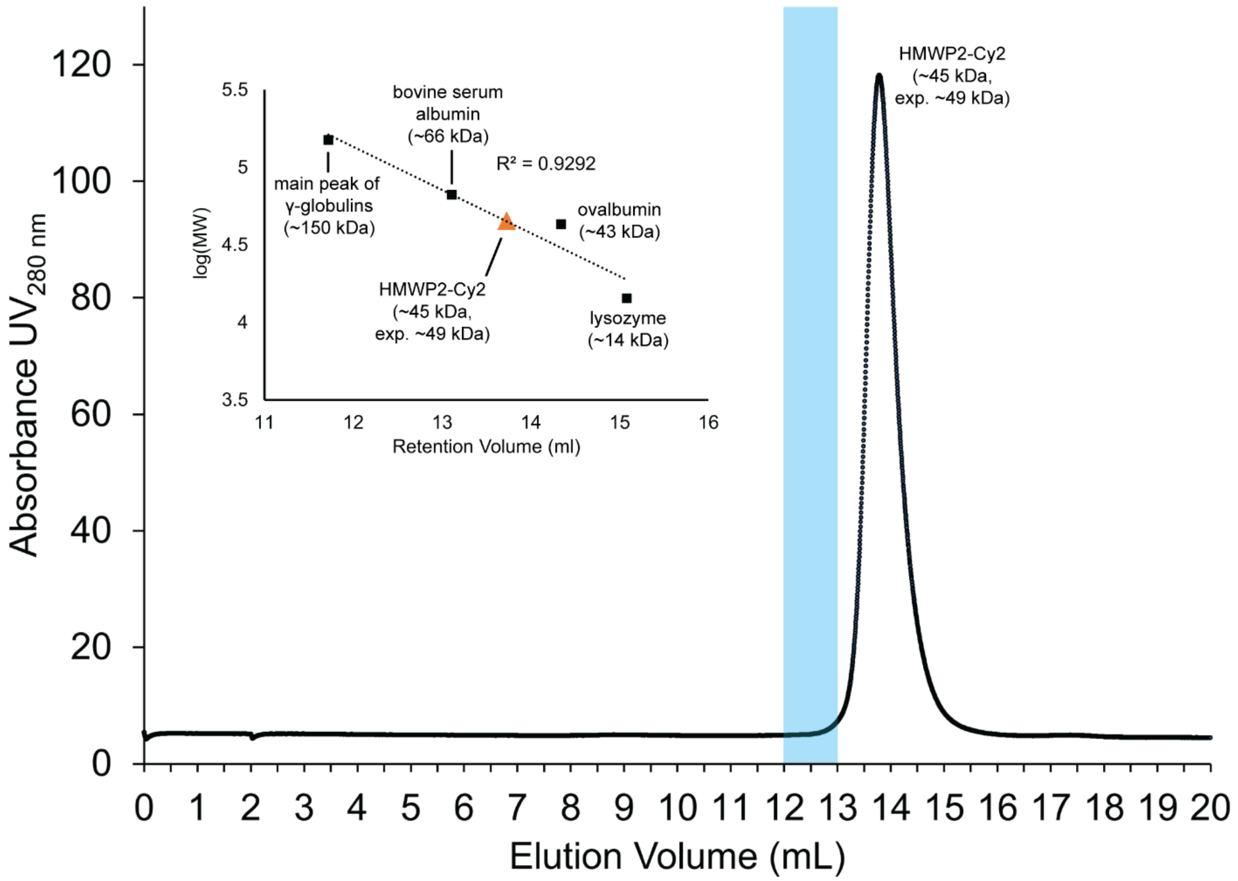


**Figure S4 – Size exclusion chromatography verifying the monomeric state of HMWP2-Cy2.** The main plot of absorbance at 280 nm versus elution volume shows a single, homogenous peak for HMWP2-Cy2 corresponding to a molecular weight just lower than expected (~45 kDa versus the expected ~49 kDa). The blue stripe centered on 12.5 mL indicates the volume around which a dimer of HMWP2-Cy2 would be expected to elute. No feature is observed in this region. The inset panel shows the regression curve fitting the logarithm of molecular weight versus retention volume for four protein standards, as well as an orange triangle marker for HMWP2-Cy2 and the $R^2$ value of the fit, 0.9292.

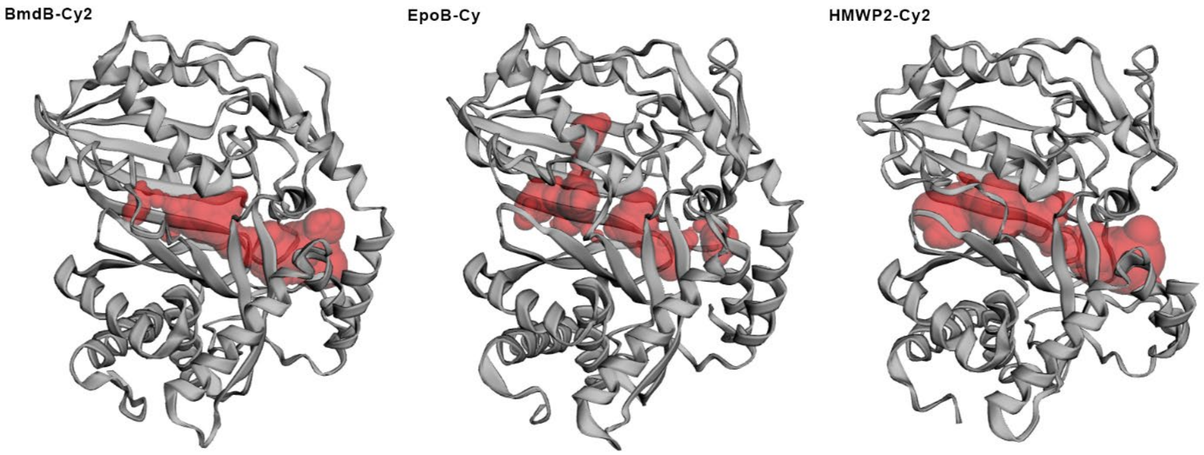


BmdB-Cy2

| Pocket ID | SA Surface Area (Å²) | SA Volume (Å³) |
|---|---|---|
| 1 | 625.675 | 382.242 |

EpoB-Cy

| Pocket ID | SA Surface Area (Å²) | SA Volume (Å³) |
|---|---|---|
| 2 | 316.777 | 105.019 |
| 4 | 179.873 | 73.007 |
| 9 | 60.498 | 20.570 |

HMWP2-Cy2

| Pocket ID | SA Surface Area (Å²) | SA Volume (Å³) |
|---|---|---|
| 1 | 774.503 | 543.174 |

**Figure S5 – Tunnel representations and solvent-accessible measurements from the CASTp 3.0 server for three Cy domain crystal structures.** These panels are taken from output in the graphical user interface of CASTp 3.0. Note that the tunnel in the EpoB-Cy crystal is formed by a composite of three tunnel definitions, indicating that the tunnel is discontinuous to a probe of water radius (1.4 Å) in this model. The sum of the EpoB-Cy volumes is 198.6 $Å^2$.

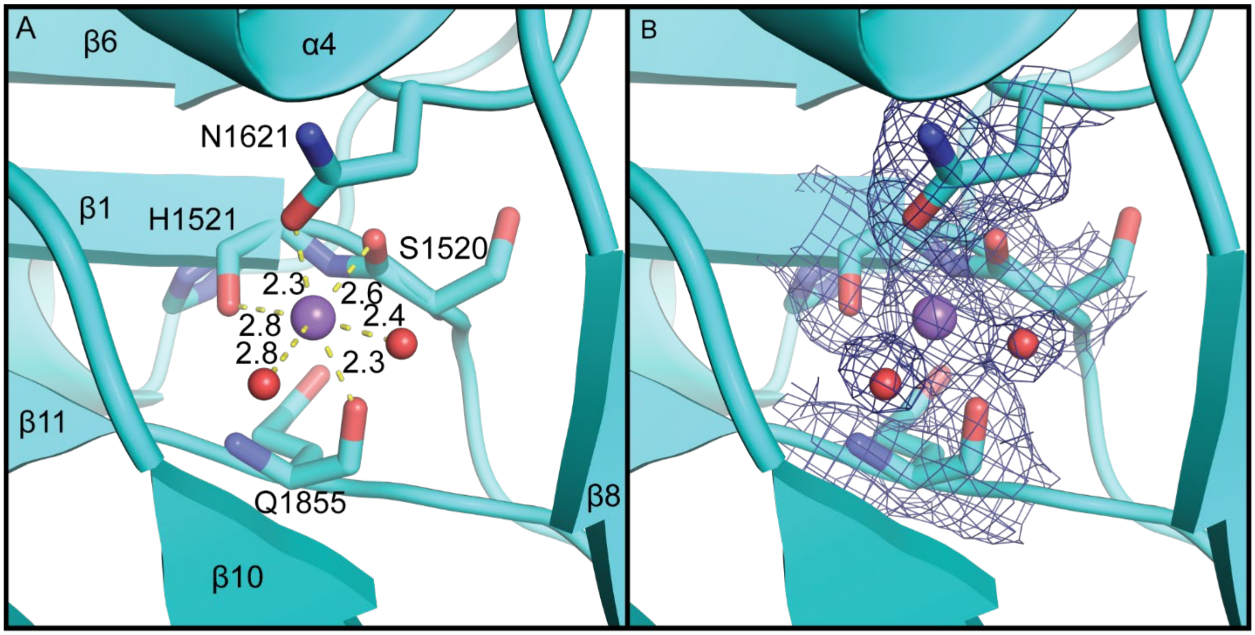


**Figure S6 – Crystallographic sodium site in the active site tunnel at the interface of strands β1 (N-terminal subdomain) and β11 (C-terminal subdomain). A**, Distances between oxygen ligands and the sodium ion are labeled in Å (yellow dashed lines). **B**, composite omit $2mF_o$-$DF_c$ electron density map contoured at 1 σ. Mesh is displayed within 2 Å of S1520, H1521 and Q1855 backbone atoms, N1621 side chain atoms and the two displayed water oxygen atoms.

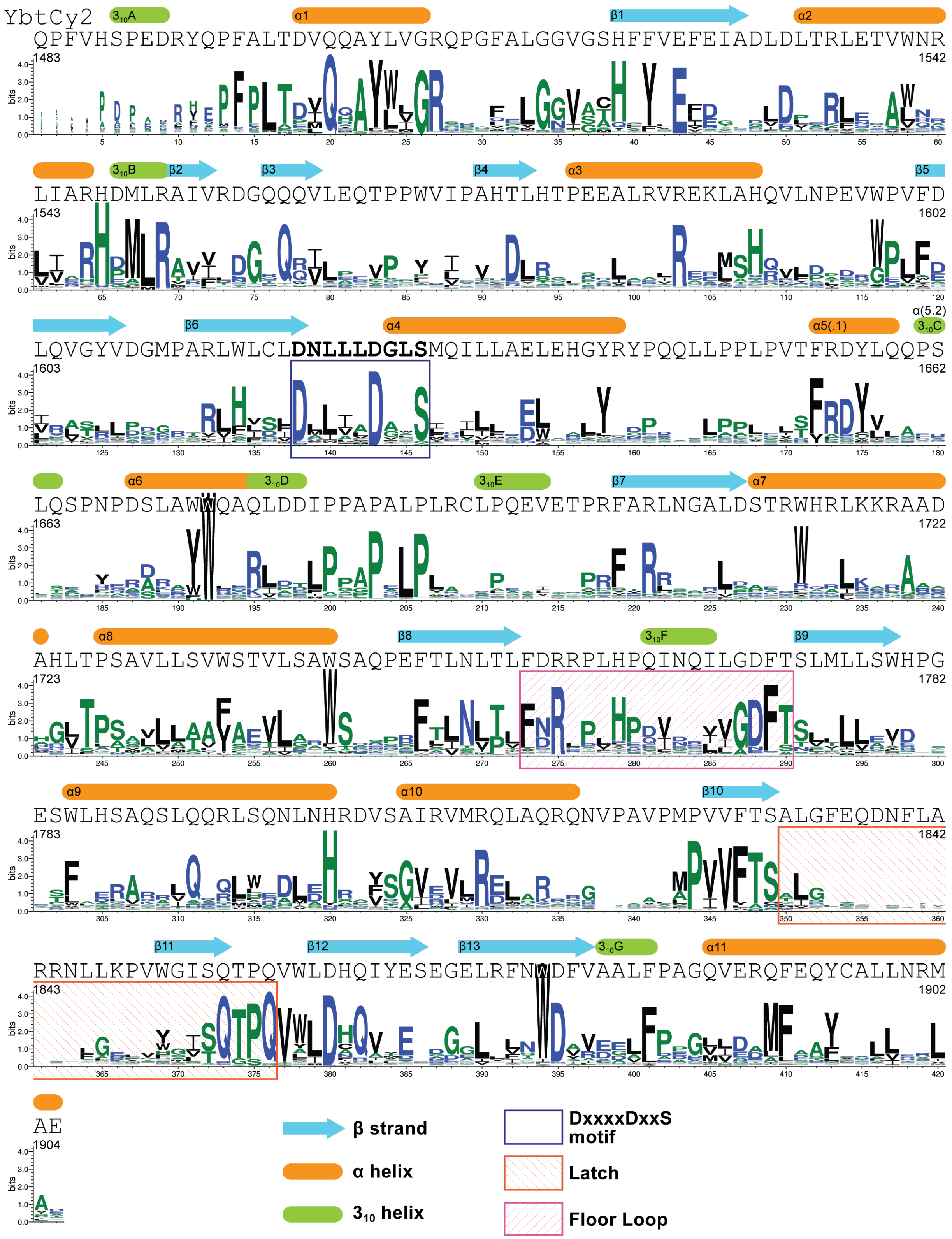


**Figure S7 – Annotated sequence logo representing 1040 Cy domain sequences.** The sequence of HMWP2-Cy2 is aligned above the logo. Secondary structure annotations correspond to features identified by STRIDE (Frishman and Argos, 1995) in the HMWP2-Cy2 model, and $3_{10}$ helix definitions match those in Fig. S3. Seed sequences used to search UniProtKB with SANSparallel (Koskinen and Holm, 2012; Somervuo and Holm, 2015) were from anguibactin, bacillamide, bacitracin, bleomycin, epothilone, JBIR-34/35, mycobactin, myxothiazole, pyochelin, vibriobactin, and yersiniabactin biosynthesis. JalView (Waterhouse et al, 2009) was used to perform an 80% redundancy cut of sequences longer than 400 amino acids, providing 1040 sequences for alignment by Clustal Omega (Sievers et al., 2011). The sequence logo in this figure was generated using WebLogo3 (Crooks et al., 2004).

| | | |
|---|---|---|
| HMWP2-Cy2 | DNLLLDGLS | |
| BlmIV-Cy1 | DALICDAHS | 42.72 |
| BlmIV-Cy2 | DLLIADAHS | 41.29 |
| MtaC-Cy1 | DAITADASA | 39.68 |
| HMWP1-Cy3 | DLLQFDVQS | 39.23 |
| MtaD-Cy2 | DLLTADAFS | 39.04 |
| PchF-Cy2 | DFTLVDYAS | 38.86 |
| PchE-Cy1 | DLLAADVES | 38.28 |
| BacA1-Cy1 | DPLICDDSS | 37.83 |
| HMWP2-Cy1 | DLLIMDASS | 36.87 |
| FmoA3-Cy | DLQLMDASS | 35.78 |
| BmdB-Cy2 | DALLMDGAS | 35.20 |
| EpoB-Cy | DLINVDLGS | 35.03 |
| MbtB-Cy1 | DMQAADAMS | 32.57 |
| VibF-Cy1* | DMIACDAQS | 27.44 |
| AngN-Cy1 | DMIAIDPDS | 25.88 |
| VibF-Cy2† | DALIVDGRT | 25.81 |
| AngN-Cy2 | DALILDARS | 21.99 |

**Figure S8 – Alignment of the DXXXXD(XXS) motif of sequences used in SANSparallel to search UniProtKB.** The sequences are sorted by identity to HMWP2-Cy2 (percent ID is in the right column). Yellow squares indicate Cy domains using cysteine, red squares indicate Cy domains using hydroxyl-bearing acceptors. Blue boxes mark the conserved positions of the motif. The black brackets in the bottom left indicate pairs of tandem Cy domains.

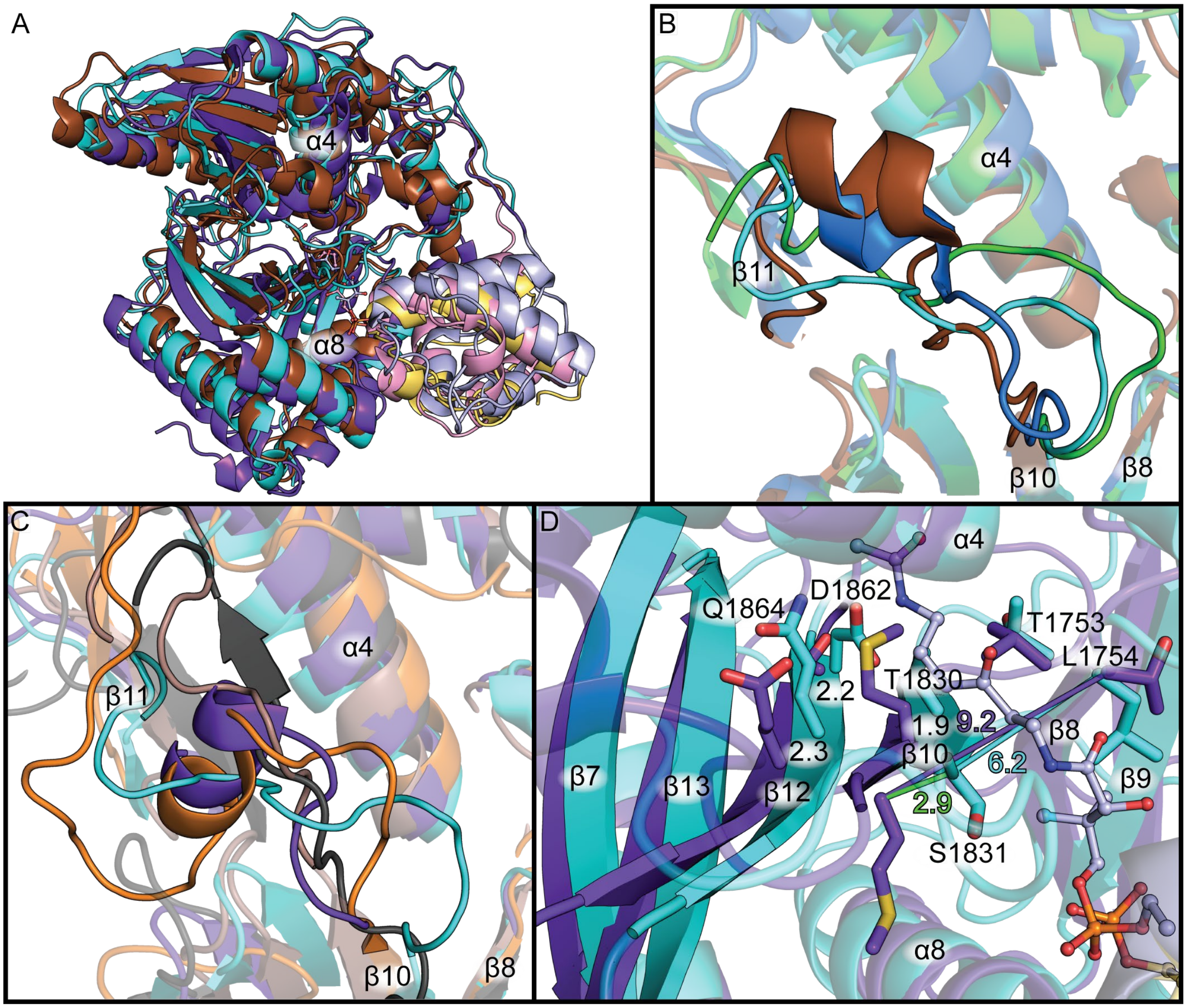


**Figure S9 – Required displacements at the upstream tunnel entrance of HMWP2-Cy2 (cyan) to achieve an open state. A**, A superimposition of the protein-protein docking model of HMWP2-PCP1-Cy2 (PCP yellow, Cy cyan; see also Fig. 3A-D) with LgrA-C (C purple, PCP light blue, Ppant analog light blue ball and stick, PDB ID: 6MFX) and PchE-Cy (Cy brown, PCP pink, Ppant pink ball and stick, PDB ID: 7EN1). **B**, A superimposition of Cy domain structures with the latch loop in opaque cartoon (EpoB-Cy is green, HMWP2-Cy2 is cyan, BmdB-Cy2 is blue, and PchE-Cy is brown). PDB IDs are EpoB-Cy: 5T7Z, BmdB-Cy2: 5T3E, PchE-Cy: 7EN1. **C**, A superimposition of C domain structures and HMWP2-Cy2 (cyan) with the latch loop in opaque cartoon (AB3403-C is orange, LgrA-C is purple, ObiF1-C is black). PDB IDs are AB3403-C: 4ZXI, LgrA-C: 6MFX, ObiF1-C: 6N8E. **D**, The gap between strand β8 leading into the floor loop and strand β10 leading into the latch would expand by nearly 3 Å (from ~6.2 Å in HMWP2-Cy2 to 9.2 Å in LgrA-C) to accommodate binding of the donor pantetheine. Approximately 2 Å also separate the Cα atoms of the Cy catalytic dyad from the Cα atoms of their counterparts in LgrA-C, reflecting that the entire lobe of the C-terminal subdomain sheet containing β-strands 7, 13, 12, and 10 is rotated.

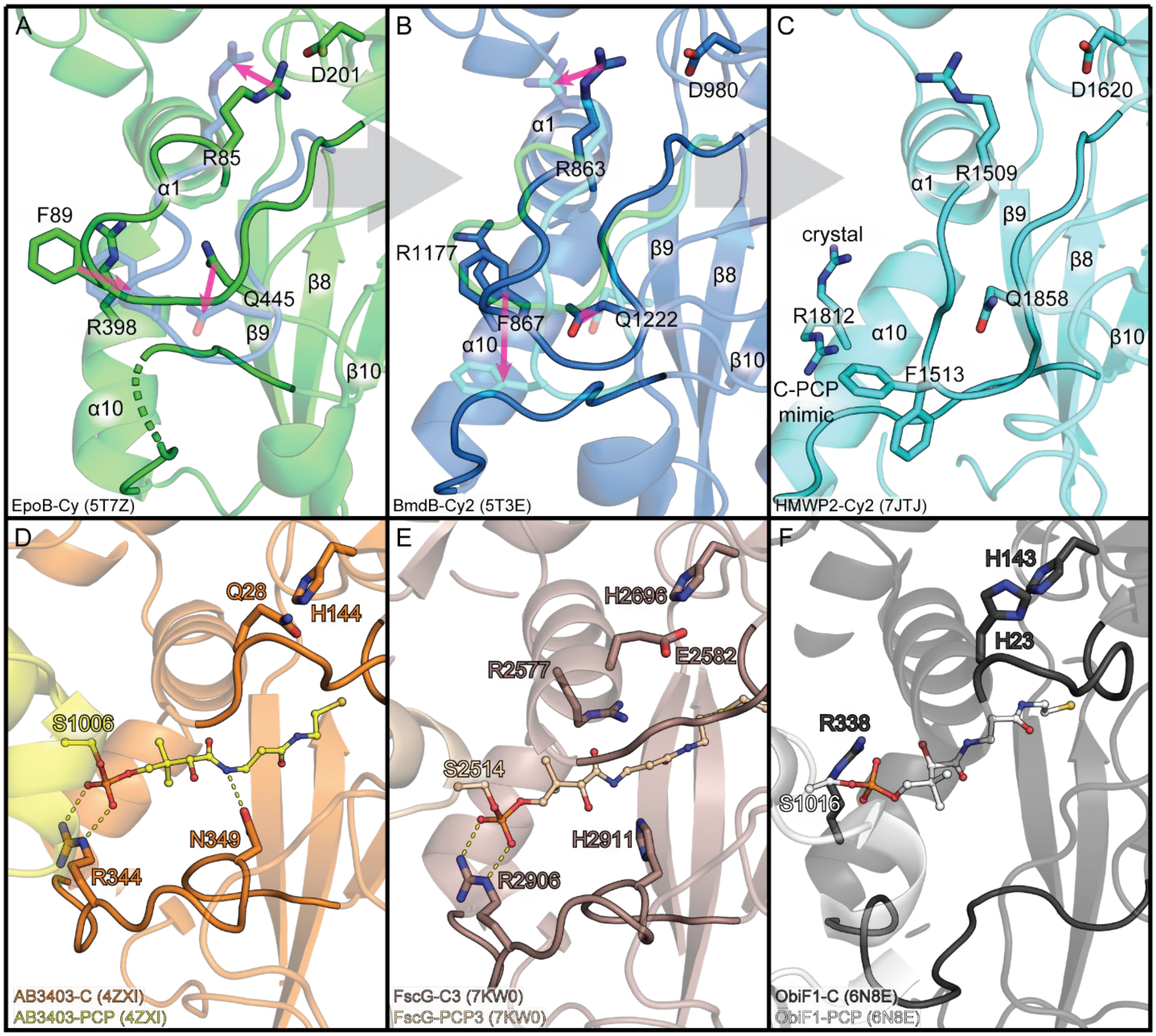


**Figure S10 – Variation in loops of the downstream tunnels of Cy domains and representative C-PCP$_{acceptor}$ models.** The system/name and PDB ID of each model is listed in the bottom left corner of each panel. **A**, EpoB-Cy is displayed in green. The blue, transparent cartoon/stick overlay shows the position of loop 1 in BmdB-Cy2 for direct comparison. Pink arrows highlight the movements of loop 1 residues displayed in sticks. **B**, BmdB-Cy2 is displayed in blue. The cyan and green transparent cartoon/stick overlays show the positions of loop 1 in EpoB-Cy and HMWP2-Cy2 for direct comparison, and pink arrows highlight movements of loop 1 residues between BmdB-Cy2 coordinates and HMWP2-Cy2 coordinates. **C**, The downstream tunnel entrance state in HMWP2-Cy is shown. Both crystallographic F1513 rotamers are displayed, and the alternate R1812 rotamer used in docking as well as the crystallographic R1812 rotamer are shown. **D**, The C-PCP$_{acceptor}$ model from AB3403 is displayed, showing the R344-S1006 (phosphate) interaction that inspired testing the alternate HMWP2-Cy2 R1812 rotamer, and a single hydrogen bond between N349 and the pantetheine (yellow ball and stick representation). The visible secondary structure shares the same numbering as the Cy domains in panels A-C. **E and F**, The same view as panel D is shown, but for FscG and ObiF1. Whereas FscG-C3 offers a potential polar contact to pantetheine at H2911, ObiF1 shows no polar contact to pantetheine in this position. Additionally, ObiF1 demonstrates a different downstream tunnel entrance arginine configuration that resembles the Cy domains (R338) and the pantetheine is positioned differently so that the thiol is not as deep into the active site region and the dimethyl moiety is rotated around the axis of the pantetheine by approximately 180° relative to AB3403 and FscG models.

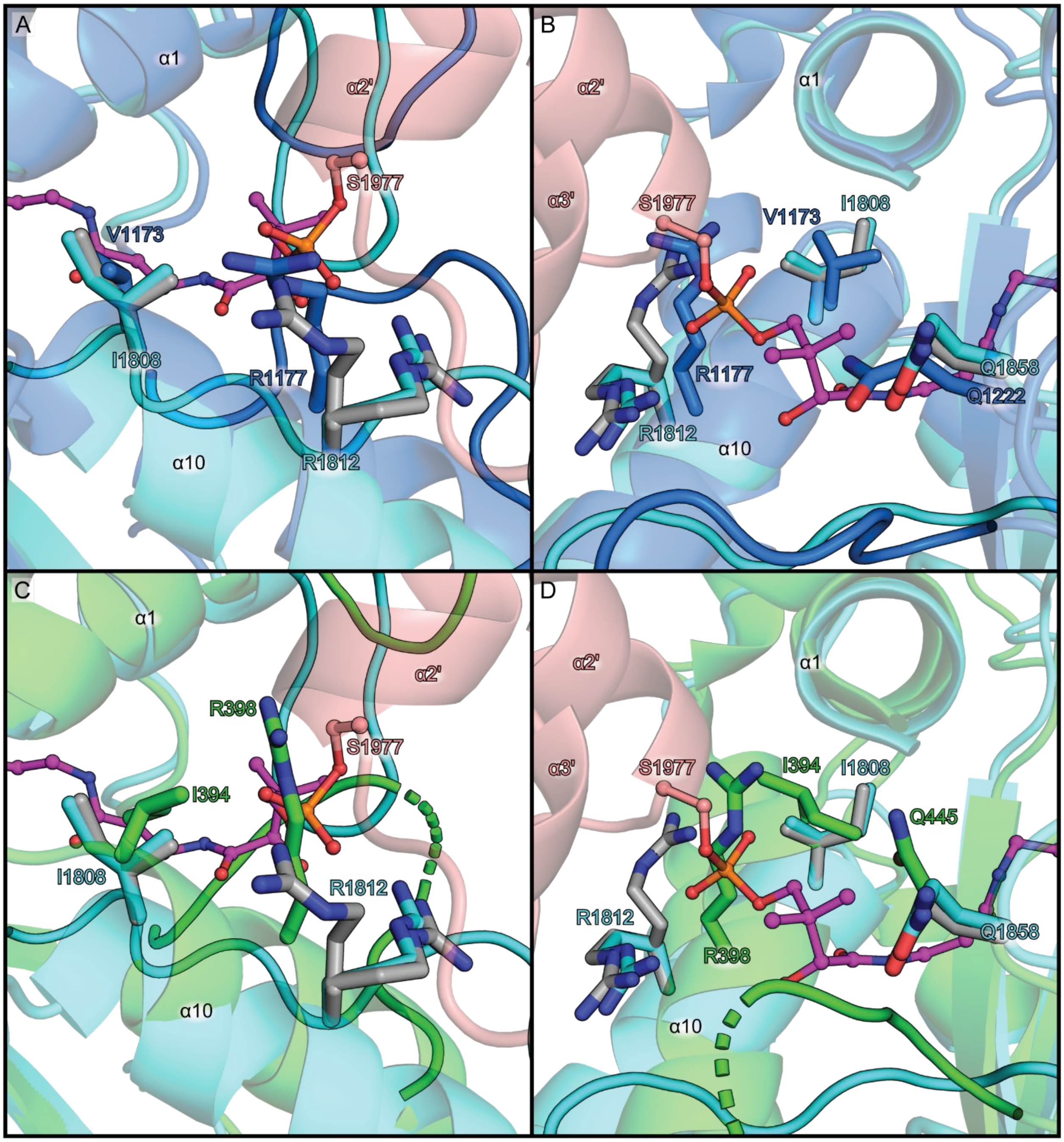


**Figure S11 – Comparison of downstream tunnel entrances in Cy domains. A and B**, BmdB-Cy2 (blue) is superimposed on the docked HMWP2-Cy2/HMWP2-PCP model (cyan/salmon) with the top *R* cyclodehydration intermediate pose (magenta ball and stick, described further in the results section on cyclodehydration intermediate docking and Figs. 4 and S17-23). Gray sticks are HMWP2-Cy2 crystal structure coordinates or the docking input model (R1812 alternate conformation). Non-transparent cartoon loops are the loops following α1 and α10. Panel B is approximately related to panel A by a 140° rotation around the Y axis. Note that both the crystallographic and C-PCP$_{\text{acceptor}}$-based HMWP2-R1812 rotamers are displayed in panel B. Also, the conserved glutamines (HMWP2-Q1858 and BmdB-Q1222) are in different rotameric states. **C and D**, The same views are shown as in panels A and B, but for comparison of HMWP2-Cy2 and EpoB-Cy (green). In HMWP2-Cy2, EpoB-Cy and BmdB-Cy2, a small hydrophobic residue (I/V) is found near the N-terminus of helix α10 in the vicinity of the pantetheine dimethyl groups of the C-PCP models from AB3403 and FscG (PDB IDs 4ZXI or 7KW0, respectively), not ObiF1 (PDB ID 6N8E, see panels D and E in Fig. S10 and panels C and D in Fig. S13-15).

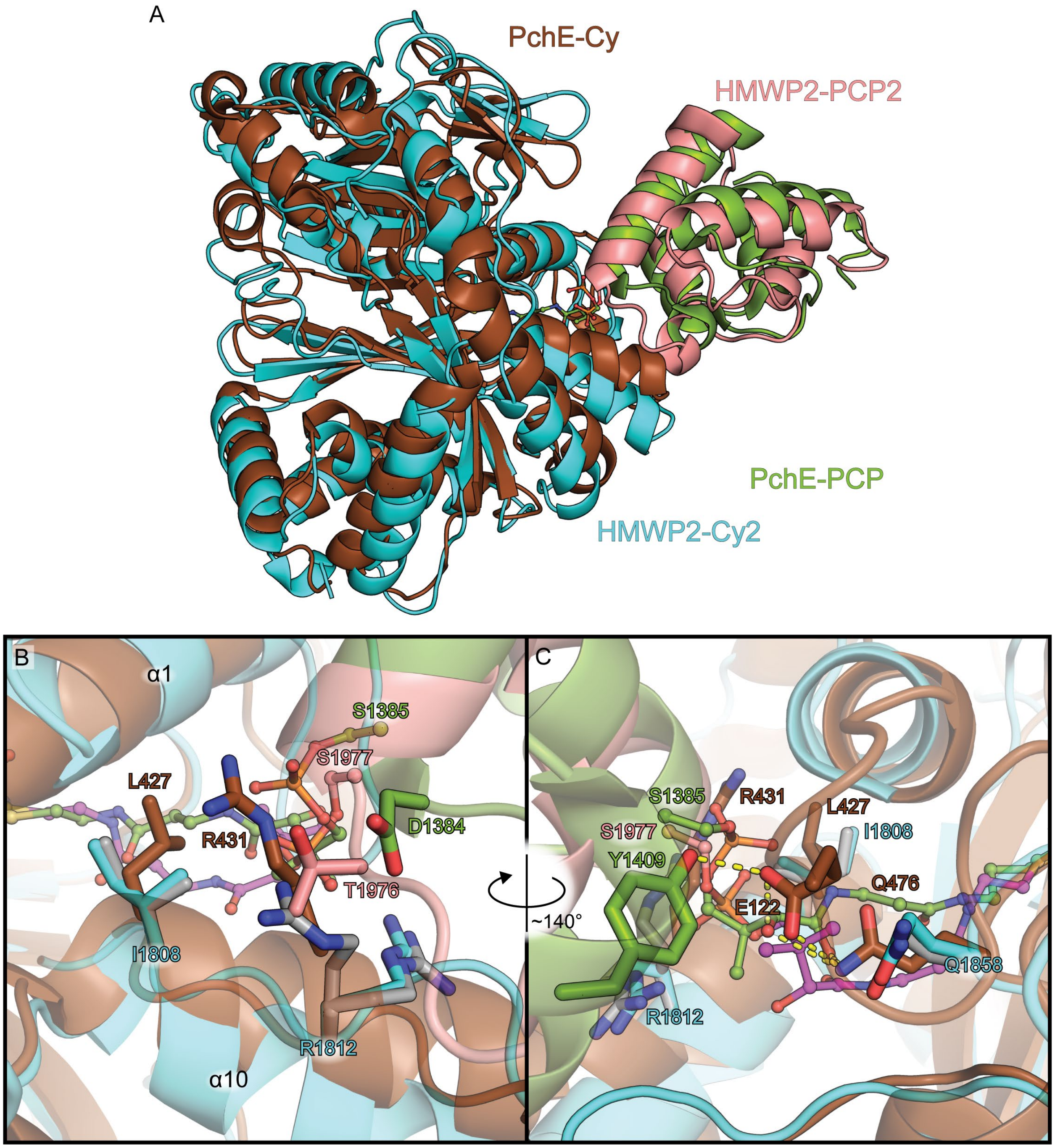


**Figure S12 – Comparison of the top HMWP2-PCP2 docking model to the Cy-PCP complex from PchE (PDB ID 7EN1). A**, The positioning of the HMWP2-Cy2-PCP2 model resembles the recent PchE-Cy product-bound complex (Wang et al., 2022), with the most notable difference in secondary structure being rotation of α10 around its N-terminus in PchE, bringing its C-terminus toward the PCP. **B,C**, Unlike the top HMWP2-Cy2-PCP2 complex, the PchE-Cy structure places R431 in a rotameric state like the crystal structure of HMWP2-Cy2, and E122 in loop 1 permits a hydrogen bonding network involving the conserved Cy residue Q476, pantetheine and Y1409 of the acceptor PCP that is not possible in the Cy systems crystallized to date due to lack of conservation of E122 and Y1409. An additional arginine, absent in HMWP2-Cy2, is found at the C-terminus of α10 and interacting with Y1409 in PchE but is omitted from this figure (HMWP2-Cy2 is cyan, HMWP2-PCP2 is salmon, PchE-Cy is brown and PchE-PCP is green).

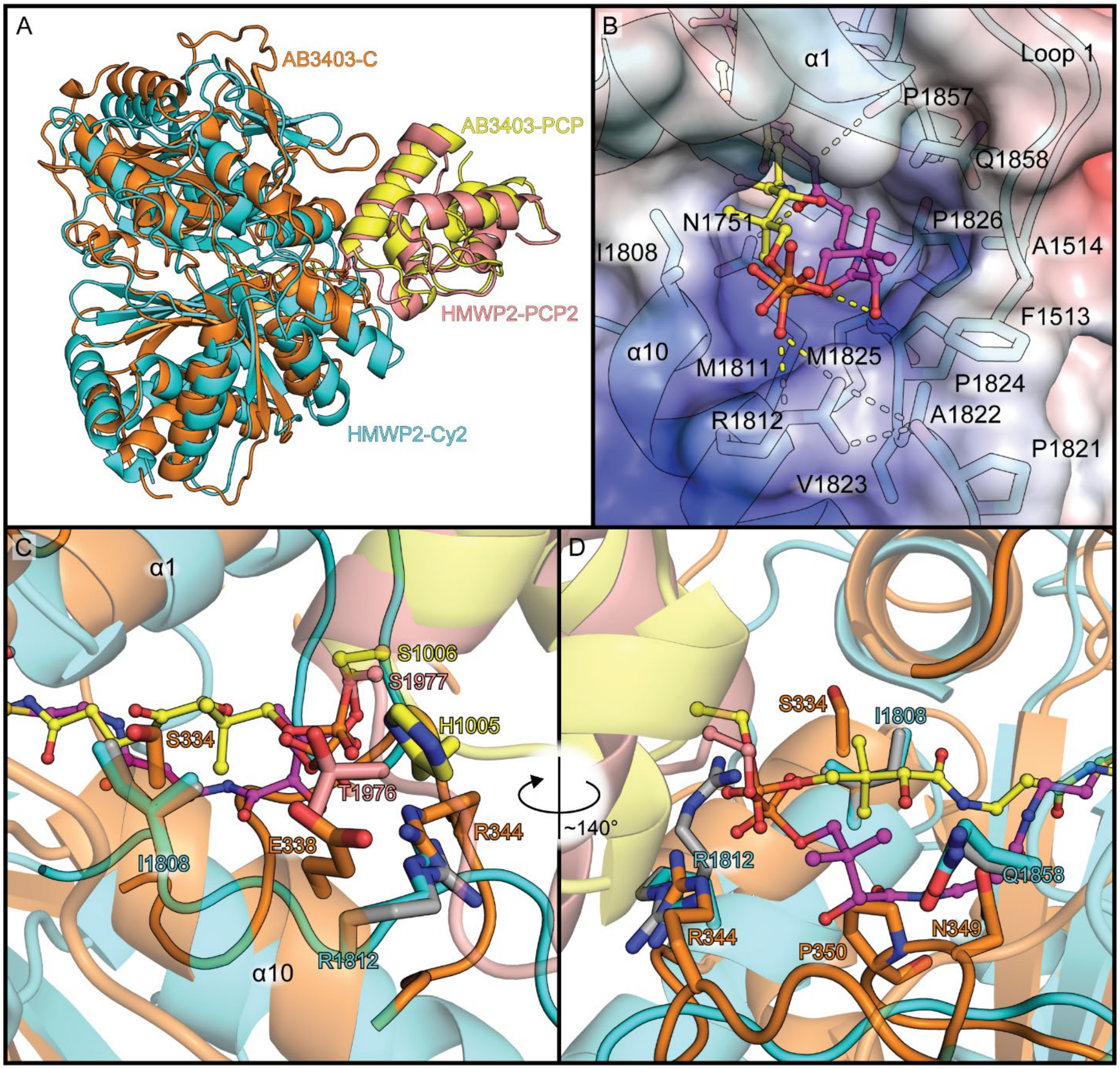


**Figure S13 – Comparison of the top HMWP2-PCP2 docking model to the C-PCP complex from AB3403 (PDB ID 4ZXI). A**, Fairly close correspondence between the HMWP2 (docked) and AB3403 (crystallographic) complexes can be seen in a superimposition of C/Cy domain strand and helix Cα atoms. **B**, The electrostatic potential surface of the top HMWP2-Cy2 Ppant-2HPT(*R*)T-OH is displayed over HMWP2-Cy2 sticks and cartoon. The docked Ppant pose from this work is in magenta, and the reference Ppant model from the AB3403 complex is in yellow. **C**, Unlike the HMWP2-Cy2-PCP2 complex, the AB3403 complex includes an interaction between H1005 and E338 across the C-PCP interface. Additionally, whereas Cy domain structures compared here have small hydrophobic residues near the N-terminus of helix α10, AB3403 has a serine at this position, suggesting the polar/hydrophobic character of this position may not bear as greatly on Ppant placement (i.e. dimethyl orientation) as the size of the side chain at this position). Gray sticks are HMWP2-Cy2 crystallographic coordinates, and coloration otherwise matches panel A. **D**, The view of the downstream tunnel entrance is shown rotated approximately 140° around the Y axis relative to panel C.

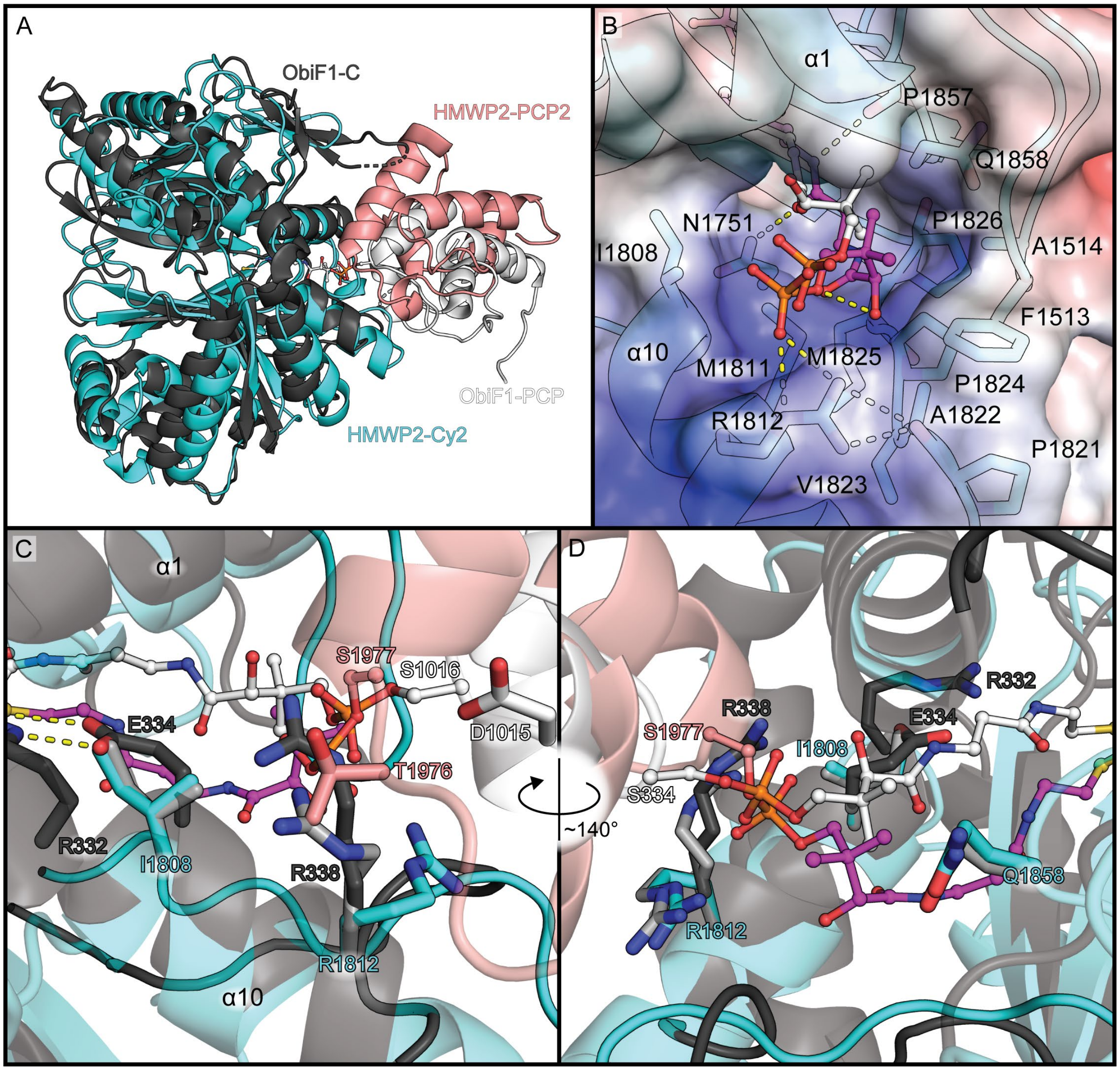


**Figure S14 – Comparison of the top HMWP2-PCP2 docking model to the C-PCP complex from ObiF1 (PDB ID 6N8E). A**, The PCP domains of HMWP2 and ObiF1 do not superimpose as well as HMWP2 and AB3403 (PDB ID 4ZXI, Fig. S13A). **B**, The alternate Ppant placement (white) observed in ObiF1 (relative to AB3403) more closely resembles the top Ppant-2HPT(*R*)T-OH pose (purple). The same electrostatic potential surface and underlying HMWP2-Cy2 sticks/cartoon are displayed as in Fig. S12B. **C**, A main difference between ObiF1 and the other C and Cy domains compared here is E334, a bulkier, charged residue toward the N-terminus of helix α10, which forms a salt bridge with R332. **D**, No specific pantetheine interactions are observed in ObiF1 (unlike in the other C models and the HMWP2 docking model).

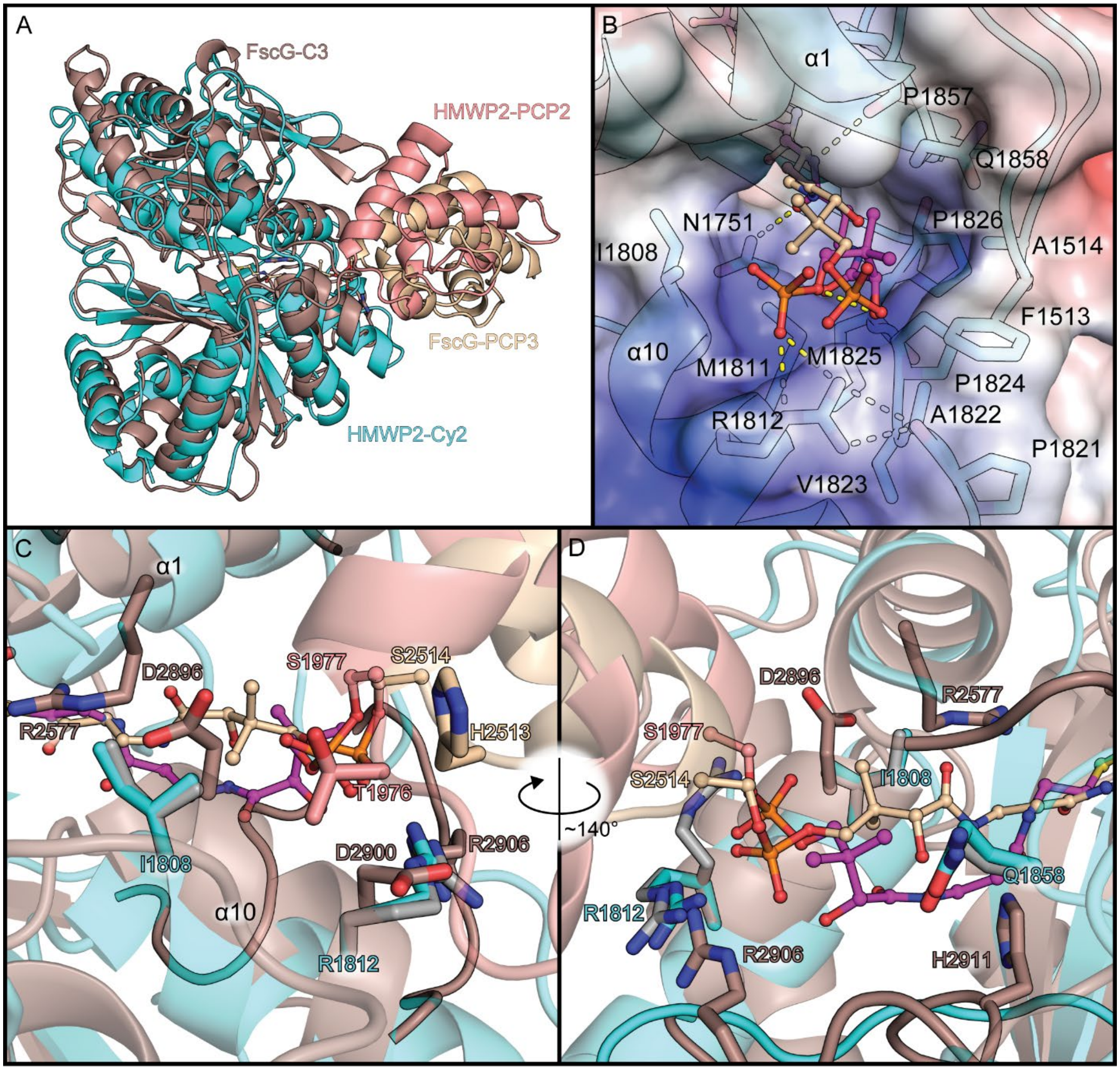


**Figure S15 – Comparison of the top HMWP2-PCP2 docking model to the C-PCP complex from FscG (PDB ID 7KW0). A**, The PCP domains of HMWP2 and FscG do not superimpose as well as HMWP2 and AB3403 (PDB ID 4ZXI, Fig. S13A), but the FscG pose seems to be intermediate between the AB3403 and ObiF1 (see Fig. S12). **B**, A Ppant binding mode like that of the AB3403 complex is observed in the FscG model, which is interesting to note given that the FscG-PCP and AB3403-PCP orientations differ considerably. It should be noted that the FscG model harbors a stabilized Ppant-glycine substrate mimic which is thought to resemble a condensation acceptor-bound state. **C and D**, Although the C-PCP$_{acceptor}$ orientation in FscG differs from the orientation in AB3403, the positioning of Ppant in the downstream tunnel of each model is similar. The difference in C-PCP$_{acceptor}$ orientation may be related to the increased distance between the ion pair H2513-D2900 in FscG. It is interesting to consider the relationship between D2896 and R2577 as it relates to the residue pair R332-E334 in ObiF1 in which the arginine does not form an interaction with Ppant, unlike R2577 in FscG, which forms weak electrostatic interactions with oxygen atoms of the Ppant amides.

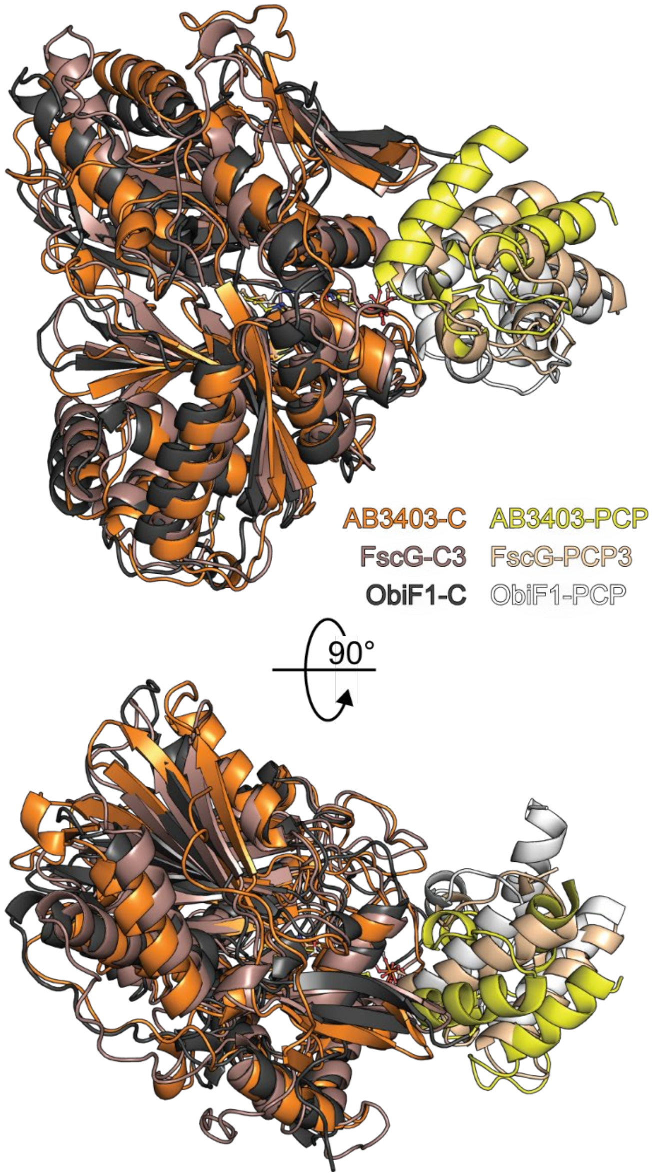


**Figure S16 – Superimposition of C domain strand and helix Cα atoms in C-PCP$_{acceptor}$ models used in evaluating protein-protein docking results.** The colors of each component are labeled in the bottom left of the top part of the figure. The top part of the figure shows a "side-on" view of the complexes, and the bottom part of the figure shows a "top-down" view (viewed from N-terminal subdomain). Notice that the helical bundles of the PCP domains are shifted relative to each other. The C-PCP placement in AB3403 most closely resembles the top protein-protein docking result presented here (Fig. 3E-H). (PDB IDs: 4ZXI – AB3403, 7KW0 – FscG, 6N8E – ObiF1).

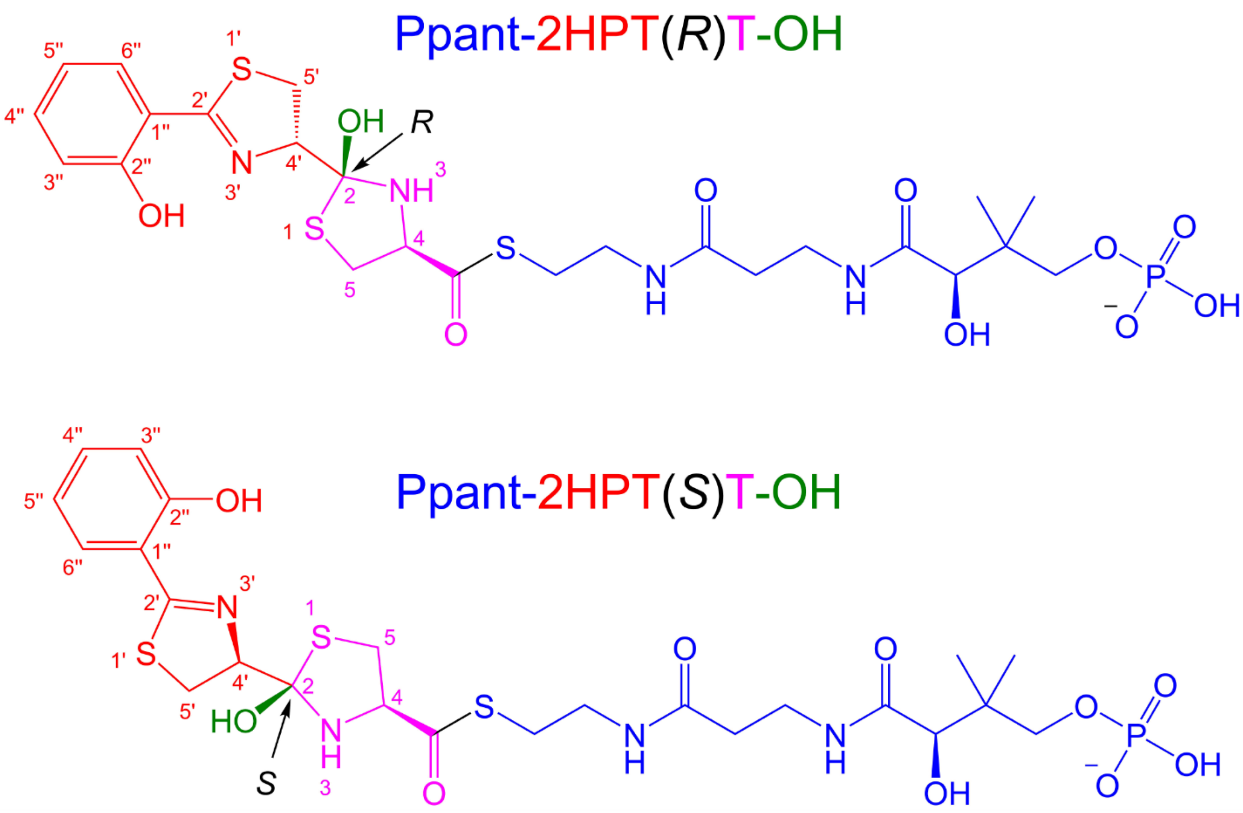


**Figure S17 – Cyclodehydration intermediate models used in covalent docking experiments.** Structures of both chiralities of cyclodehydration intermediates tested here are color coded by moiety. The 2HPT side chain is in red, the hydroxyl (water leaving group) is in green, the thiazolidine and carbonyl derived from cysteine in HMWP2-Cy2 is in magenta, and the phosphopantetheine is in blue. The structures differ only by their configuration at the 2 position of the reactive hydroxythiazolidine moiety. The structures are displayed in such a way that the orientation of the side chain roughly resembles its orientation in the top covalent docking results for each intermediate (the hydroxyl-toward-α4 classes of pose). Ring numbering uses primes (') to differentiate between rings.

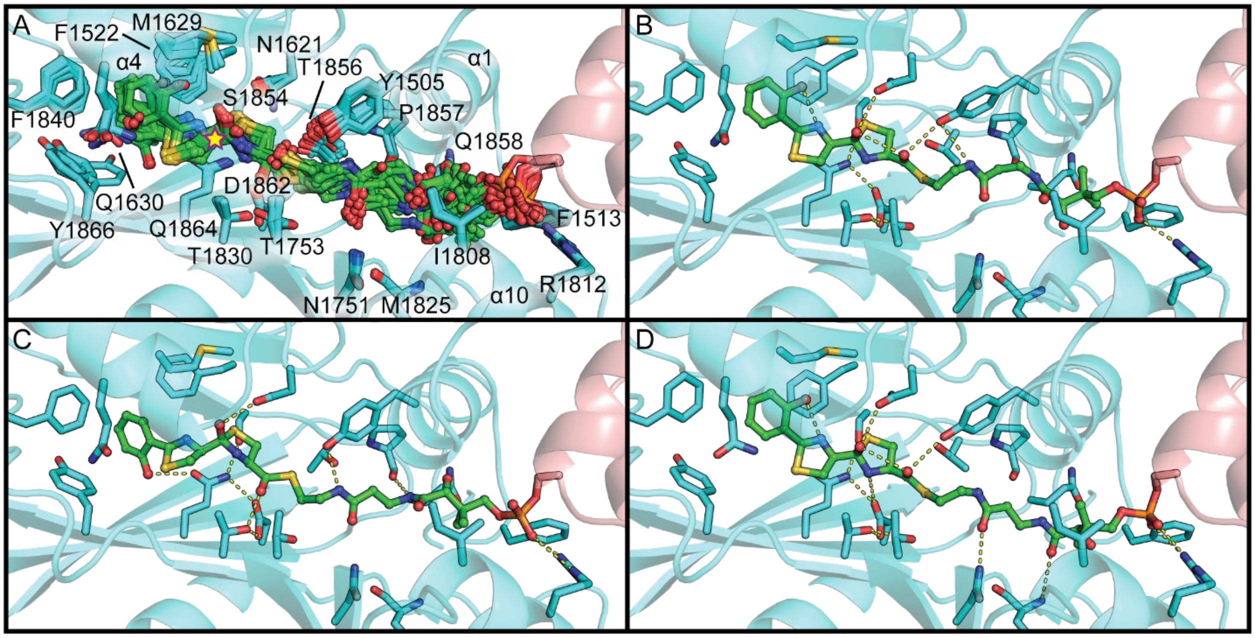


| | | | | | Prime MM-GBSA | | | | | |
|---|---|---|---|---|---|---|---|---|---|---|
| # | Panel | Dyad State | Glide Docking Score | Prime Energy kcal/mol | dG Bind kcal/mol | Complex Energy kcal/mol | Receptor Energy kcal/mol | Receptor Strain Energy kcal/mol | Ligand Energy kcal/mol | Ligand Strain Energy kcal/mol |
| 1 | B | neutral | -8.3 | -20949 | -62 | -21090 | -20918 | 3.5 | -109 | 19 |
| 2 | C | neutral | -11.2 | -20946 | -75 | -21084 | -20913 | 7.1 | -95.4 | 11 |
| 3 | D | neutral | -10.8 | -20943 | -63 | -21085 | -20916 | 5.9 | -106 | 17 |

**Figure S18 – Hydroxyl-toward-α4 Ppant-2HPTT(*S*)-OH poses from covalent docking.** HMWP2-Cy2 is in cyan, and Ppant-2HPTT-OH is in green. **A**, A superimposition of the class of poses in which the leaving group oxygen is directed toward the N-terminus of helix α4 is shown. A yellow star marks the leaving group oxygen. This class consistently positions the pantetheine so that Y1505 can interact with the thioester or the adjacent pantetheine amide. In these poses, Y1505 is too far from the reactive hydroxythiazolidine ring to form direct interactions with it. Both orientations of 2HPT are observed. **B-D**, Representative top poses are displayed in the same order as in the table at the bottom of the figure (sorted by Prime Energy of the refined covalent docking complex). The pantetheine dimethyl group is found in orientations resembling either AB3403/FscG (PDB IDs 4ZXI or 7KW0, respectively) or ObiF1 (PDB ID 6N8E). The pose in panel C places the hydroxyl to the side of the N-terminus of helix α4. The table displays docking scores and energy properties for each pose in panels B-D. Table coloration uses the same scale in Figs. S18-23 and serves to highlight more favorable poses. Darker green corresponds to favorable energy properties. Darker red corresponds to less favorable strain energy terms.

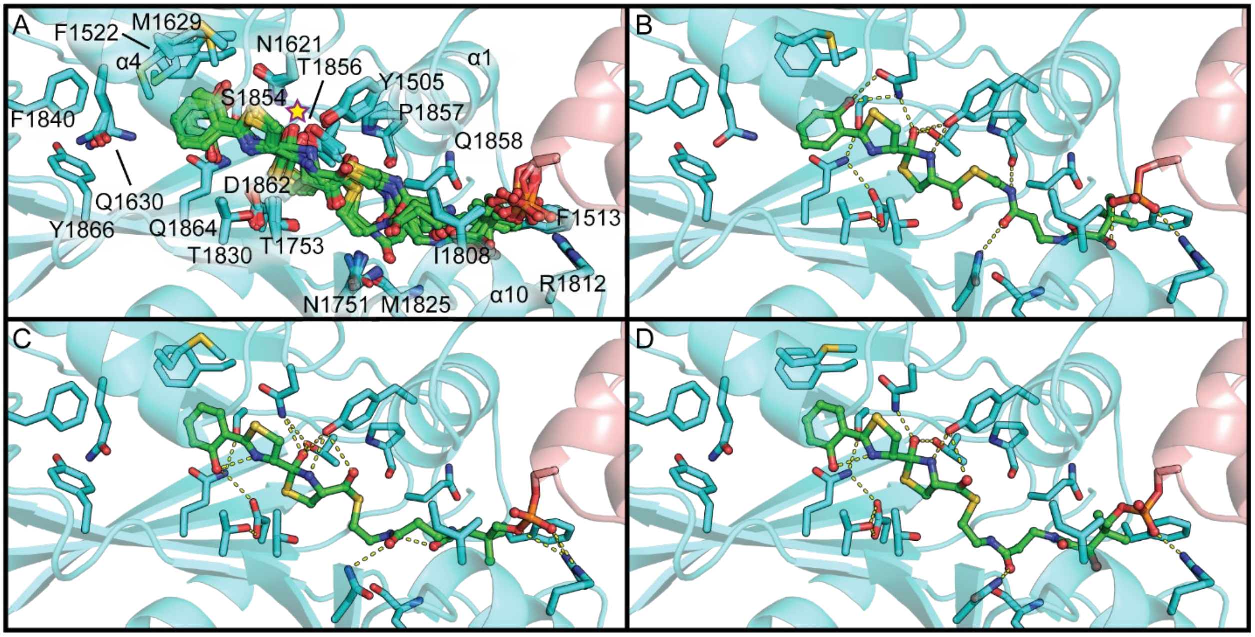


| # | Panel | Dyad State | Glide Docking Score | Prime Energy kcal/mol | Prime MM-GBSA dG Bind kcal/mol | Complex Energy kcal/mol | Receptor Energy kcal/mol | Receptor Strain Energy kcal/mol | Ligand Energy kcal/mol | Ligand Strain Energy kcal/mol |
|---|---|---|---|---|---|---|---|---|---|---|
| 4 | B | neutral | -10.2 | -20932 | -76 | -21072 | -20900 | 11 | -96.5 | 12 |
| 5 | C | neutral | -10.6 | -20924 | -60 | -21072 | -20914 | 11 | -101 | 20 |
| 6 | D | negative | -8.9 | -20917 | -68 | -21066 | -20896 | 12 | -83.7 | 26 |

**Figure S19 – Hydroxyl-toward-N-terminal-sheet Ppant-2HPTT(*S*)-OH poses from covalent docking.** HMWP2-Cy2 is in cyan, and Ppant-2HPTT-OH is in green. **A**, A superimposition of the class of poses in which the leaving group oxygen is directed toward the N-terminal subdomain β sheet is shown. A yellow star marks the leaving group oxygen. This class positions the pantetheine thioester so that Y1505 can interact with its carbonyl oxygen, but Y1505 also forms interactions with the hydroxythiazolidine ring nitrogen and the leaving group oxygen in some poses. Both orientations of 2HPT are observed. **B-D**, Representative top poses are displayed in the same order as in the table at the bottom of the figure (sorted by Prime Energy of the refined covalent docking complex). All these poses position the pantetheine dimethyl similarly to ObiF1 (PDB ID 6N8E). These poses all necessitate a flip of the leaving group oxygen from toward the N-terminus of helix α4 (where it would be expected to reside following the condensation reaction) to toward the N-terminal sheet, which would likely also necessitate substantial motion of the 2HPT side chain. The table displays docking scores and energy properties for each pose in panels B-D. Table coloration uses the same scale in Figs. S18-23 and serves to highlight more favorable poses. Darker green corresponds to favorable energy properties. Darker red corresponds to less favorable strain energy terms.

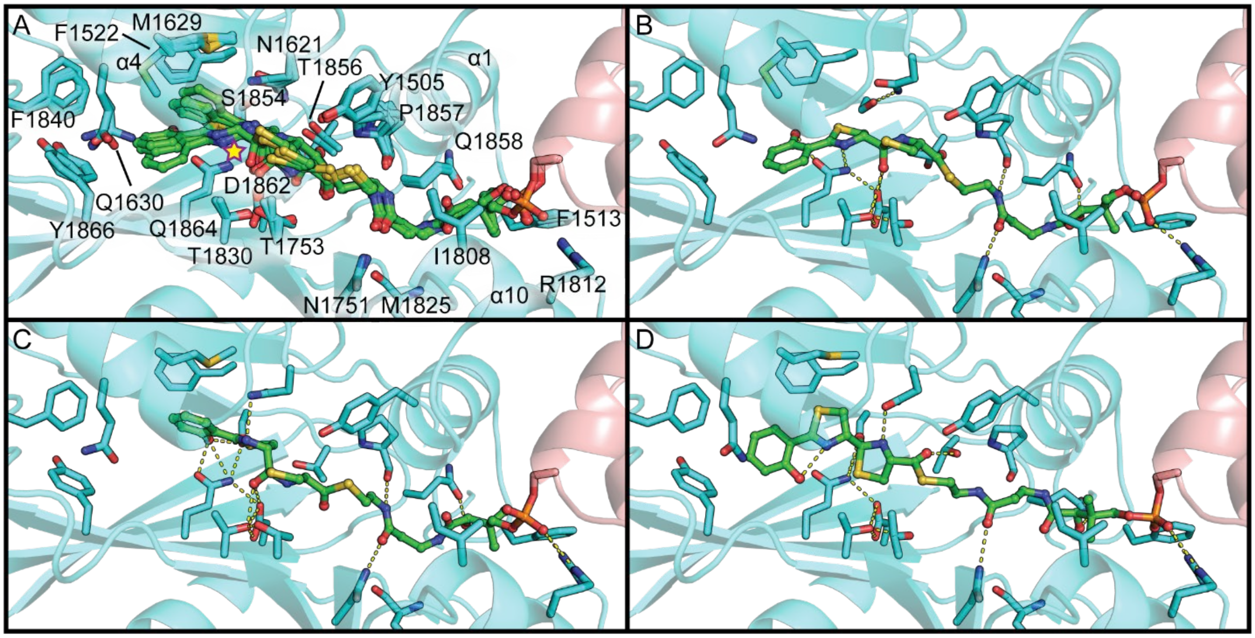


| # | Panel | Dyad State | Glide Docking Score | Prime Energy kcal/mol | Prime MM-GBSA dG Bind kcal/mol | Complex Energy kcal/mol | Receptor Energy kcal/mol | Receptor Strain Energy kcal/mol | Ligand Energy kcal/mol | Ligand Strain Energy kcal/mol |
|---|---|---|---|---|---|---|---|---|---|---|
| 7 | B | negative | -8.5 | -20935 | -79 | -21073 | -20896 | 5.4 | -98.1 | 18 |
| 8 | C | negative | -10.0 | -20906 | -55 | -21045 | -20895 | 8.2 | -94.9 | 9.1 |
| 9 | D | negative | -8.8 | -20925 | -73 | -21063 | -20887 | 5.0 | -92.2 | 12 |

**Figure S20 – Ppant-2HPTT(*S*)-OH covalent docking poses in hydroxyl-toward-dyad or deeply placed hydroxyl-toward-N-terminal-sheet orientations.** HMWP2-Cy2 is in cyan, and Ppant-2HPTT-OH is in green. **A**, A superimposition of the class of poses in which the leaving group oxygen is directed toward the putatively catalytic dyad (HMWP2 T1830-D1862) is shown. A yellow star marks the leaving group oxygen. This class positions the pantetheine thioester so that Y1505 could potentially interact with its sulfur. These poses consistently place the Ppant to allow hydrogen bonding with N1751 and typically also with the carbonyl of P1857. Only the 2HPT orientation with the phenol hydroxyl toward the thiazoline sulfur are observed. **B and C**, Representative top poses of the hydroxyl-toward-dyad orientation are displayed in the same order as in the table at the bottom of the figure (sorted by Prime Energy of the refined covalent docking complex). These poses position the Ppant dimethyl in states resembling either AB3403 (PDB ID 4ZXI) or ObiF (PDB ID 6N8E), and all necessitate a flip of the leaving group oxygen from toward the N-terminus of helix α4 (where it would be expected to reside following the condensation reaction) to toward the dyad, which would likely also necessitate substantial motion of the 2HPT side chain. **D**, This pose is an outlier of the hydroxyl-toward-N-terminal-sheet class placing the 2HPT side chain most deeply in the side chain-binding region and displays low divergence from 2HPT planarity in the orientation with the phenol hydroxyl toward the thiazoline nitrogen. The leaving group oxygen would have to undergo a large rotation from its hypothesized position during condensation to obtain this position during cyclodehydration, which would probably also necessitate similarly largescale motion of the 2HPT side chain. The table displays docking scores and energy properties for each pose in panels B-D. Table coloration uses the same scale in Figs. S18-23 and serves to highlight more favorable poses. Darker green corresponds to favorable energy properties. Darker red corresponds to less favorable strain energy terms.

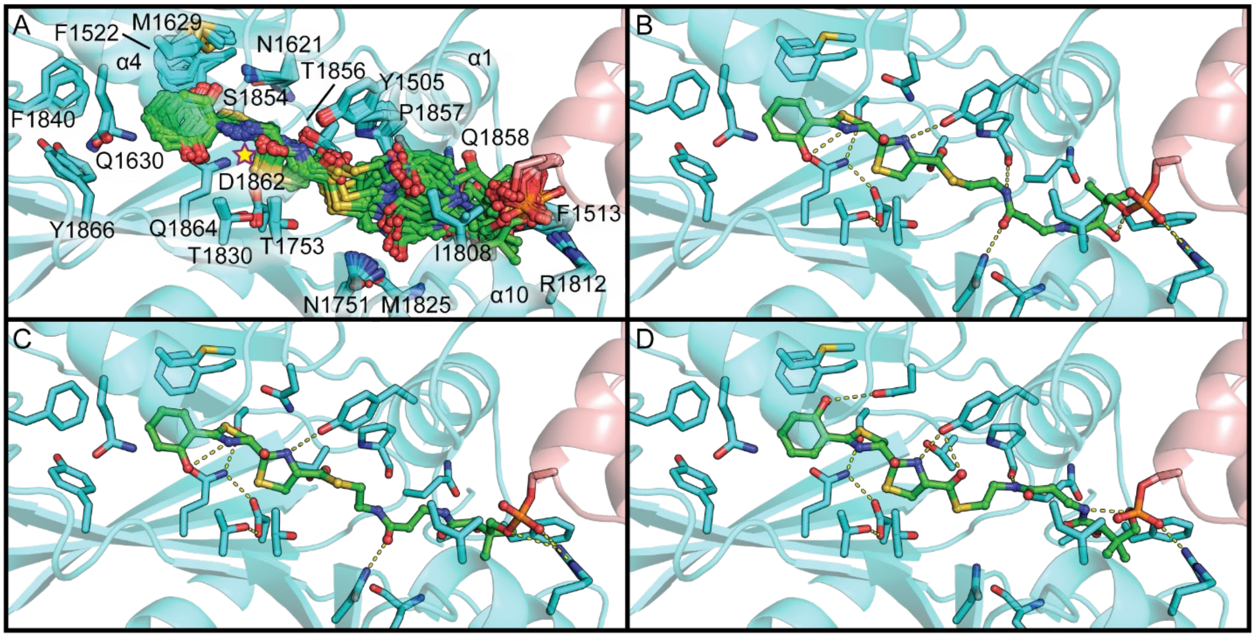


| # | Panel | Dyad State | Glide Docking Score | Prime Energy kcal/mol | Prime MM-GBSA: dG Bind kcal/mol | Prime MM-GBSA: Complex Energy kcal/mol | Prime MM-GBSA: Receptor Energy kcal/mol | Prime MM-GBSA: Receptor Strain Energy kcal/mol | Prime MM-GBSA: Ligand Energy kcal/mol | Prime MM-GBSA: Ligand Strain Energy kcal/mol |
|---|---|---|---|---|---|---|---|---|---|---|
| 10 | B | neutral | -10.4 | -20936 | -82 | -21083 | -20904 | 6.1 | -96.4 | 8.3 |
| 11 | C | neutral | -10.0 | -20935 | -80 | -21080 | -20903 | 5.1 | -96.4 | 9.1 |
| 12 | D | neutral | -9.2 | -20931 | -64 | -21069 | -20913 | 10 | -91.6 | 9.8 |

**Figure S21 – Hydroxyl-toward-α4 Ppant-2HPTT(*R*)-OH poses from covalent docking.** HMWP2-Cy2 is in cyan, and Ppant-2HPTT-OH is in green. **A**, A superposition of the class of poses in which the leaving group oxygen is directed toward helix α4 is shown. This class displays both 2HPT orientations and some degree of diversity at the pantetheine thioester linkage and the pantetheine amine position near the dimethyl group. Some poses place the dimethyl moiety of pantetheine toward I1808 in a manner similar to AB3403 (PDB ID 4zxi), but most poses place it in the opposite direction in a way that resembles ObiF1 (PDB ID 6N8E) or between the two, with the dimethyl directed toward the loop after helix α10. Additionally, the pantetheine amide nearer the thioester linkage appears to orient either so its N-H forms a hydrogen bond with the carbonyl of P1857 or so its carbonyl oxygen can act as hydrogen bond acceptor to N1751—sometimes also allowing hydrogen bonding between its N-H and the carbonyl of P1857. As observed in some of the Ppant-2HPTT(*S*)-OH poses, the hydroxyl of Y1505 is in the vicinity of the pantetheine thioester linkage and/or the adjacent pantetheine amide, suggesting its ability to act as hydrogen bond partner with these groups. **B-D**, Representative top poses are displayed in the same order as in the table at the bottom of the figure (sorted by Prime Energy of the refined covalent docking complex). Table coloration uses the same scale in Figs. S18-23 and serves to highlight more favorable poses. Darker green corresponds to favorable energy properties. Darker red corresponds to less favorable strain energy terms.

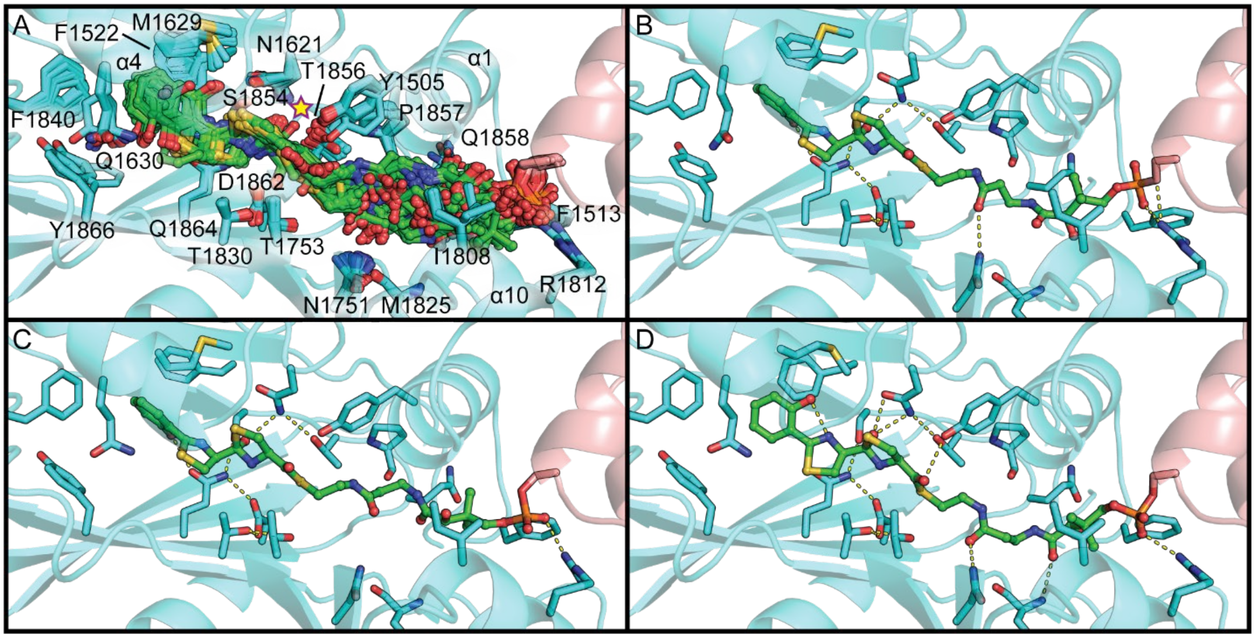


| # | Panel | Dyad State | Glide Docking Score | Prime Energy kcal/mol | Prime MM-GBSA dG Bind kcal/mol | Complex Energy kcal/mol | Receptor Energy kcal/mol | Receptor Strain Energy kcal/mol | Ligand Energy kcal/mol | Ligand Strain Energy kcal/mol |
|---|---|---|---|---|---|---|---|---|---|---|
| 13 | B | neutral | -9.9 | -20954 | -73 | -21098 | -20923 | 6.7 | -103 | 8.3 |
| 14 | C | neutral | -10.4 | -20951 | -74 | -21094 | -20919 | 3.8 | -102 | 6.4 |
| 15 | D | neutral | -9.5 | -20947 | -76 | -21096 | -20920 | 6.7 | -100 | 13 |

**Figure S22 – Hydroxyl-toward-N-terminal-sheet Ppant-2HPTT(*R*)-OH poses from covalent docking.** HMWP2-Cy2 is in cyan, and Ppant-2HPTT-OH is in green. **A**, A superposition of the class of poses in which the leaving group oxygen is directed toward the N-terminal β sheet is shown. This class displays a range of 2HPT and pantetheine conformations. Some poses place the dimethyl moiety of pantetheine toward I1808 in a manner similar to AB3403 (PDB ID 4ZXI), but most poses place it in the opposite direction in a way that resembles ObiF1 (PDB ID 6N8E) or between the positions, directed toward the loop after helix α10. Additionally, positioning of the pantetheine amides does not appear to particularly favor interactions with N1751 or P1857. Notably, however, Q1858 occupies several different rotameric states resembling EpoB-Cy/BmdB-Cy2 states. This allows for interactions between Q1858 and the pantetheine hydroxyl or the amide nearest to it. In these poses, the hydroxyl of Y1505 is in the vicinity of the carbonyl of the pantetheine thioester linkage. **B-D**, Representative top poses are displayed in the same order as in the table at the bottom of the figure (sorted by Prime Energy of the refined covalent docking complex). As noted for the other diastereomer's poses of this type, the leaving group oxygen would have to traverse a large distance from its expected position during the condensation reaction to obtain the positions observed in these poses. Table coloration uses the same scale in Figs. S18-23 and serves to highlight more favorable poses. Darker green corresponds to favorable energy properties. Darker red corresponds to less favorable strain energy terms.

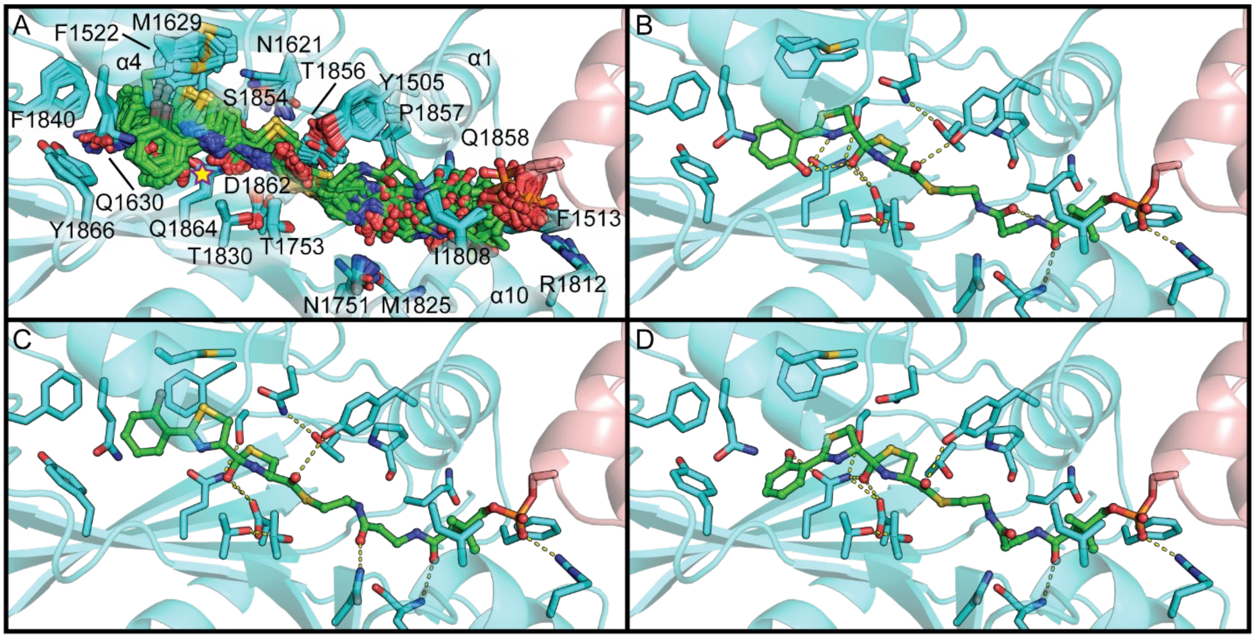


| # | Panel | Dyad State | Glide Docking Score | Prime Energy kcal/mol | Prime MM-GBSA: dG Bind kcal/mol | Complex Energy kcal/mol | Receptor Energy kcal/mol | Receptor Strain Energy kcal/mol | Ligand Energy kcal/mol | Ligand Strain Energy kcal/mol |
|---|---|---|---|---|---|---|---|---|---|---|
| 16 | B | neutral | -9.3 | -20960 | -72 | -21104 | -20927 | 8.1 | -105 | 19 |
| 17 | C | neutral | -9.4 | -20954 | -72 | -21096 | -20923 | 8.7 | -100 | 14 |
| 18 | D | neutral | -8.7 | -20945 | -77 | -21087 | -20913 | 2.5 | -96.2 | 17 |

**Figure S23 – Hydroxyl-toward-dyad Ppant-2HPTT(*R*)-OH poses from covalent docking.** HMWP2-Cy2 is in cyan, and Ppant-2HPTT-OH is in green. **A**, A superposition of the class of poses in which the leaving group oxygen is directed toward the putatively catalytic dyad is shown. This class displays both 2HPT orientations and consistently permits interaction between Y1505 and the carbonyl of the pantetheine thioester linkage. Unlike the other classes of poses, many of these poses, including the class representatives, place the dimethyl moiety of pantetheine toward I1808 in a manner similar to AB3403 (PDB ID 4zxi). Additionally, positioning of the pantetheine amides appears compatible with interactions with N1751 or the backbone N-H of M1825, and the amide nearest the pantetheine dimethyl moiety is consistently positioned in a way that is interesting when compared against alternative rotameric states of Q1858 observed in EpoB-Cy or BmdB-Cy2 (PDB IDs 5T7Z or 5T3E, respectively). **B-D**, Representative top poses are displayed in the same order as in the table at the bottom of the figure (sorted by Prime Energy of the refined covalent docking complex). As noted for the other diastereomer's poses of this type, the leaving group oxygen would have to traverse a large distance from its expected position during the condensation reaction to obtain the positions observed in these poses. Table coloration uses the same scale in Figs. S18-23 and serves to highlight more favorable poses. Darker green corresponds to favorable energy properties. Darker red corresponds to less favorable strain energy terms.

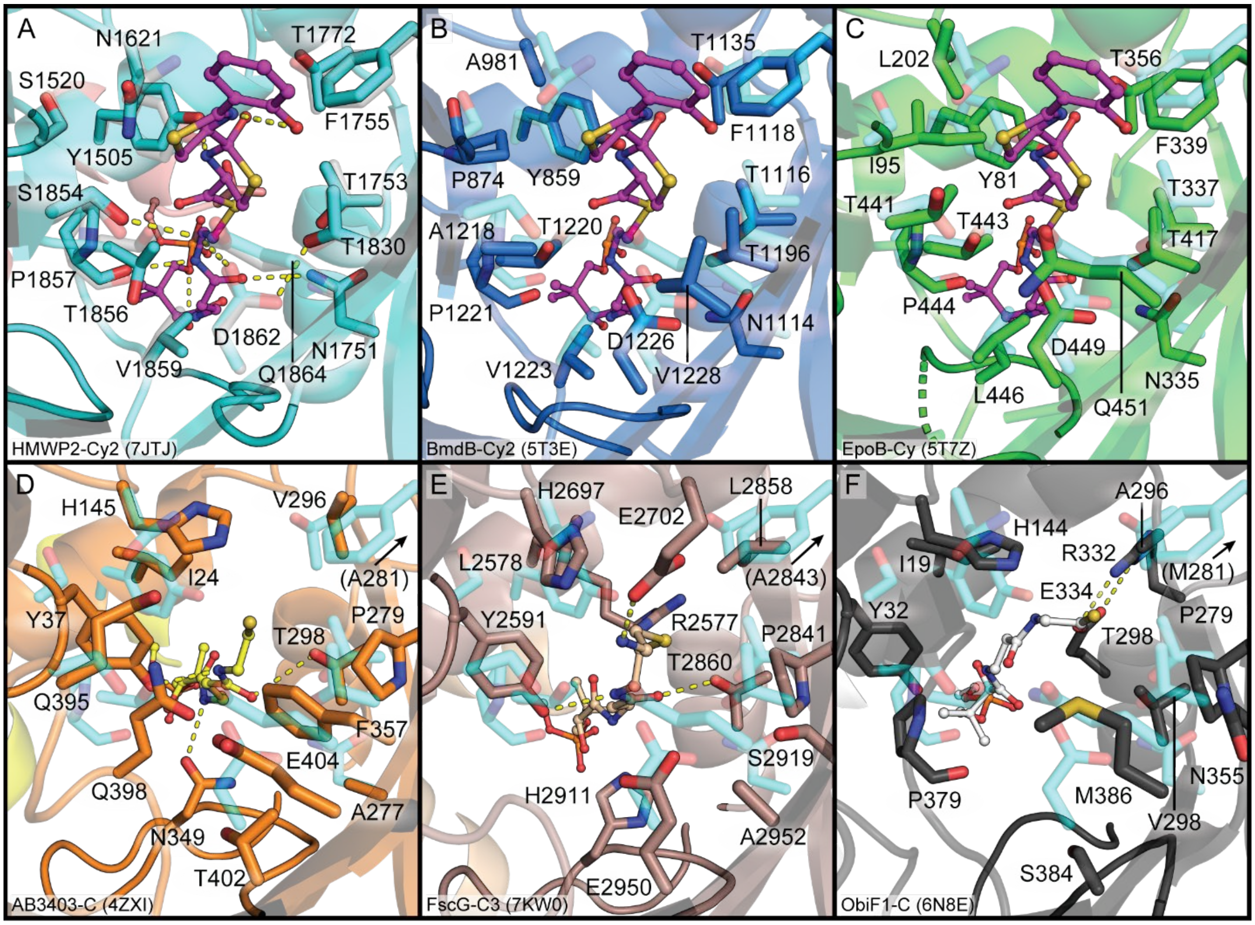


**Figure S24 – Comparison of possible pantetheine interactions in the downstream tunnels of Cy domains against observed C domain pantetheine interactions.** The system in each panel is listed in the bottom left corner of each panel along with the PDB ID. Every panel contains HMWP2-Cy2 crystallographic coordinates in transparent gray (panel A) or transparent cyan (panels B-F) sticks. Loops in opaque cartoon representation are those after helices α1 and α10. **A**, The top Ppant-2HPT(*R*)T-OH pose is in magenta ball and stick, and polar contacts/hydrogen bonds within the ligand or between ligand and receptor are shown as yellow dashed lines. In the foreground, putatively catalytic aspartate (D1862) is displayed in transparent sticks. **B and C**, Displayed are superimpositions of the HMWP2-Cy2 crystallographic active site coordinates with BmdB-Cy2 or EpoB-Cy2, respectively, with the top Ppant-2HPT(*R*)T-OH pose in magenta ball and stick. The missing residue in the foreground of panel C is an arginine that is not conserved in the Cy family but may act with the conserved arginine of the downstream tunnel entrance in EpoB to bind the phosphopantetheine phosphate. **D-F**, Displayed are superimpositions of the HMWP2-Cy2 crystallographic active site coordinates with AB3403-C, FscG-C3 or ObiF1-C, respectively. Each system's ligand is shown in ball and stick with carbon atoms colored corresponding to the PCP for that system. In panel D, three hydrogen bonding opportunities between pantetheine and protein are observed involving Y37, N349 and T298. In panel E, only two hydrogen bonds are identified (Y2591/T2860), but H2911 is also positioned to interact with a pantetheine amide. Most notably in this structure, E2702 forms a salt bridge with the ammonium group of the stabilized Ppant-glycine substrate mimic, but it is also interesting to note that R2577 extends from helix α1 in that system to form weak interactions with the pantetheine carbonyls. In panel F, no explicit polar contacts between protein and pantetheine are observed, and it may be that this conformation of pantetheine does not resemble a condensation-compatible state and would adopt different interactions in the reaction complex, perhaps involving R332 in a role similar to R2577 in panel E.

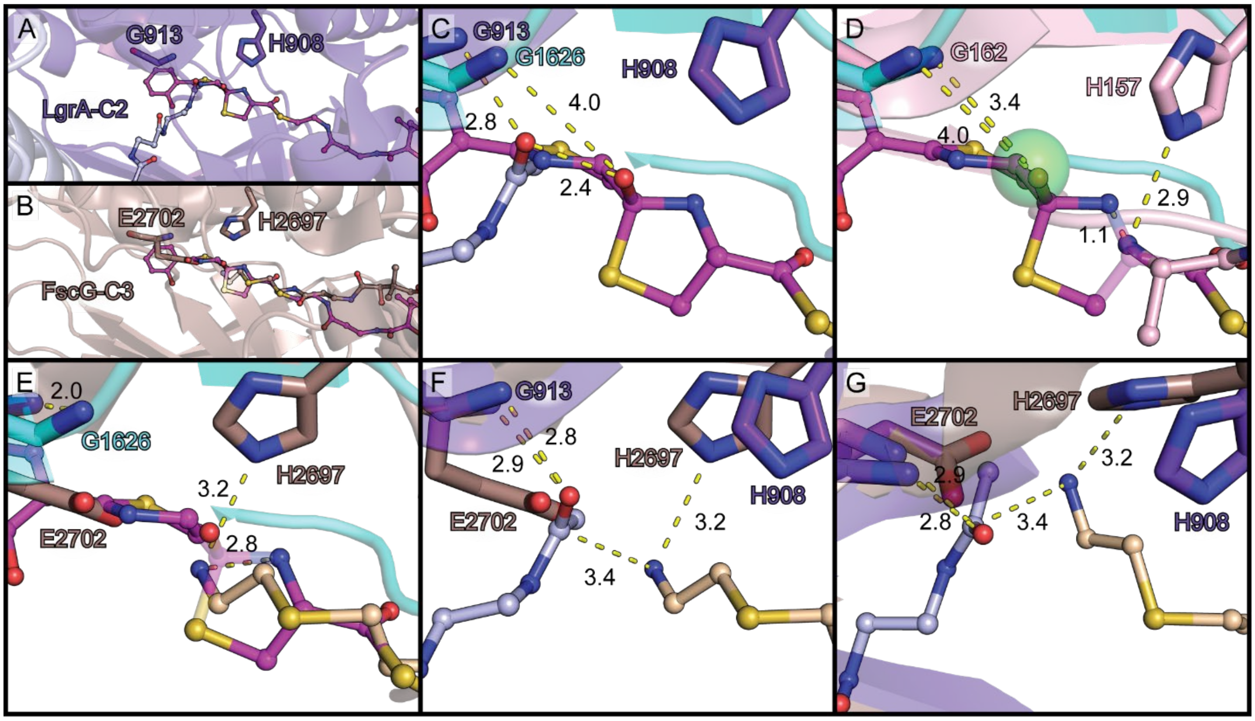


**Figure S25 – Geometric rationale for a proposed pre-condensation state shared by C and Cy domains. A and B**, These panels provide an overview of the active site regions of LgrA-C2 and FscG-C3, respectively. Coloration in these panels is continued through C, E, F and G. G913 and E2702 reside at the N-terminus of helix α4, and H908 is the conserved C domain histidine that may act as a base or to stabilize the nucleophile or zwitterionic, tetrahedral condensation transition state. The ball and stick representations are the ligands bound in LgrA-C2 (light blue) or FscG-C3 (wheat) superimposed with the top Ppant-2HPT(*R*)T-OH hydroxyl-toward-α4 pose (magenta). **C**, Displayed is the superimposition of the top Ppant-2HPT(*R*)T-OH hydroxyl-toward-α4 pose (magenta ball and stick and cyan protein) with LgrA-C2. The helix α4 N-terminal residue's backbone N-H in both systems is directed toward the leaving group oxygen. There is an approximately 1.2 Å increase in the distance between these groups going from HMWP2-Cy2 to LgrA-C2, but 2.4 Å separate the leaving group oxygen atoms in these models. **D**, Displayed is the superimposition of the top Ppant-2HPT(*R*)T-OH hydroxyl-toward-α4 pose (magenta ball and stick and cyan protein) with an acceptor substrate mimic-bound state of CDA-C1 (PDB ID 5DUA chain B). This substrate mimic was found to be competent to undergo the condensation reaction to yield a tethered condensation product. Interestingly, in this active site at the N-terminus of helix α4, there is a chloride ion (green transparent sphere) that corresponds rather closely to the leaving group oxygen of the Ppant-2HPT(*R*)T-OH pose (~0.5 Å between centers). This superposition places the hydroxythiazolidine of the cyclized intermediate between analogous groups in the CDA-C1 model, with the nucleophilic nitrogen 1.1 Å from the hydroxythiazolidine nitrogen. Superposition of this CDA-C1 model with the LgrA-C2 model (not shown here) results in a slightly greater nucleophile-to-electrophile distance than that implied by the superimposition of LgrA-C2 and FscG-C3 in panels F and G. **E**, Displayed is the superimposition of the top Ppant-2HPT(*R*)T-OH hydroxyl-toward-α4 pose (magenta ball and stick and cyan protein) with FscG-C3 including a stabilized Ppant-glycine substrate mimic (wheat). The tetrahedral center bearing the leaving group oxygen of the cyclodehydration intermediate is shown in transparency so that the nucleophile position in FscG-C3 and the measurement between nitrogen atoms are visible. Notably, by interacting with E2702 the acceptor substrate mimic's terminal nitrogen is drawn closer to the N-terminus of helix α4 than in the CDA-C1 or HMWP2-Cy2 models. The position of E2702 is not commonly a large polar or charged residue, and the authors reporting the FscG-C3 structure note that it likely plays a role in positioning very small acceptor substrates such as glycinyl Ppant. **F and G**, Shown are views of the superposition of LgrA-C2 and FscG-C3 models related by a 90° rotation around the X axis. Most importantly, FscG E2702 occludes the path of the donor mimic in LgrA-C2. Additionally, the rotameric states of the conserved histidine residues in these models are rather different, demonstrating the ability of a residue at this position to rotate through a wide angle without largescale differences in surrounding residues. Although the nucleophile is relatively close to the electrophile in this superimposition, and their relative orientations form a favorable Bürgi-Dunitz angle (>90°), the nucleophile is offset from direct attack by a Flippin-Lodge angle around 45°.

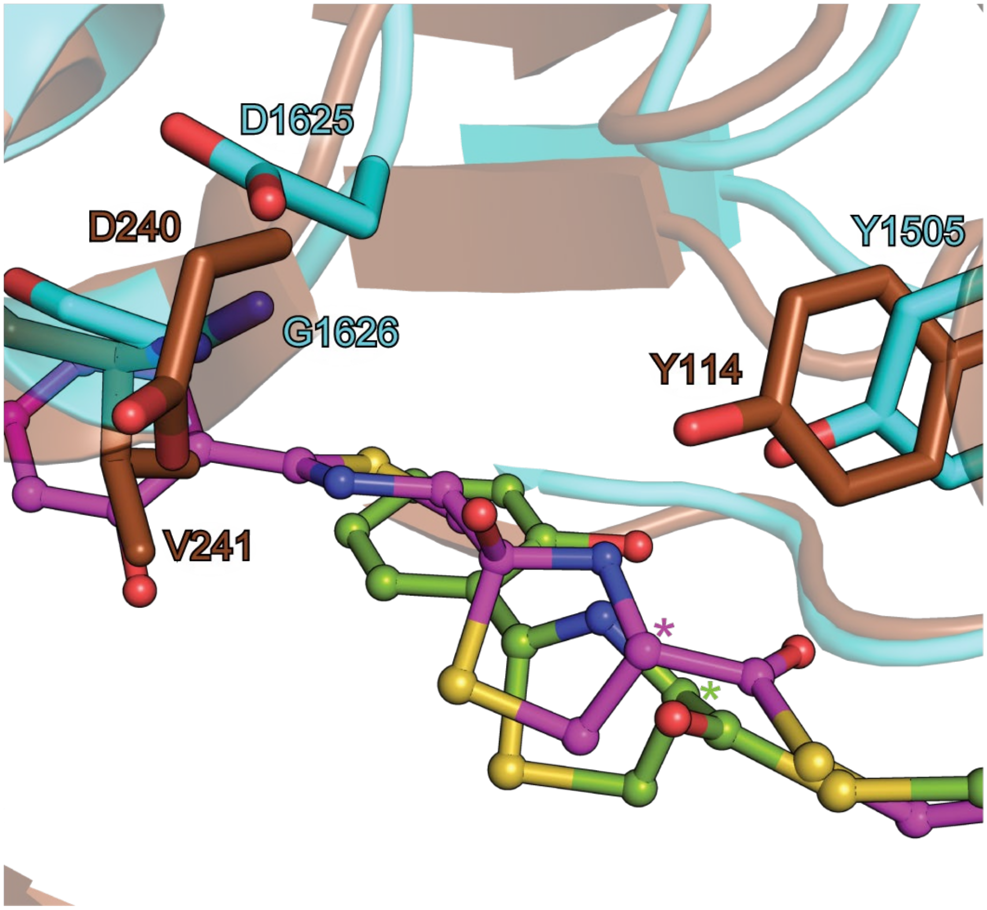


**Figure S26 – Comparison of the top cyclodehydration intermediate pose from HMWP2-Cy2 and the product-bound state reported for PchE-Cy.** The vicinities of the active sites differ between PchE-Cy (PDB ID 7EN1) and HMWP2-Cy2; however, the cyclodehydration intermediate docking model for HMWP2-Cy2 (magenta) is placed fairly similarly to the product from PchE-Cy. Asterisks mark the Cα atoms of the acceptor cysteines from both models, which display opposite chirality, with the L-cysteine thought to be accepted by HMWP2-Cy2 and D-cysteine reported by Wang et al. (Wang et al., 2022).

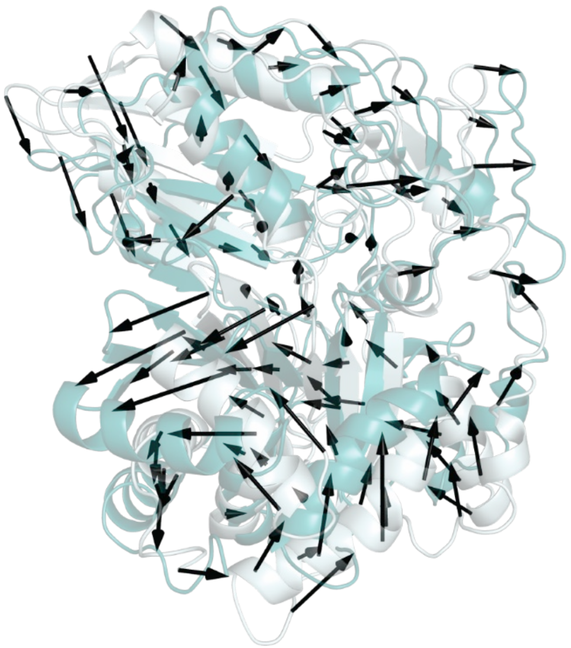

**Figure S27 – The high-amplitude HMWP2-Cy2 low-frequency normal mode number 7.** Mode 7 of HMWP2-Cy2 exhibits the greatest relative amplitude among the 10 lowest frequency modes. Here the extrema coordinates of the mode trajectory are displayed as pale cyan and deep teal, with distances between equivalent Cα atoms undergoing substantial displacement marked by black arrows. The left-most lobe of the C-terminal subdomain, comprised of strands 10, 12, 13 and 7, makes large movements away from the floor loop (larger vectors pointing toward the left side of the figure). With this motion, the tip of the N-terminal subdomain swings in toward the C-terminal subdomain. Consistent with comparisons against C domains in condensation donor-like states, part of the floor loop can also be seen to move away from the upstream tunnel entrance, toward the PCP1 linker and the hinge region between helices α5 and α6. (See also the movies of normal mode animations in the electronic supporting information.)

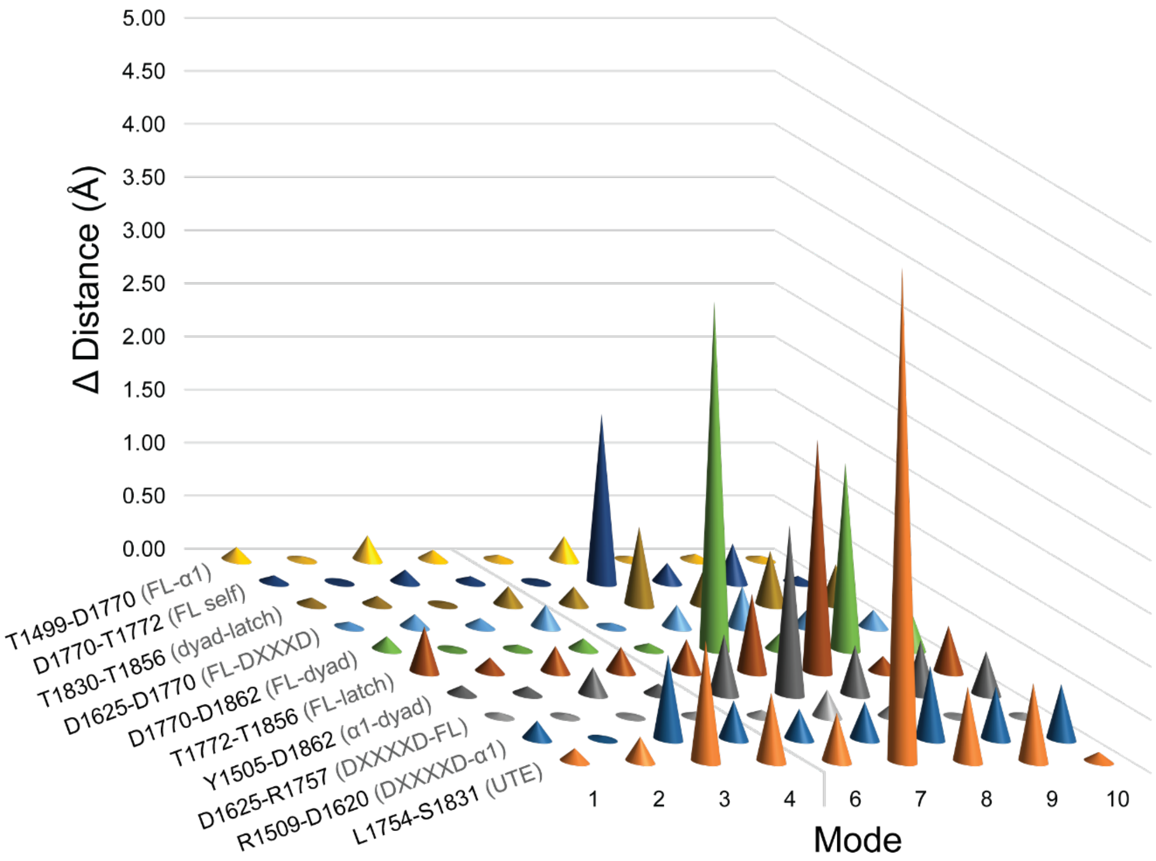


**Figure S28 – Change in inter-residue distances with the top 10 low-frequency normal vibrational modes of HMWP2-Cy2.** A model of HMWP2-Cy2 returns a variety of motions in the normal mode approximation, primarily involving changes of orientation between the N- and C-terminal subdomains. Marked in the rows of this plot are a number of inter-residue distances relevant for considerations of active site shape and/or the structure of PCP/Ppant-binding regions. Of most interest for reaching a condensation-like state is the distance L1754-S1831, which spans the gap between the C-termini of strands β8 and β10. The highest amplitude excursion for this gap is found in mode 7, which also influences the floor loop aspartate to active site aspartate distance (D1770-D1862), the floor loop aspartate to floor loop threonine distance (D1770-T1772) and the α1 (Y1505) to active site aspartate distance (Y1505-D1862). The width of the active site in the direction roughly orthogonal to the helix α4 to active site aspartate distance varies with mode 8, and mode 9 influences the floor loop/active site relationship. Of the lowest frequency modes, mode 1 also influences active site width, and mode 3 weakly influences upstream tunnel entrance width (L1754-S1831) and the distance between the conserved α1/loop1 arginine (R1509) and the first aspartate of the DXXXXD motif (D1620, see Fig. S10C for an overview of these residues in the global context of the downstream tunnel). The light gray line separating modes 4 and 6 indicates omission of mode 5, which involved only side chain motion of R1817 and did not influence the measurements presented here.

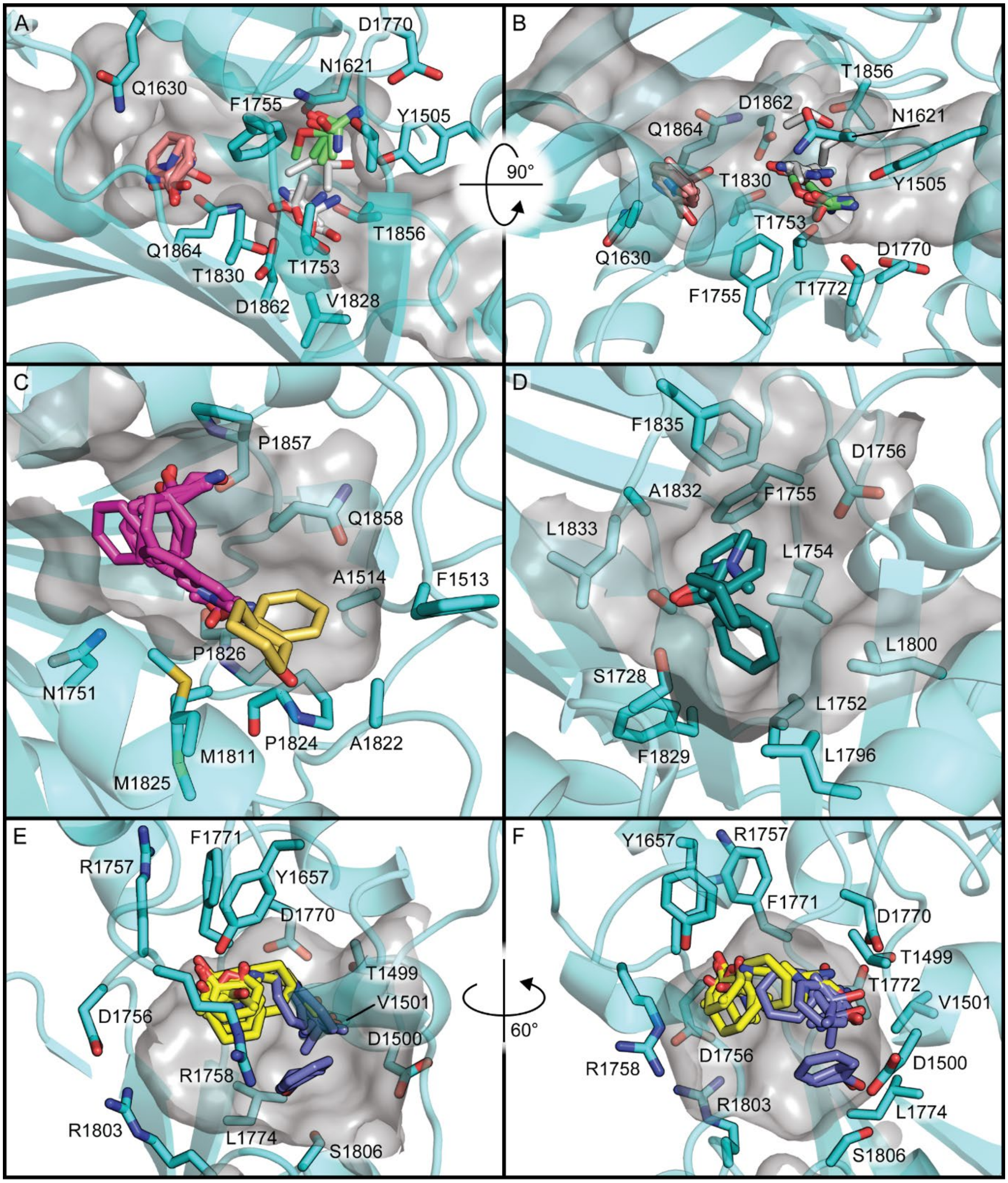


**Figure S29 – FTmap server results for HMWP2-Cy2. A and B**, Two views of small ligand clusters in the HMWP2-Cy2 active site are shown related by a 90° rotation around the X axis. Overall, binding of these fragments lends support to preferential positioning of the cyclodehydration intermediate in accordance with our proposal based on covalent docking. **C**, Two clusters of small organic ligands in the downstream tunnel are shown. These positions do not offer much in the way of specific interactions on which to build. **D,** Binding of a cluster of small organic ligands at the upstream tunnel entrance relies largely on hydrophobic packing, but specific polar contacts could also occur with S1728, which marks the expected phosphopantetheine phosphate binding site at the end of helix α8. **E and F**, Two clusters of small organic ligands bind in a cavity between the floor loop, hinge, and helices α1 and α9. Although some polar contacts are observed between protein and members of this cluster, docking at this site is probably largely driven by burying hydrophobic surface area. It is interesting to note that this pocket is in the region identified by Mishra et al. 2021 as a dynamical electrostatic environment coupled to conformational states of HMWP2-Cy1 (investigated in that work) as well as the larger C domain family.

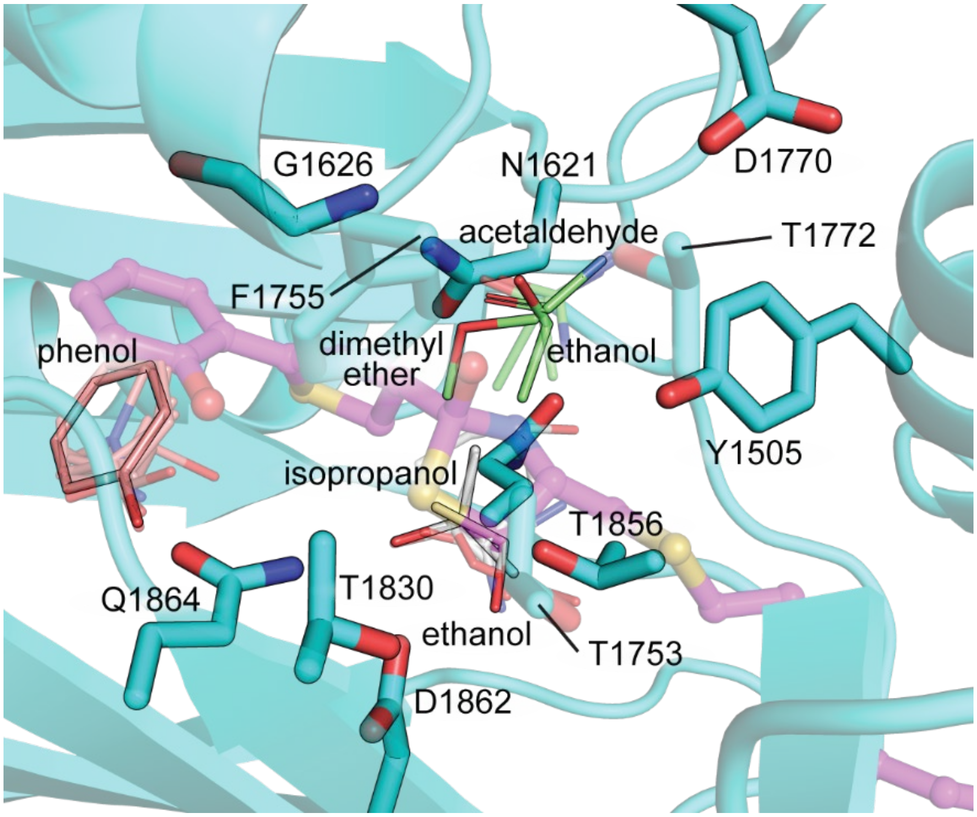


**Figure S30 – The HMWP2-Cy2 active site binds small organic molecules in FTmap docking.** Three clusters are displayed in salmon, white or light green thin sticks. Members of these clusters that are interesting in consideration of moieties of the cognate substrates are opaque, whereas other members of each cluster are transparent. Notably, hydroxyls of ethanol and isopropanol dock preferentially near the putatively catalytic dyad, oxygen atoms of ethanol and dimethyl ether are positioned at the N-terminus of helix α4 (G1626), loosely resembling crystallographic water sites, and phenol binds somewhat similarly to the hydroxyphenyl of Ppant-2HPT(*R*)T-OH in our top docking model (magenta ball and stick). Phenol is deeper into the side chain binding region than 2HPT of the docked pose, suggesting 2HPT may not be fully accessing the basin of attraction occupied by phenol in this volume. HMWP2-Cy2 residues from the floor loop (in the foreground) are displayed with transparency and are labeled with lines pointing to their Cα atoms.

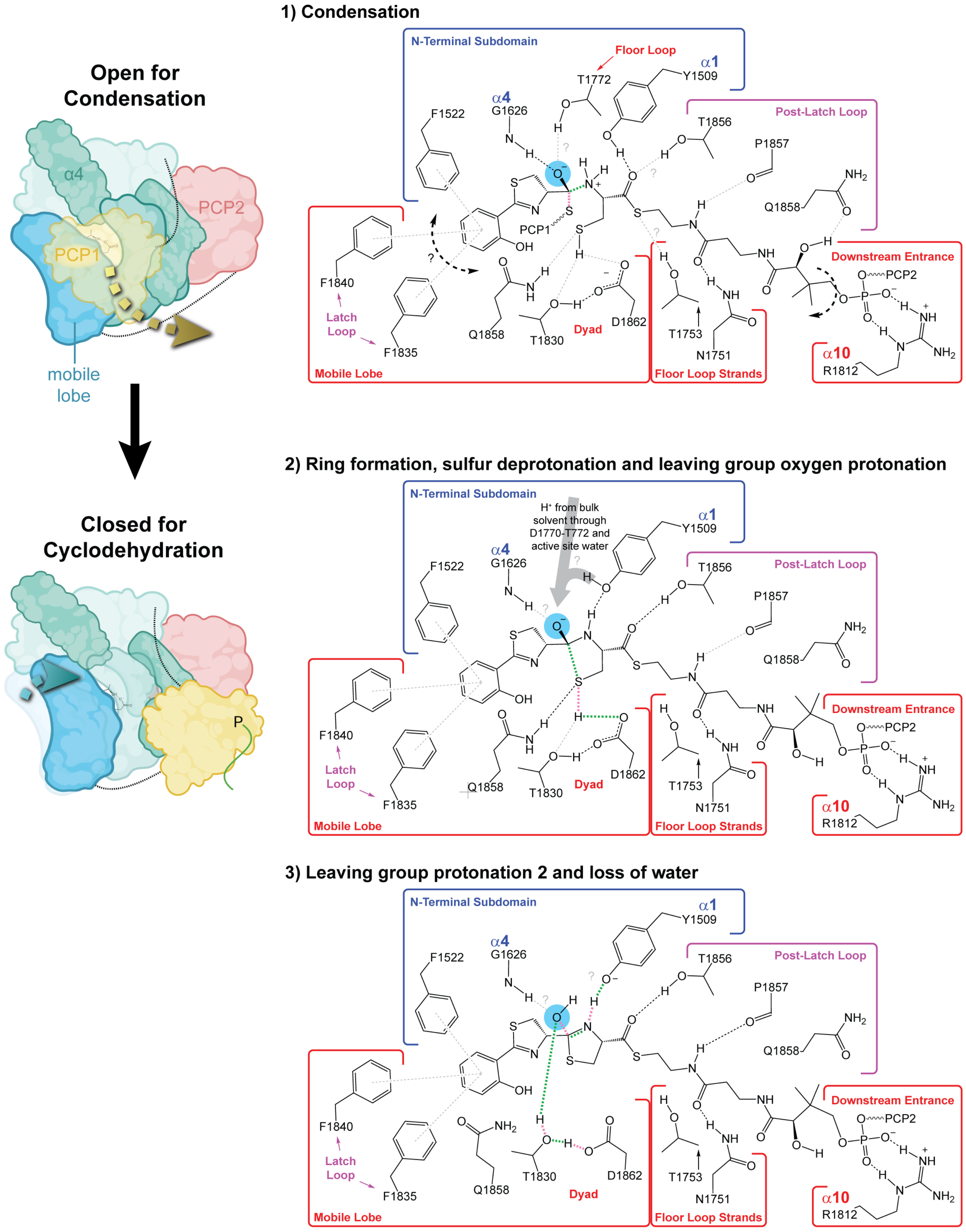
Open for Condensation
α4
PCP2
PCP1
mobile lobe
Closed for Cyclodehydration
P
1) Condensation
N-Terminal Subdomain
Floor Loop
α1
Y1509
T1772
α4
G1626
F1522
Post-Latch Loop
T1856
P1857
Q1858
NH2
PCP1
OH
Mobile Lobe
F1840
Latch Loop
F1835
Q1858
T1830
Dyad
D1862
Floor Loop Strands
T1753
N1751
Downstream Entrance
PCP2
α10
R1812
2) Ring formation, sulfur deprotonation and leaving group oxygen protonation
H⁺ from bulk solvent through D1770-T772 and active site water
3) Leaving group protonation 2 and loss of water

**Figure S31 – Proposals for a global Cy domain conformational change associated with catalysis and three main reaction steps.** On the left is a schematic representation of the conformational changes involved in transitioning from the open-for-condensation state to the closed-for-cyclodehydration state. On the right are proposals for active site organization during three main reaction steps. Green and pink dashed lines indicate bonds formed or broken in each step, respectively. Light gray dashed lines indicate interactions suggested by our model but for which there is less confidence given variation in our docking models, including omission of interacting water molecules. The first step, condensation, occurs in the open-for-condensation state of the Cy domain. Gray question marks in step 1 point out potential interactions between residues around the active site and the Ppant thioester or oxyanion. In the state depicted in step 1, it appears possible that the donor side chain could form a range of interactions with hydrophobic residues around the side chain-binding region (symbolized by the bold, black, dashed arrow and question mark on the left). After condensation, a Ppant conformational change characterized primarily by rotation of the dimethyl group by ~180° may be concomitant with the open-to-closed transition (bold, black, dashed arrow on the right), following displacement of the donor Ppant and PCP. The second step, in which ring closure occurs, may be partially synchronous with the open-to-closed transition, perhaps bringing T1753 within hydrogen bonding distance of T1830, as observed in the Cy domain crystal structures, thereby altering the potential of T1753 to interact with the Ppant thioester as suggested in step 1. The final step, including loss of a water, is envisioned as happening entirely in the closed-for-cyclodehydration state. Gray question marks in steps 2 and 3 point out that it is unclear whether the N-terminus of α4 interacts directly with the leaving group in those steps and that it is unknown whether Y1505 acts as a proton donor in step two, which would allow it to act as a base (phenolate form) in step 3 to deprotonate the ring nitrogen. The roles of active site waters—perhaps in sites like those observed crystallographically (Fig. 4)—are unknown, but their connection to the rather conserved residues Y1505, D1770 and T1772 suggests they could act to shuttle protons into or out of the active site during turnover. Additionally, release of the water leaving group into the environment of those waters could contribute to a global conformational change for product release.
